# The genomic basis of evolutionary stasis in the 500-million-year-old red seaweed genus *Ahnfeltia*

**DOI:** 10.64898/2026.05.26.727817

**Authors:** Hocheol Kim, Rachel Dobson, Seok-Wan Choi, Jihoon Jo, Chung Hyun Cho, Louis Graf, Danilo E. Bustamante, Martha S. Calderon, Andres Mansilla, Gayle I. Hansen, Anna V. Skriptsova, Allyson Nardelli, Kathy Ann Miller, Brenda Konar, Jong-Hwan Kwak, Choongwon Jeong, Daehan Lee, Alex Farnsworth, Paul Valdes, Daniel J. Lunt, Dongseok Kim, Yongsung Lee, Duckhyun Lhee, Rob Mrowicki, Juliet Brodie, Shuhai Xiao, Gary W. Saunders, Christine A. Maggs, Erin Saupe, Cock van Oosterhout, Thomas Mock, Hwan Su Yoon

## Abstract

The red seaweed genus *Ahnfeltia* is an ancient lineage that has persisted for over 500 million years with remarkably limited diversification despite a global distribution in cold-temperate intertidal habitats. Compared to the highly diverse sister lineage, Rhodymeniophycidae, *Ahnfeltia* provides a unique system for investigating long-term evolutionary persistence in marine macroalgae. Here, we generated chromosome-scale genomes from five populations across three species and combined population genomics with paleogeographic niche modelling. Our results reveal remarkable genomic conservation, strong geographic isolation with limited gene flow, high homozygosity, and evidence of purifying selection. Niche projections indicate long-term stability and spatial connectivity of suitable cold-temperate habitats. These findings suggest that *Ahnfeltia*’s persistence and limited diversification are linked to genomic constraints and stable ecological niches over geological timescales. This study provides new insights into the genomic basis of evolutionary stasis in ancient marine lineages and highlights potential vulnerability to ongoing climate change affecting cold-water coastal ecosystems.

## Introduction

Over geological time, many organisms have diversified, adapted, or disappeared as climates shifted, continents moved, and habitats were reorganized. Yet some lineages appear to have persisted through these changes with remarkably little diversification or phenotypic change. Understanding how such lineages survive over long evolutionary timescales without generating substantial diversity remains an important question in evolutionary biology (*1*). This pattern is particularly evident in so-called “living fossil” lineages, which typically exhibit ancient origins, long generation times, low species diversity, and limited phenotypic change over time (*2, 3*). Although several explanations have been proposed to account for this persistence, including slow mutation rates and duplicated gene families, particularly involved in DNA repair, the mechanisms underlying long-term evolutionary stasis remain debated (*2, 4, 5*).

Red seaweeds, like their green and brown counterparts, have diversified largely through dispersal and adaptation to environmental change throughout Earth’s history (*6–8*). Within the five subclasses of the class Florideophyceae, Ahnfeltiophycidae diverged from their last common ancestor about 508 million years ago (Ma) (*9*). Unlike its sister groups, Corallinophycidae (1,147 spp.) and Rhodymeniophycidae (5,298 spp.), Ahnfeltiophycidae includes only two genera (*Ahnfeltia* & *Pihiella*) and 11 species in total (*10, 11*), of which only one belongs to filamentous *Pihiella*. Although ten species have been reported within *Ahnfeltia*, comprehensive taxonomic studies have confirmed only three species (*i.e. Ahnfeltia plicata, Ahnfeltia borealis,* and *Ahnfeltia fastigiata*) with robust genetic evidence (*12–14*).

Species in the genus *Ahnfeltia* E.M. Fries have a dark red or brown phenotype composed of thin cylindrical branches (*15, 16*). This genus is known for producing high-quality agar with a low sulfate ratio (*17–19*). *Ahnfeltia* species usually grow in the intertidal and upper subtidal zone of rocky shores (including sandy-covered rocks) at higher latitudes where the annual average of seawater temperatures is below 14 °C (*12, 14, 16*). They are long-lived, with life spans commonly reaching three years and occasionally extending up to 5-10 years (*16, 20*). Although *Ahnfeltia* usually reproduce sexually, asexual reproduction has been reported under certain environmental conditions (*21, 22*). For example, vegetative propagation via thallus fragmentation has been observed in unattached populations of *A. fastigiata* at depths greater than 3 m in Peter the Great Bay and Sakhalin Island, where extensive mats may form that potentially consist of genetically identical clones (*23*). Growth rates are relatively low; apical elongation in *A. plicata* has been reported to remain below 20 µm day□¹ under cold-water conditions (*16, 24*). They are resilient to environmental stress, tolerating desiccation, low and even freezing temperatures, and strongly reduced solar irradiance during winter (*16, 25*).

*Ahnfeltia* species are largely restricted to higher latitudes (≥ 40°N/S) and exhibit geographic isolation among populations, which may contribute to the genetic differentiation observed between populations (*13*). However, species richness and overall genetic diversity are markedly lower than expected for a lineage that diverged over 500 million years ago and occupies geographically isolated habitats across multiple oceanic regions. Together, these traits raise the question of why diversification appears to be limited in this ancient lineage despite a longstanding evolutionary history and a global distribution of *Ahnfeltia* species and their populations.

To address this conundrum, we generated chromosomal level genomes from five populations of three *Ahnfeltia* species and re-sequenced individuals from geographically distinct populations. The lack of fossils for *Ahnfeltia* meant that our evolutionary reconstructions had to be inferred by the combination of different *in-silico* approaches. We applied a combination of comparative genomics, population genetics, and palaeogeographic niche modelling to examine patterns of genomic variation and historical suitability for the species. Our results suggest that *Ahnfeltia* has persisted with limited diversification despite long-term geographic isolation, likely associated with low genetic variation and stable ecological niches. Together, this study provides a genomic perspective on the long-term persistence of ancient marine lineages such as *Ahnfeltia* in physically stressful coastal environments.

## Results and Discussion

### Chromosomal genome assembly and global distribution of *Ahnfeltia*

The genus *Ahnfeltia* has been studied since the late 1970s because of its ecological relevance to rocky shore habitats and its considerable commercial value for agar products (*17, 21, 22, 26*). Historical assessments of its diversity and distribution have relied largely on morphologies and molecular markers such as 18S rDNA, *rbc*L and COI-5P. Several phylogenetic studies indicate that there are likely only three *Ahnfeltia* species - *A. plicata, A. borealis*, and *A. fastigiata*. This represents remarkably low species diversity for such an old lineage with a broad modern geographic distribution (table S1 and S2; *12, 14*).

Based on this framework, we generated high-quality whole-genome assemblies using PacBio sequencing for representatives of each species. For *A. plicata* and *A. fastigiata*, two isolates from different continents were selected, whereas for *A. borealis,* fresh material could be obtained only from Sakhalin Island. In total, five chromosome-level reference genomes were generated. To further assess the genomic diversity of *Ahnfeltia* populations on a global level, we have re-sequenced 102 individuals from 10 populations (Fig. 1, table S3). They are distributed exclusively in coastal oceans on rocky shores of higher latitudes on both hemispheres (ca. ≥ 40°), which is consistent with the distributional patterns reported in previous global phylogenetic studies of diverse *Ahnfeltia* populations (*12, 14*). The three recognized species have largely distinct distribution, although partial geographic overlap occurs in Northern Pacific and Arctic areas. *A. plicata* occurs in the North Atlantic, Tasmania, and Chile; *A. fastigiata* inhabits the North Pacific coasts; and *A. borealis* is found in the North American Arctic and Sakhalin Island in Russia. Overall, *Ahnfeltia* species share remarkably conserved phenotypes with wiry fronds branching from a discoid holdfast. This phenotypic conservation is consistent with long-term stabilizing selection in a persistent ecological niche, a pattern that may have constrained morphological divergence and contributed to the lineage’s low species diversity (Fig. 1).

**Fig. 1.**
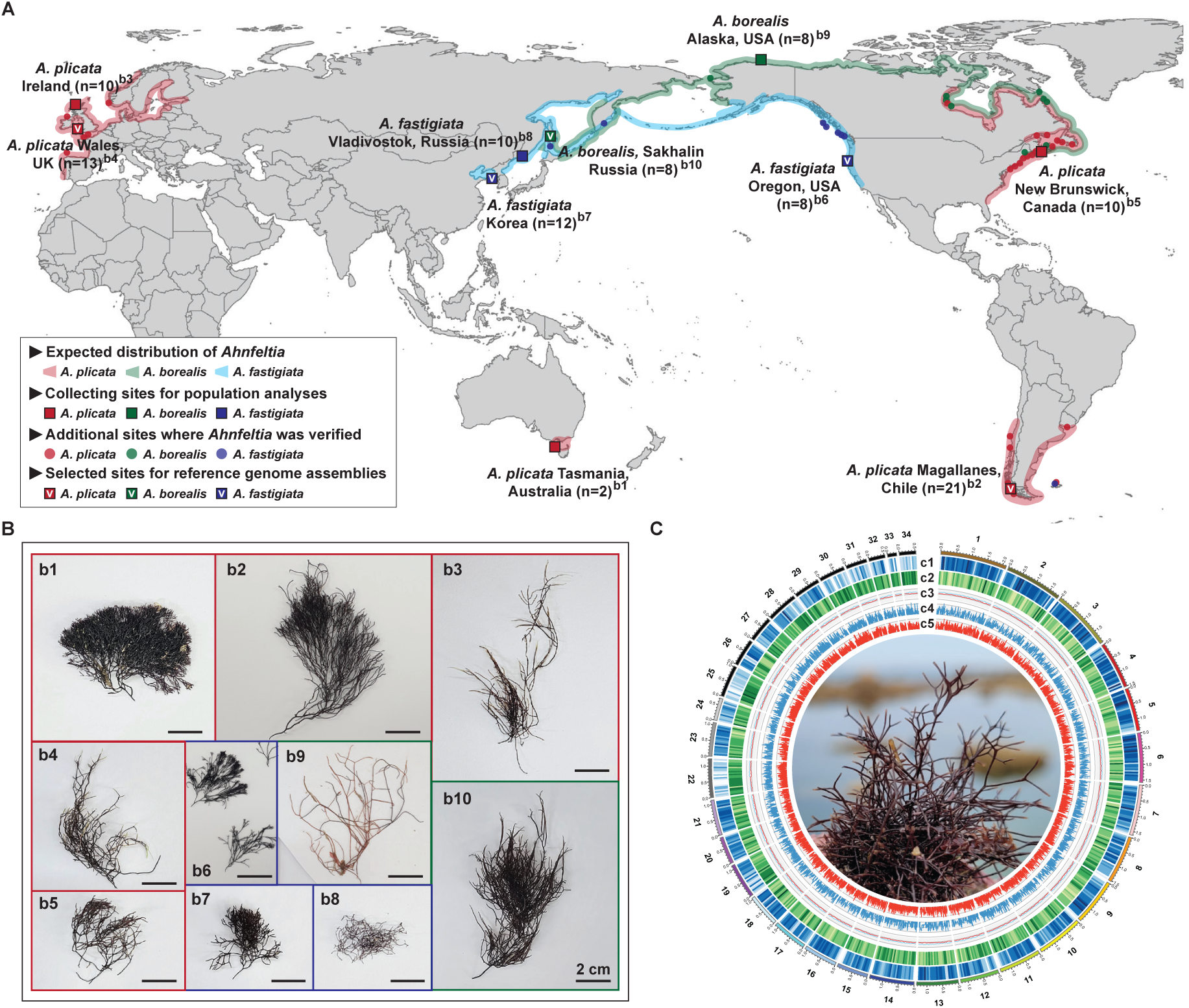
Sampling sites, morphology, and chromosome-level genome assembly of *Ahnfeltia*. (A) Sampling locations of *Ahnfeltia* populations included in this study. Ten populations were sampled across the global distribution range of the genus. White check marks indicate populations selected for chromosome-level nuclear genome assemblies, and square symbols indicate populations used for population genomic analyses. Previously reported *Ahnfeltia* sampling sites supported by molecular evidence are shown as circles, and the expected distribution of the genus is inferred from these records. (B) Morphology of collected *Ahnfeltia*: (b1) *A. plicata*, Tasmania; (b2) *A. plicata*, Chile; (b3) *A. plicata*, Ireland; (b4) *A. plicata*, New Brunswick; (b5) *A. plicata*, Wales; (b6) *A. fastigiata*, Oregon; (b7) *A. fastigiata*, Korea; (b8) *A. fastigiata*, Vladivostok; (b9) *A. borealis*, Alaska; (b10) *A. borealis*, Sakhalin. All scale bars represent 2 cm. (C) Features of the assembled genome. Circos plots show (c1) gene density, (c2) transposable element (TE) density, (c3) GC and AT content, (c4) Kimura divergence of retrotransposons, and (c5) Kimura divergence of DNA transposons.

To estimate genome size and heterozygosity of *Ahnfeltia*, we conducted *k*-mer analyses. The results revealed largely homozygous genomic content, although *k*-mer distributions for some individuals exhibited a subtle secondary peak, indicating residual heterozygosity rather than complete homozygosity (fig. S1 and S2; further information is described in Supplementary Text S1). The estimated genome sizes for the three *Ahnfeltia* species ranged from approximately 30 to 40 Mbp (fig. S1). Bacterial contamination levels were estimated to range from approximately 11.54-69.23% of PacBio reads; however, the remaining data still provided 115-415x coverage of the expected genome size (table S4). Contigs were removed if more than 80% of their predicted genes showed ≥90% amino acid identity to prokaryotic sequences in BLASTp searches (*27*).

The assemblies covered most chromosomes in each of the five genomes (Fig. 1C and fig. S3). However, the most complete telomere to telomere (T2T) assembly was obtained for *A. plicata* UK genome (fig. S4). The assembled genome sizes were 39.2-40.3 Mbp, similar to the result of size estimation (fig. S1). Between 8,246-8,374 genes were predicted in each of the *Ahnfeltia* genomes, which is less than observed in other published genomes of multicellular red algae such as *Bostrychia moritziana* and *Kappaphycus striatus* (table S4; *28, 29*). BUSCO analysis of all five *Ahnfeltia* genomes revealed completeness between 74.9-77.7% based on the Eukaryota odb10 database. This completeness is comparable to previous reports for red algae (*28, 29*), likely reflecting lineage-specific gene loss and the limited representation of rhodophyte orthologs in the BUSCO dataset (table S5). Approximately 90% of predicted genes were functionally annotated using complementary homology-based approaches (see Supplementary Methods S1, fig. S5, and table S6).

### Genomic conservation and low evolutionary rates

A central question raised by *Ahnfeltia* is how such an ancient lineage has remained so species-poor despite its broad geographic distribution and long evolutionary history. We hypothesize that this genus has persisted by tracking a remarkably stable ecological niche through time, while retaining a highly conserved phenotype adapted to intertidal and shallow rocky-shore habitats. If so, long-term occupation of these physically stressful and environmentally variable habitats may have favored evolutionary conservation in *Ahnfeltia* rather than promoting diversification. To test this hypothesis, despite the absence of a fossil record for *Ahnfeltia*, we reconstructed its evolutionary history using a combination of a) phylogenetic reconstructions, b) MCMC-based divergence time estimation, c) population structure analyses, d) fixation index (F_ST_) estimates, e) historical changes in effective population size (Ne), f) linkage disequilibrium (LD) decay analyses, and g) ecological niche modelling.

Nuclear and organelle (plastid and mitochondria) genome phylogenies of all sequenced *Ahnfeltia* species revealed a consistent divergence pattern characterized by an exceptionally low substitution rate within the Rhodophyta (Fig. 2A, fig. S6, and S7). Correspondingly, the number of recognized species is fewer, and their genetic distances inferred from mitochondrial and plastid phylogenies are markedly lower than those observed in other red seaweed genera (Fig. 2B and fig. S8A). The overall divergence of *Ahnfeltia* from the red algal common ancestor is lower than that observed in many other genera in Florideophyceae and Bangiophyceae. In addition, branch lengths from the most recent common ancestor of *Ahnfeltia* to each extant species are extremely short, indicating limited sequence divergence within the clade (Fig. 2C and fig. S8B). Together, these patterns suggest that *Ahnfeltia* represents an ancient lineage, yet has undergone only a small amount of genetic change since its divergence.

**Fig. 2.**
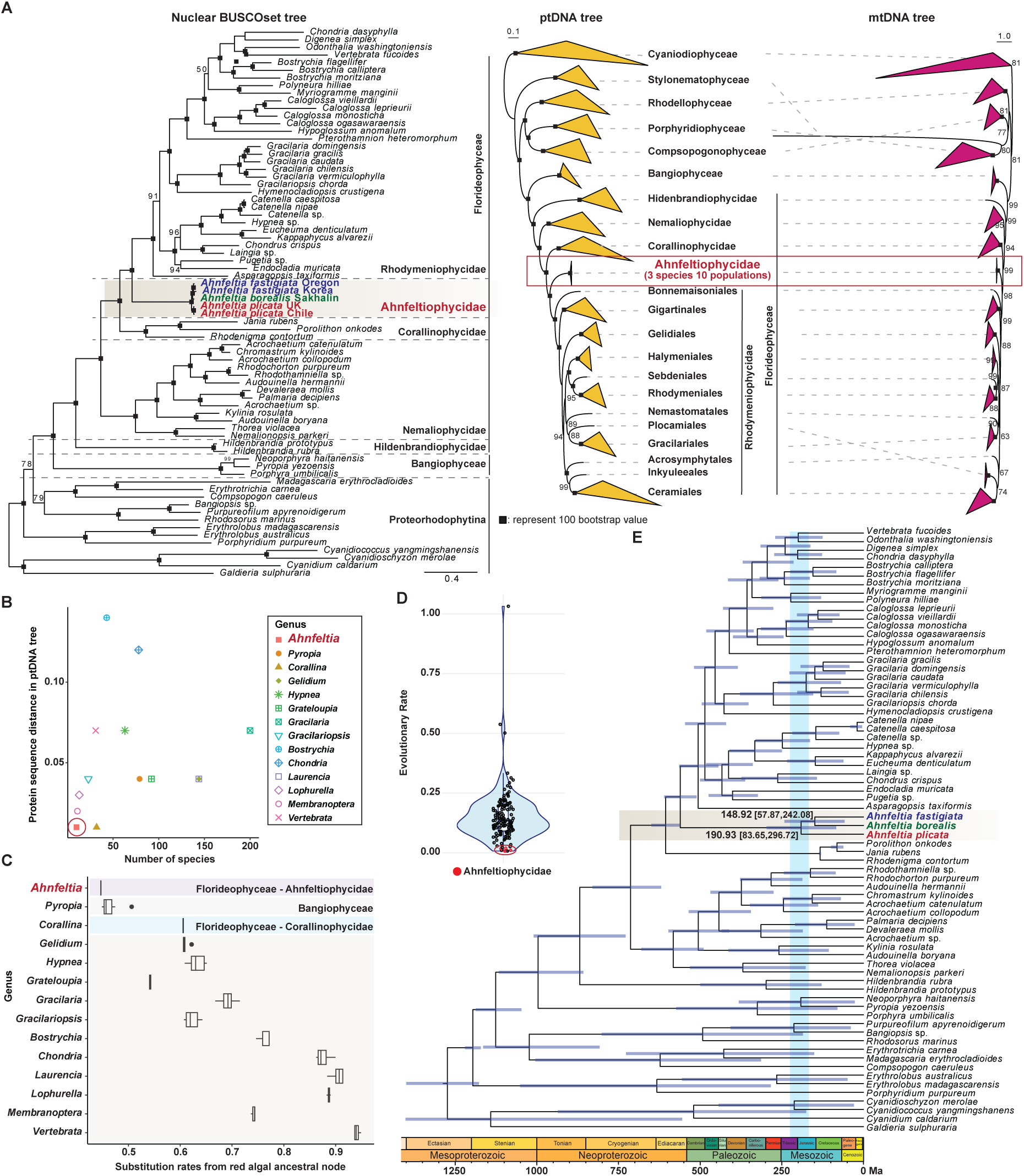
Phylogenomic relationships and divergence time estimates in *Ahnfeltia*. (A) Phylogenetic trees reconstructed from nuclear, plastid, and mitochondrial datasets. The *Ahnfeltia* clade shows extremely short branches and low substitution rates across all phylogenies. (B) Comparison between the number of described species and protein-sequence genetic distances inferred from plastid phylogeny. (C) Branch lengths of *Ahnfeltia* and other red seaweed genus measured from the ancestral node in the plastid phylogeny. (D) Relative evolutionary rates of *Ahnfeltia* compared with other red algal lineages inferred from divergence time estimates. (E) Divergence time estimation of *Ahnfeltia* species based on MCMC analysis using nuclear BUSCO gene sets, suggesting that diversification of the genus occurred approximately 190-148 Ma.

This hypothesis was further tested using Bayesian MCMC analyses based on the nuclear BUSCO gene set. The evolutionary rate in *Ahnfeltia* was estimated to be between 6.8x10^-9^ and 1.69x10^-8^ substitutions per site per year (Fig. 2D), which is similar to the rate estimated in the red seaweed *Pyropia yezoensis*: 2.97 × 10□□ per site per generation in the sexual cycle and 4.37 × 10□□ in the asexual life cycle (*30*). As no other estimates are available for red seaweeds, we compared relative evolutionary rates inferred under the same Bayesian MCMCtree framework and calibration scheme (Supplementary Text S2). Within this context, *Ahnfeltia* shows the lowest evolutionary rate among the studied red algal lineages (Fig. 2D).

*Ahnfeltia* speciation likely occurred approximately between 190 and 148 Ma within an ancient lineage that dates back to 500 Ma (Fig. 2E). This speciation time estimate is consistent with general red algal evolutionary trends (*9*). In particular, the 190 Ma estimate coincides with the breakup of Pangea (∼200 Ma), a period associated with a significant increase in nutrient supply especially in coastal regions (*31*) caused by newly emerging coastal margins. In this period, the species diversity of brown algae also significantly increased, corroborating the hypothesis that Pangea’s breakup may also have driven the speciation in *Ahnfeltia* (*32, 33*).

The low genetic diversity of *Ahnfeltia* species was corroborated by our comparative analysis of the assembled genomes. Although some differences were observed among species (Supplementary Text S3), the genomes are largely conserved. Nearly all scaffolds from the five assemblies could be colinearly aligned, apart from four translocated regions in the *A. plicata* Chile genome (Fig. 3A). Genome sizes of *A. plicata* and *A. borealis* were similar (ca. 38 Mbp); however, the assembled genomes size of *A. fastigiata* was approximately 2 Mbp larger than those of two species, which was primarily attributed to the expansion of TEs (Fig. 3B and fig. S9). LTRs and DNA/hobo-activator were largely responsible for the genome expansion of *A. fastigiata* (Fig. 3C, fig. S10, and table S7). Based on the distribution of the Kimura distance, TE insertions in the *A. fastigiata* genomes occurred relatively recently (≤ 107.6 million years), but close to the estimated speciation time of *A. fastigiata* (ca. 148 Ma; Fig. 3C). This result suggests that increased TE activity likely contributed to the divergence of *A. fastigiata* from the common ancestor of the *A. borealis*/*A. fastigiata* lineage.

**Fig. 3.**
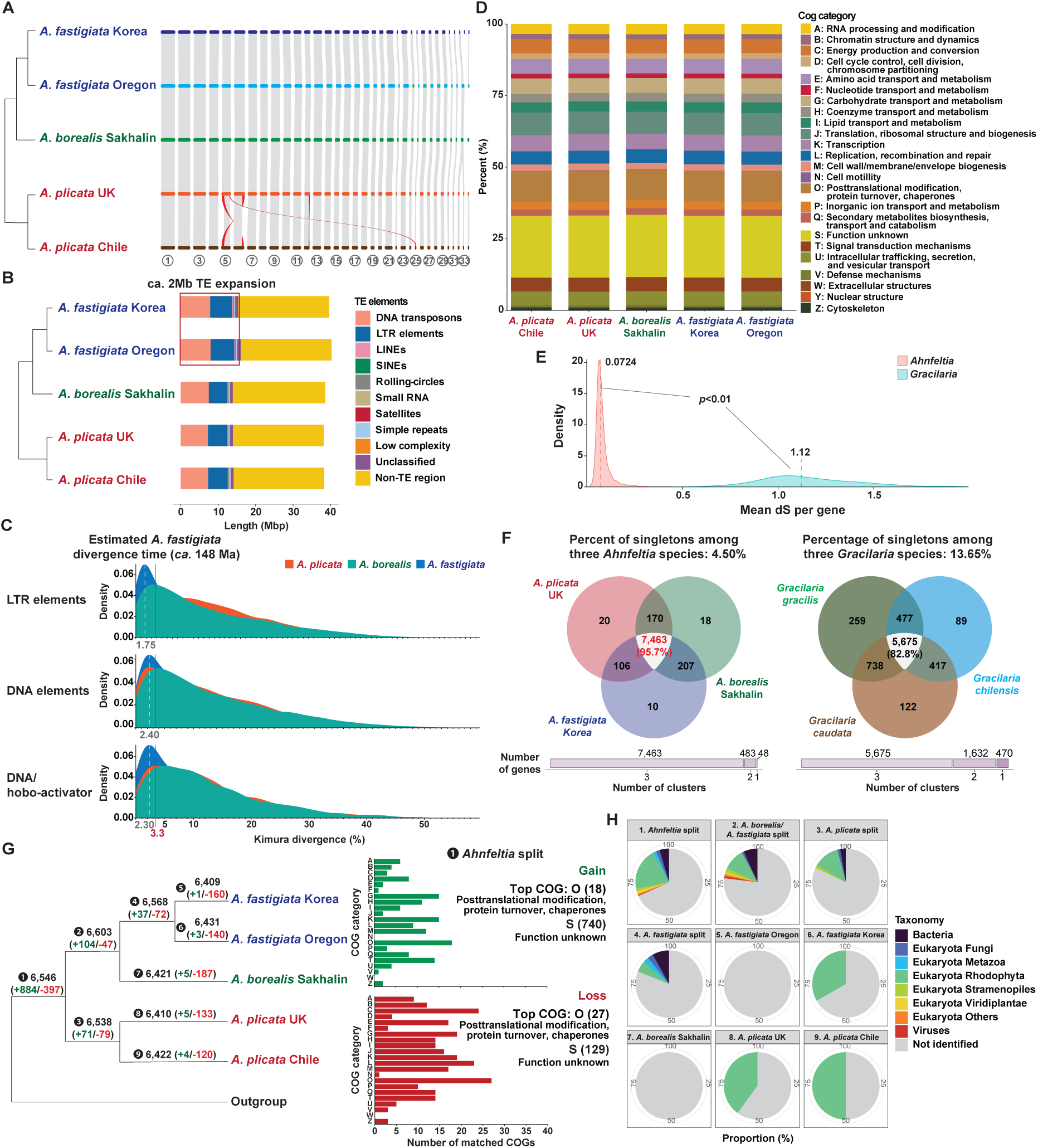
Genome structure and comparative genomic features of *Ahnfeltia*. (A) Gene synteny among *Ahnfeltia* genomes. Nearly all contigs were collinearly aligned, with the exception of four translocated regions in the *A. plicata* Chile genome. (B) Expansion of transposable elements (TEs) in *A. fastigiata* genomes, resulting in an approximately 2 Mbp increase in genome size. (C) Recently expanded TE families in *A. fastigiata*, including long terminal repeats (LTR) and DNA elements (particularly DNA/hobo-activator). (D) Functional classification of predicted genes based on Clusters of Orthologous Groups (COGs). (E) Pairwise comparisons of synonymous substitutions rates (dS) between *Ahnfeltia* and *Gracilaria* species, showing significantly lower substitution rates in *Ahnfeltia* (Wilcoxon rank-sum test, *p*<0.01). (F) Comparison of singleton gene content between *Ahnfeltia* and *Gracilaria* species. Variation among *Ahnfeltia* genomes is limited (4.5%) compared to *Gracilaria* (13.65%). (G) Gene gain and loss patterns inferred from orthologue clustering and Dollo parsimony analysis. (H) Candidate horizontally transferred genes (HGTs) identified in *Ahnfeltia* species and populations.

The gene content of all five assemblies also exhibited high conservation. The five genomes share more than 95% conserved core functions (≥ 7,400 genes, Fig. 3D). This high level of conservation is particularly striking when compared with the genomes of *Gracilaria*, one of the most divergent lineages within red algae (*11*). Genetic divergence among *Ahnfeltia* species (mean dS = 0.0724) is significantly lower than that observed among *Gracilaria* species (mean dS = 1.12, p < 0.01; Fig. 3E). Singletons specific to only one species of *Ahnfeltia* represent less than 5% of all protein encoding genes (Fig. 3F and fig. S11). Despite this high conservation, some gains and losses were identified (Fig. 3G and fig. S12). The most enriched COG categories among gained and lost genes were related to posttranslational modification, protein turnover, and chaperones (Fig. 3G). Several of these gained genes (10%) showed potential evidence of horizontal gene transfer (HGT) from bacteria (4.75%), fungi (1.70%), or metazoans (1.26%; Fig. 3H and table S8).

### Geographic isolation and limited populational divergence

Given the strong genomic conservation revealed by our phylogenetic and comparative genome analyses, we next examined evolutionary patterns at the population level. Population structure analyses showed strong geographic structuring across the global distribution of *Ahnfeltia*. Principal component analyses (PCAs) revealed clear genetic separation among the three *Ahnfeltia* species and ten populations, with minimal overlap. A clear divide was observed between Northern and Southern Hemisphere populations of *A. plicata*, as well as among populations of *A. fastigiata* and *A. borealis* (Fig. 4A and fig. S13 to S15). ADMIXTURE analyses were largely consistent with the PCA results across a range of K values (Fig. 4B), although minor shared ancestry components (∼0.3) were detected in some North Atlantic populations (fig. S16 and S17).

**Fig. 4.**
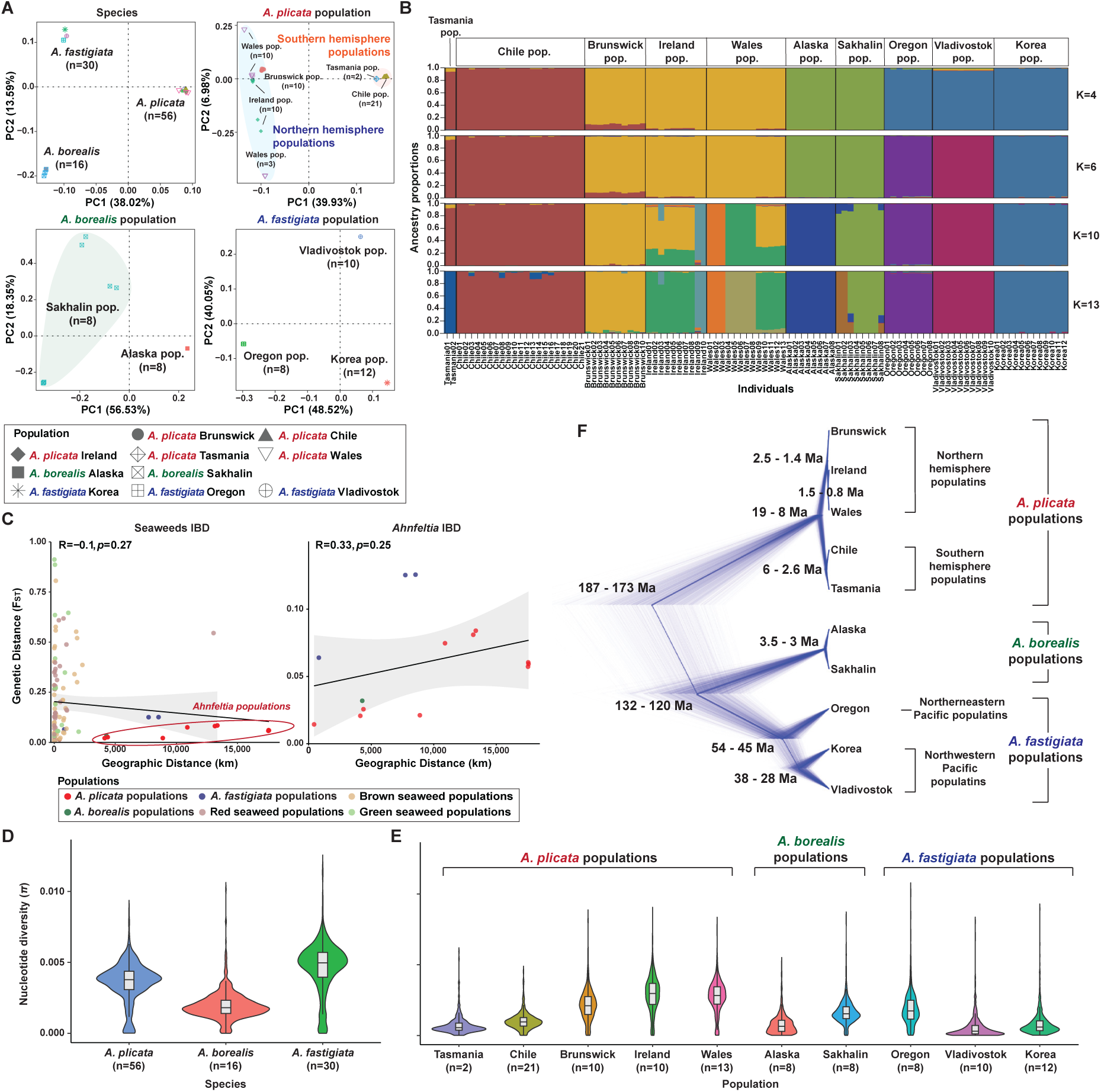
Population structure and genetic diversity of *Ahnfeltia*. (A) Principal component analysis (PCA) of three *Ahnfeltia* species across ten populations. (B) Population admixture structure at K=4, K=6, K=10, and K=13. Both PCA and admixture analyses reveal clear genetic differentiation among *Ahnfeltia* populations, with limited evidence of gene flow. (C) Isolation by distance (IBD) analysis of *Ahnfeltia* and other seaweed species, showing weak correlation between genetic and geographic distance in *Ahnfeltia* based on Pearson correlation test (left: *p*=0.27; right: *p*=0.25) (D) Nucleotide diversity (π) within *Ahnfeltia* species. (E) Nucleotide diversity (π) within *Ahnfeltia* populations. (F) Divergence time estimation of *Ahnfeltia* populations based on SNAPP using pruned SNP datasets.

Despite this clear geographic structure, we detected no significant signal of isolation-by-distance (IBD). Geographic separation would generally be expected to restrict gene flow and promote genetic divergence over evolutionary time, yet pairwise *F*_ST_ did not increase significantly with distance (*p* = 0.25), even across comparisons spanning almost the entire globe (Fig. 4C). Furthermore, the overall level of differentiation was much lower than other seaweed species (Fig. 4C). These results indicate that the *Ahnfeltia* species complex has not undergone typical allopatric speciation as commonly observed across diverse taxa, suggesting that geographic distance alone has not been the primary factor structuring genetic variation.

Among the three species, *A. plicata* provides the clearest example of limited population divergence across a vast geographic range. It was sampled from both the Northern and Southern Hemispheres and spans the broadest latitudinal and longitudinal range in the dataset. Nevertheless, its overall nucleotide diversity was lower than that of *A. fastigiata* (Fig. 4D), indicating comparatively weak divergence among populations. Southern Hemisphere populations from Chile and Tasmania also had lower nucleotide diversity than those from the Northern Hemisphere (Fig. 4E). Divergence between Northern and Southern Hemisphere populations was estimated to date to the Miocene, around 19 to 9 Ma (Fig. 4F and fig. S18). During this period, global cooling may have facilitated transequatorial dispersal by reducing thermal barriers in equatorial regions (*34*). Periods of moderate global cooling likely provided optimal conditions for dispersal of *Ahnfeltia* by reducing thermal barriers without extensive ice coverage. This interval also coincided with the establishment of the modern Antarctic Circumpolar Current and the expansion of Antarctic glaciation (*35*). These changes may have promoted dispersal into suitable cold-water coastal habitats, consistent with the present-day distribution of *A. plicata* in cooler oceans (*12, 14, 16*).

Ancestral populations of *A. borealis* and *A. fastigiata* were already established by the Early Cretaceous (approximately 132-120 Ma; Fig. 4F), based on divergence time estimates, when global climates were predominantly greenhouse and high-latitude regions were considerably warmer and largely ice-free (*36*). Given its preference for cold conditions, ancestral populations likely persisted within geographically shifting regions that maintained suitable temperatures, rather than inhabiting uniformly warm environments. However, subsequent global cooling during the Cenozoic likely restructured the distribution of suitable habitats. Repeated glacial-interglacial cycles may have driven alternating phases of contraction and expansion, with long-term persistence largely restricted to high-latitude regions. Such patterns of range contraction and redistribution following Cenozoic cooling have also been reported for several boreotropical taxa and Antarctic benthic invertebrates (*37, 38*). Two *A. borealis* populations appear to have undergone more recent differentiation. The estimated divergence between the Alaska and Sakhalin populations (3-3.5 Ma) corresponds to a period of pronounced sea-level fluctuations during the late Pliocene, when repeated changes in connectivity across the Bering Strait may have restricted gene flow between the North Pacific regions (*39, 40*).

*A. fastigiata* is primarily distributed in the North Pacific and has recently been recorded from the Falkland Islands with molecular evidence (table S1). Divergence among North Pacific *A. fastigiata* populations is inferred to have occurred earlier, during a significant cooling period between ca. 56 and 28 Ma (Eocene & Oligocene; Fig. 4F) when Earth transitioned from a warm ‘greenhouse’ to an ‘icehouse’ climate (*41*). The expansion of cooler marine environments during this period may have facilitated the dispersal of cold-adapted taxa such as *Ahnfeltia*. Subsequently, the development of warmer currents likely acted as thermal barriers that reinforced geographic isolation. For example, the Korea population of *A. fastigiata* appears to have followed a post-glacial retreat pattern similar to that observed in several marine organisms in the Yellow Sea, shifting toward the Bohai Sea after deglaciation (*13, 42, 43*).

Most genetic variation in *Ahnfeltia* was structured among species rather than among populations. Molecular variance tests (AMOVA) indicated that 85.15% of the total variance was explained by interspecific differences, whereas only 7.63% was attributable to variation among populations within species (table S9), indicating limited population-level differentiation. Overall, pairwise mean F_ST_ values were low to moderate (0.03-0.13), suggesting weak genetic structuring among populations. Relatively higher differentiation (0.13 mean F_ST_) was observed between Northeastern and Northwestern Pacific populations of *A. fastigiata* (e.g., Oregon vs. Korea; Oregon vs. Vladivostok). In particular, this divergence was not driven by a small number of highly differentiated loci but instead reflected a genome-wide elevation of F_ST_ across most loci. Consistently, most loci exhibited low dN/dS ratios (< 0.66), indicating the predominance of purifying selection. These patterns suggest that the observed low differentiation is primarily shaped by demographic processes and restricted gene flow (Fig 5A, 5B, and fig. S19).

**Fig. 5.**
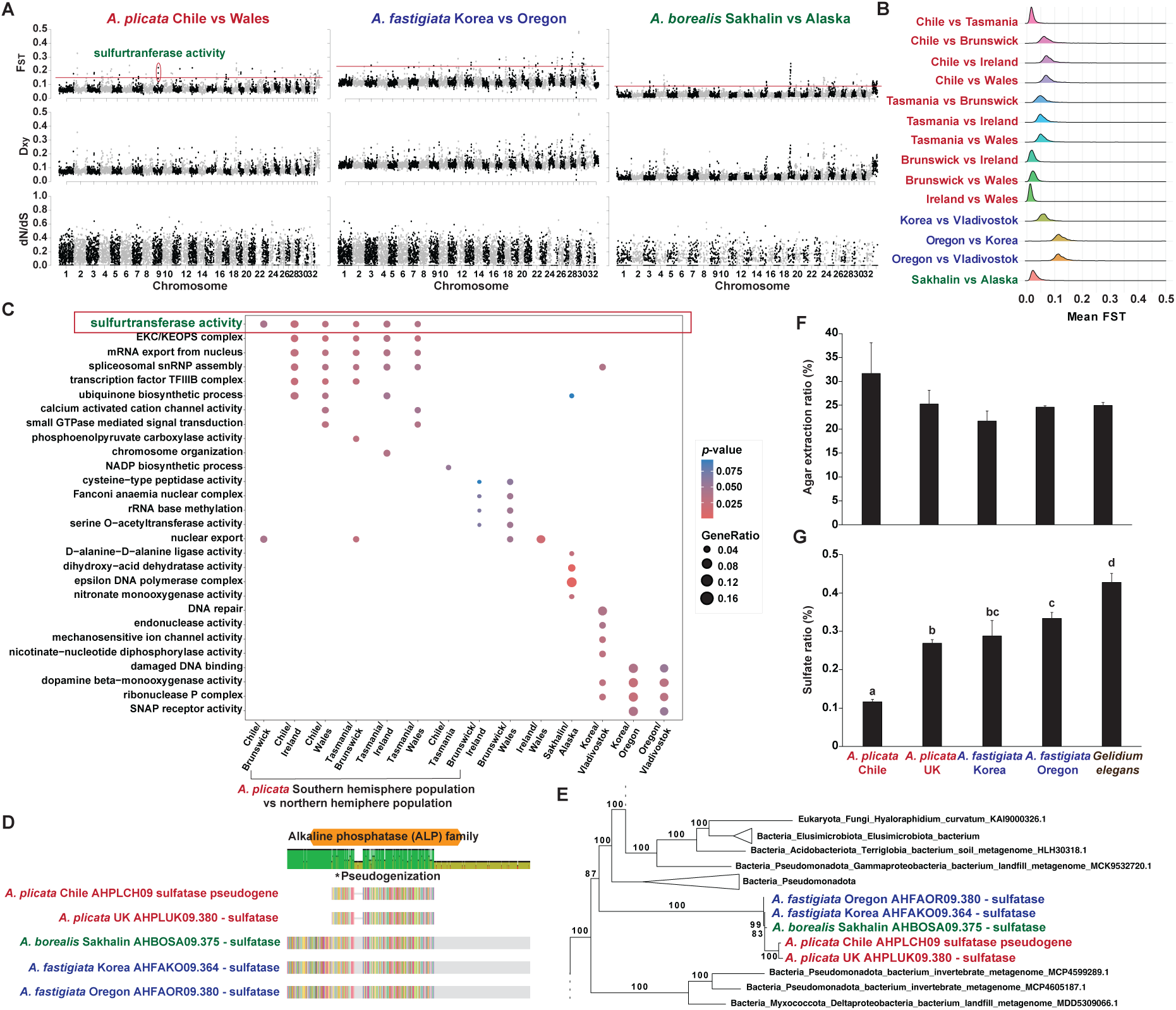
Genomic divergence and functional variation among *Ahnfeltia* populations. (A) Genomic landscape of fixation index (FST), genetic distance (Dxy), and nonsynonymous-to-synonymous substitution ratios (dN/dS) based on pairwise comparison of *Ahnfeltia* populations using the *A. plicata* Chile, *A. borealis* Sakhalin, and *A. fastigiata* Oregon reference genomes (B) Overall FST among *Ahnfeltia* populations. (C) Gene Ontology (GO) enrichment of highly divergent loci (high FST) for each pairwise comparison. One highly divergent region between Southern and Northern Hemisphere populations was associated with sulfotransferase activity. (D) Sequence alignment of a sulfatase pseudogene identified in *A. plicata* Chile with functional sulfatase genes from other *Ahnfeltia* genomes. (E) Phylogeny of sulfatase genes identified in *Ahnfeltia* genomes. (F) Comparison of agar properties among two *Ahnfeltia* species from four populations and *Gelidium*. (G) Comparison of agar sulfate content. Letters indicate significant differences based on Tukey’s post hoc test following ANOVA (p < 0.01). Error bars represent standard deviations of duplicate samples in (F) and triplicate samples in (G).

The most tractable genetic and phenotypical differentiation was observed in the *A. plicata* Chile population. Among the significantly divergent loci, one was associated with sulfurtransferase activity (Fig. 5C and table S10). In addition, a putative bacterial-derived sulfatase (EC 3.1.6.-) was pseudogenized by a single 1-bp deletion, resulting in a frameshift and premature truncation (Fig 5D, 5E, and fig. S20). These genetic variations may contribute to the reduced sulfur content observed in this strain compared with *Gelidium elegans* and other *Ahnfeltia* populations (Fig. 5F and 5G; *44*). Given that reduced sulfur content is known to influence agar quality, this population may therefore have potential commercial value (*17–19*). However, these localized genetic differences do not appear to correspond to species-level differentiation. Taken together, the lack of a clear IBD pattern, low nucleotide diversity, and deep divergence times is consistent with the possibility that large-scale Earth system changes structured the evolutionary history of the genus rather than gradual geographic isolation.

### Evidence of a high level of homozygosity and long-term niche suitability

Genome-wide estimates of the inbreeding coefficient (F_IS_; Fig. 6A) and observed homozygosity (Ho; Fig. 6B and fig. S21) revealed an overall high level of homozygosity of allelic sites. In addition, extensive runs of homozygosity (ROH) were detected across *Ahnfeltia* genomes (Fig. 6C and fig. S22), and the length of homozygous blocks ranged from 20 to over 200 kbp (Fig. 6D). Their proportion as a measure of inbreeding (F_FROH_) was consistently high across populations, ranging from 0.42 to 0.76 (Fig. 6E), suggesting significant inbreeding and a small effective population size (Ne). Indeed, we identified a recent reduction of Ne, which likely facilitated the observed homozygosity (Fig. 6F and fig. S23). Consistent with the PCA and admixture analyses, these findings suggest that *Ahnfeltia* lineages have long persisted as geographically isolated populations with limited gene flow. This evolutionary a history likely contributed to the high homozygosity observed across the genus.

**Fig. 6.**
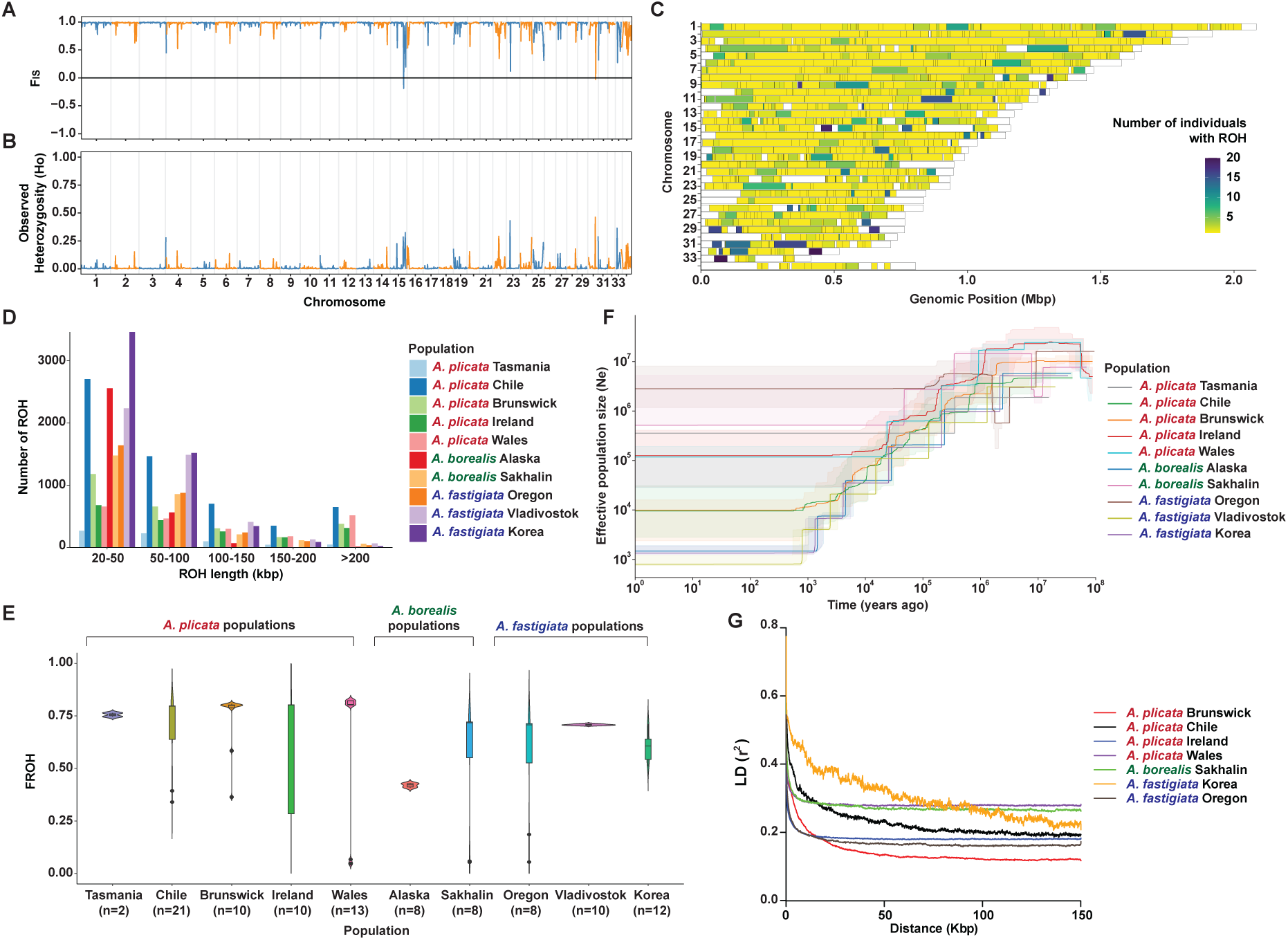
Patterns of homozygosity and demographic history in *Ahnfeltia*. All analyses were conducted using the *A. plicata* UK genome as the reference. (A) Inbreeding coefficient (FIS) across *Ahnfeltia* populations. (B) Observed homozygosity (Ho) across populations. (C) Runs of homozygosity (ROH) identified in *Ahnfeltia* genomes. (D) Number and size distribution of ROH blocks in *Ahnfeltia* populations. All these analyses indicated elevated levels of inbreeding within *Ahnfeltia* population. (E) Proportion of the genome in runs of homozygosity, used to estimate the inbreeding coefficient (FROH) across *Ahnfeltia* populations. (F) Changes in effective population size (Ne) inferred using Stairway Plot 2. (G) Linkage disequilibrium (LD) decay of *Ahnfeltia* populations.

The “new” environments into which *Ahnfeltia* migrated were probably not fundamentally different from the ancestral habitat, suggesting limited ecological divergence despite broad geographic expansion. The high homozygosity observed across most *Ahnfeltia* species and populations is more likely to reflect small effective population sizes, founder effects, and prolonged geographic isolation than selection associated with colonization of novel habitats. Heterogeneous patterns of linkage disequilibrium (LD) decay, with particularly elevated LD in the Chilean population of *A. plicata* and the Korean population of *A. fastigiata*, further indicate region-specific demographic trajectories within *Ahnfeltia*, consistent with but not fully explained by divergence time estimates among populations. If genetic variation had simply accumulated following populational divergence, early diverged populations would be expected to exhibit high LD decay (Fig. 6G and fig. S24).

Over the past ∼200 million years, *Ahnfeltia* likely persisted within suitable cold-water habitats, with populational dispersal potentially occurring during cooler periods. Climatic warming in lower-latitude coastal regions may have intermittently reduced habitat availability, likely leading to bottlenecks and local extinctions, especially in small, genetically depauperate populations. Together with continental drift, these processes likely promoted long-term geographic isolation, with persistence restricted to suitable cold, nutrient-rich environments. Accompanied life-history traits such as a long-life cycle of over 3 years, their occasional asexual reproduction, and resilient phenotypes may have contributed to their persistence without significant genetic changes over a long evolutionary time in a well-defined ecological niche (*15, 21, 22;* Supplementary Text S4).

To test whether suitable conditions would have been consistently available to *Ahnfeltia* over its long evolutionary history, we projected the realized niche for the genus through time using several series of HadCM3L palaeoclimate simulations (*45, 46* see Supplementary Methods S6 for more details). We compared niche dynamics for *Ahnfeltia* to the red algal genus *Gracilaria* as a control because, like *Ahnfeltia*, it is an agarophyte within the class Florideophyceae, but comprises more than 250 described species with higher genetic divergence and differs ecologically by exhibiting a predominantly warm and tropical-water global distribution (see Supplementary Text S5 for details). Accordingly, niche analyses were conducted at the genus rather than species level to enable comparison of broad-scale niche patterns between the two genera (see Supplementary Methods S6). We projected models onto multiple time intervals over the last 505 Ma for *Ahnfeltia* and 300 Ma for *Gracilaria*, reflecting molecular estimates of the latter’s crown lineage divergence (*9*). Models were constructed using 342 and 419 present-day occurrences for *Ahnfeltia* and *Gracilaria*, respectively, confirmed through genetic identification (table S1 and S3).

Paleo-niche projections indicate that suitable conditions for *Ahnfeltia* were consistently available at high latitudes over the past 505 million years, with higher suitability concentrated toward cold-temperate and polar regions (≥ 60°) regardless of plate tectonic changes (Fig. 7A and 7B). In contrast, *Gracilaria* exhibited broader environmental suitability across lower latitudes throughout geological time (Fig. 7A, 7B, and fig. S25 to S32). Our niche modeling suggests that *Ahnfeltia* occupies colder conditions than *Gracilaria*, with only 37% overlap in climate space (Fig. 7C, table S11). Notably, suitable conditions for *Ahnfeltia* appear to have been more spatially contiguous through time compared to those of *Gracilaria* (Fig. 7D). This persistent availability of suitable habitats may have reduced opportunities for geographic isolation among populations, thereby limiting the accumulation of genetic divergence. Broadly, the proportion of coastline area predicted as suitable for *Ahnfeltia* increased through the Mesozoic and declined through the Cenozoic alongside long-term global cooling; however, over the past ∼200 million years, shorter-term fluctuations indicate that intervals of reduced mean sea surface temperature were frequently associated with increases in suitability (Fig. 7E). These findings are consistent with the relatively low levels of genetic differentiation observed across *Ahnfeltia* and suggest a role for long-term environmental stability in constraining diversification.

**Fig. 7.**
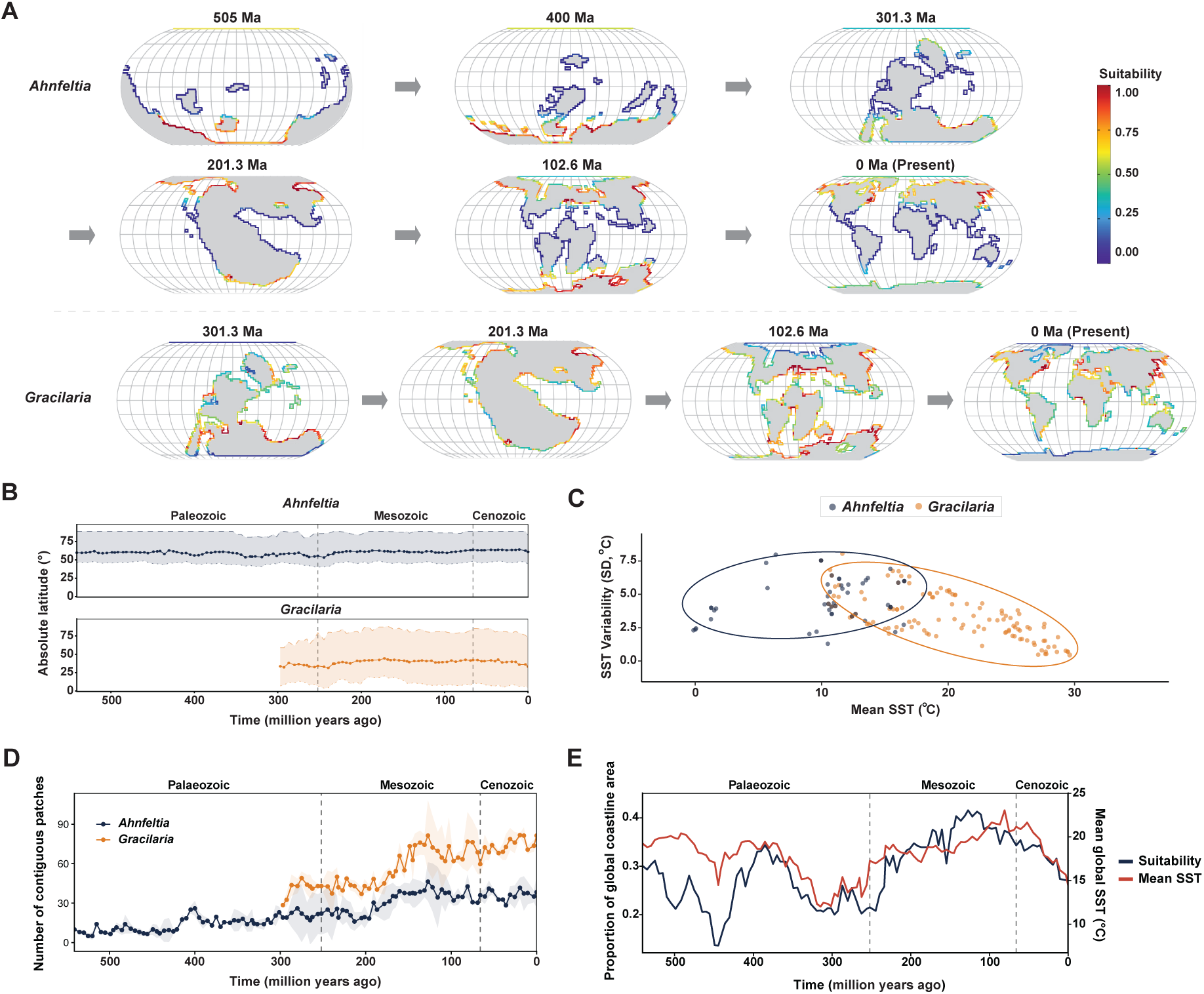
Climatic suitability and thermal niche dynamics of *Ahnfeltia* and *Gracilaria* across the Phanerozoic. (A) Predicted climatic suitability for *Ahnfeltia* and *Gracilaria* in coastal regions under palaeoclimate reconstructions at representative time intervals across the Phanerozoic (0, 102.6, 201.3, 301.3, 400, and 505 Ma). Suitability values are shown on a cloglog scale (0-1). (B) Mean absolute latitude of climatically suitable areas through time for *Ahnfeltia* and *Gracilaria*. Lines represent the mean latitude of predicted suitable cells for each time slice. Dashed and dotted lines indicate the 0.95 and 0.05 quantile latitudinal limits, respectively, with shaded ribbons showing the range between these bounds. Background labels denote major geological eras. (C) Climatic hypervolumes of *Ahnfeltia* and *Gracilaria* in thermal environmental space defined by mean sea surface temperature (SST) and SST variability (SD). Points represent observed occurrences. (D) Mean number of contiguous environmentally suitable patches through time for *Ahnfeltia* and *Gracilaria*. Solid lines and points represent the mean number of patches derived from binary suitability maps for each time slice, averaged across simulations, with shaded ribbons indicating variability among simulations. (E) Temporal trends in global mean SST and the area of suitable environmental conditions for *Ahnfeltia* derived from binary ENM projections.

## Conclusion

Although *Ahnfeltia* lacks a fossil record to corroborate its inferred evolutionary history, our integrated analyses, combining high-quality genome sequencing, phylogenomic analyses, population genetics, ecology, and niche modeling provides a coherent picture of long-term evolutionary stasis in this ancient red algal lineage. Across species and populations, *Ahnfeltia* exhibits remarkable genomic conservation, strong geographic isolation, high homozygosity, and limited diversification despite its deep evolutionary age and broad global distribution. These findings suggest that *Ahnfeltia* persisted through long-term niche tracking. In other words, the lineage appears to have survived by following geographically shifting but environmentally similar cold-temperate rocky-shore habitats through major climatic and tectonic change, rather than through extensive ecological divergence or repeated speciation. Even during global cooling events, their strict habitat specialization appears to have constrained geographic expansion and kept diversification low, with populations remaining stable in suitable cold-temperate habitats, as supported by our niche modeling (Fig. 7D and 7E).

Given its long evolutionary history, it is possible that ancestral *Ahnfeltia* was more ecologically flexible than extant species, potentially tolerating a broader range of temperatures. Although the traits of ancestral populations remain uncertain, they may have experienced several extinctions during events such as the Ordovician-Silurian (ca. 445 Ma) and Permian-Triassic (ca. 252 Ma), when suitable habitats were likely reduced (Fig. 7E). Following past possible extinction events, extant species and populations at least appear confined to cold-temperate habitats without broader environmental tolerance. Given ongoing global warming, this long-term evolutionary stasis that has supported the persistence of *Ahnfeltia* may now reduce the capacity of its populations to cope with rising temperatures in cold-water habitats. Thus, *Ahnfeltia* may serve as a bellwether for the loss of seaweeds because of changing coastal ecosystems, especially at higher latitudes, which are most affected by global warming.

## Material and methods

### Sample collection, genome sequencing, and genome assembly

Fresh samples of three *Ahnfeltia* species were collected from five populations: *A. plicata* - Seno Skyring, Rio Verde, Magallanes, Chile and Freshwater West, Pembrokeshire, *A. borealis* - Starobubskoe, Sakhalin, Russia, and *A. fastigiata* - Baeknyeong island, Korea and Seal Rock, Oregon, USA (table S3). All samples were thoroughly cleaned under a dissecting microscope prior to whole-genome and whole-transcriptome sequencing.

Approximately 10-30 μg of high-quality genomic DNAs were extracted using a modified cetyltrimethylammonium bromide (CTAB) method (*47*), and sent to JS-link Inc. (Seoul, Republic of Korea) for sequencing. PacBio Sequel I platform (Pacific Biosciences, Inc., Menlo Park, CA, USA) and Illumina NovaSeq 6000 platform (Illumina, San Diego, USA) were used to generate long-read data and short-read data, respectively. PacBio libraries were prepared using the SMRTbell Template Prep Kit (Pacific Biosciences), and Illumina paired-end libraries were prepared using the TruSeq DNA Nano 550bp Library Kit (Illumina). Two SMRT cells were sequenced, and approximately 10 Gbp of paired-end short-read data were generated.

For gene prediction, *Ahnfeltia* samples were placed under six conditions (4 °C, 15 °C, red light, blue light, white light, and dark) for at least six hours to induce various gene expression. Total RNA was extracted using RNeasy Plant Mini Kit (Qiagen, Hilden Germany) following manufacturer’s instruction. RNA-seq library were prepared using TruSeq Stranded mRNA Library Prep Kit (Illumina) and sequenced with Illumina NovaSeq 6000 (San Diego, USA).

Genome assembly was performed using Falcon (*48*) and NextDenovo (*49*) and polished with Illumina short reads using Pilon v1.24 (*50*). Transposable elements were soft-masked using RepeatModeler v2.0.3 (*51*) and RepeatMasker v4.1.2 (*52*). Gene prediction was carried out using EVidenceModeler v1.1.1 (*53*), incorporating results from BRAKER2 (*54, 55*), GeMoMa (*56*), and PASA (*57*). Genome assembly and gene model completeness were evaluated using BUSCO v4.1.2 (*58*). Detailed information is provided in Supplementary Methods 1.

### *Ahnfeltia* population re-sequencing and variant calling

102 individuals of three *Ahnfeltia* species from ten populations were collected from tidepools and intertidal zones for individual whole genome re-sequencing (table S2). In the case of eight individuals of *A. borealis* from Alaska, USA, two individuals of *A. borealis* from Sakhalin, and 10 individuals of *A. fastigiata* Russia, dried specimens were used for re-sequencing. Before sequencing, all samples were thoroughly cleaned with sterile seawater. For individual re-sequencing, about 1 μg of DNA for each sample was sent to the sequencing company and re-sequenced using TruSeq DNA Nano 550bp Library Kit and Illumina NovaSeq 6000 (Illumina, San Diego, USA). About 10 Gbp of paired-end reads were produced for each individual.

All sequence data of *Ahnfeltia* individuals were trimmed to remove adapters and low-quality bases using Trimmomatic v0.39 (*59*). Trimmed reads were then aligned to the *Ahnfeltia* genomes using BWA v0.7.17 (*60*). Aligned information was converted into bam files using Samtools v1.7 (*61*). PCR duplicates were marked and read groups were assigned with Picard v3.0.0 (*62*). HaplotypeCaller in GATK v4.1.9.0 with GVCF mode was performed for variant discovery and genotyping (*63*). To obtain high-confidence variant sites, hard filtering was conducted to exclude low-qualified variant sites using bcftools v1.9 (*64*) with the following parameters: FS>40.0 SOR>3 MQ<40 MQRankSum<-10.0 MQRankSum>10.0 QD<2.0 ReadPosRankSum<-4.0. A second round of filtering was conducted using VCFtools v0.1.17 (*65*) to retain only biallelic variants (--min-alleles 2 --max-alleles 2) with no missing data across individuals (--max-missing 1), a minimum quality score of 30 (--minQ 30), and a minimum read depth of 10 (--minDP 10). Finally, all variants were divided into SNPs and INDELs using GATK v4.1.9.0.

### Phylogenomic analyses and divergence time estimation

To evaluate phylogenetic relationships and divergence rates, phylogenomic analyses were conducted using published nuclear, plastid, and mitochondrial genomes of red algae. The genome datasets used in this study are listed in table S12. For nuclear genome phylogenetic analyses, two complementary approaches were applied. Because complete nuclear genome assemblies and protein annotations were not available for many taxa, fragmented genomic resources were used. To obtain high-confidence nuclear markers, complete single-copy BUSCO protein genes were extracted using BUSCO v4.1.2 (*58*). In parallel, to confirm consistency in phylogenetic signal, single-copy orthologous gene sets were obtained using OrthoFinder V2.5.2 (*66*). Protein sequences were concatenated and aligned using MAFFT v7.310 (*67*) and Geneious prime 2022.2.2 (https://www.geneious.com). Maximum-likelihood phylogenetic trees were constructed using IQ-TREE v1.6.8 (*68*).

To compare divergence rates among red algal lineages, protein sequence distances were estimated and contrasted with the number of described red algal species. Concatenated plastid and mitochondrial protein alignments were analyzed using MEGA X (*69*). We referred to Algaebase for the number of currently recognized red algal species (*11*). In addition, to evaluate species divergence within genera, branch lengths were calculated from ancestral nodes in the phylogenetic trees. Divergence times among *Ahnfeltia* species were estimated using MCMCtree (*70*). Detailed procedures and parameter settings for divergence time estimation are provided in Supplementary Methods S2.

To estimate divergence times among *Ahnfeltia* populations, SNP-based time inference was performed following Matschiner’s tutorial (github.com/mmatschiner/tutorials/). Independent SNPs (∼7,200 loci) were obtained by sampling one SNP every 3 kb from intergenic regions to minimize potential evolutionary bias associated with coding sequences and to reduce computational burden. We selected two individuals for each population, and vcf files were converted to PHYLIP format using vcf2phylip.py (*71*). An XML input file for SNAPP analysis was generated using the snapp_prep.rb script (*72*). Two divergence time calibration (the initial divergence within *Ahnfeltia* and the split between *A. borealis* and *A. fastigiata*) as well as the starting tree topology were derived from the MCMCtree results described above. SNAPP v.1.5.2 package in BEAST2 v.2.6.6 was run with 20,000,000 MCMC chains (*73*), and the first 20% of samples were discarded as burn-in. Convergence and parameter mixing were evaluated using Tracer v1.7.1 (*74*) All parameters showed effective sample size (ESS) values greater than 200, indicating adequate sampling and convergence.

### Comparative genomic analyses

Genomic and genetic evolution among *Ahnfeltia* species were comparatively analyzed. Orthologue analyses were conducted using OrthoVenn3 (*75*) and OrthoFinder v2.5.2 (*66*). Gene synteny among *Ahnfeltia* genomes was examined using the Python implementation of MCScan (*76*). TE composition was assessed based on the results of RepeatMasker v4.1.2 (*52*). Functional categories of *Ahnfeltia* gene sets were assigned using the Clusters of Orthologous Groups (COG) classification from eggNOG annotations (*77, 78*). Gene gain and loss were conducted using Count under the Dollo parsimony principle (*79*). Detailed methodological procedures are provided in Supplementary Methods S3.

### Population analyses

Given the heteromorphic triphasic life cycle of *Ahnfeltia*, we first assessed ploidy levels and data quality across all samples. The distribution of homozygous and heterozygous genotypes, per-site read depth, *k*-mer frequency spectra, and within-sample fixation index (FWS) were examined to distinguish haploid from potential diploid or mixed-stage samples and to evaluate clonality using moimix (*80*). Details are discussed in Supplementary Text S1. Population structure was investigated using PCA and admixture analyses in LEA (*81*).

Genomic differentiation and local adaptation of *Ahnfeltia* populations were evaluated by calculating FST (*65*) and Dxy (github.com/simonhmartin/genomics_general) in 20 kb sliding windows. To explore distinctive phenotypic characters of *Ahnfeltia* populations, agar extraction rate and sulfate composition of *Ahnfeltia* agar were measured, as described in Supplementary Methods S4. Coding-level selection between populations was assessed using KaKs_Calculator 2.0 (*82*). Within-population diversity and homozygosity were characterized based on nucleotide diversity (π), individual inbreeding coefficients (F), FIS, and runs of homozygosity (ROH) using plink v1.90b6.24 (*83*). Pairwise FSTs were compared with geographic distances to test isolation-by-distance (IBD). Linkage disequilibrium (LD) decay was estimated using PopLDdecay v3.41 (*84*). Historical changes in effective population size (Ne) were inferred using Stairway Plot v2 (*85*). Detailed information for population analyses is described in Supplementary Methods S5.

### Ecological Niche Modelling of *Ahnfeltia* and *Gracilaria* through the Phanerozoic

Genetically verified occurrence records of *Ahnfeltia* and *Gracilaria* were compiled from Barcode of Life Data System (BOLD), NCBI, and confirmed herbarium specimens (table S1 and 3), and subsequently cleaned using CoordinateCleaner v3.0.1 R package (*86*). Records with missing or invalid coordinates, duplicates, or terrestrial locations were removed. An additional unverified GBIF dataset for *Ahnfeltia* was included for sensitivity analyses.

Palaeoclimate data were derived from HadCM3L-M2.1D simulations (see Supplementary Methods 6 for more details) under Getech and Scotese palaeogeographic frameworks (*87, 88*), including both temperature-based (*89*) and CO₂-forced reconstructions (*90*). Data were interpolated to ∼100 km resolution. Monthly SST, salinity, and PAR at 5 m depth were extracted, and summary statistics (mean, min, max, range, SD) were calculated. Multicollinearity was assessed using VIF, retaining salinity (SD, max), SST (range, max), and PAR (SD, min).

Accessible areas were defined using distance-based buffers around occurrences. Target-group background sampling was implemented using Florideophyceae GBIF records. Presence and background points were spatially thinned based on Moran’s I. ENMs were fitted in Maxent using linear and quadratic features (*91*), testing regularization multipliers from 1-3. Models were evaluated using spatial cross-validation using blockCV v3.2-0 (*92*) and selected based on minimum AICc. Final models were replicated ten times with alternative background samples.

Models were projected onto Phanerozoic time slices for each genus (108 for *Ahnfeltia* to 505 Ma, 60 for *Gracilaria* to 300 Ma). Replicate projections were combined using AUC-weighted medians and masked to coastal grid cells. Binary suitability was defined using the sensitivity-specificity threshold. Latitudinal trends were summarized using the mean absolute latitude and latitudinal range of suitable cells. Minimum and maximum limits were defined using the 0.05 and 0.95 quantiles of the distribution of suitable cells to reduce sensitivity to outliers. Spatial continuity of environmentally suitable areas for each genus was quantified by calculating the number of contiguous patches of binary suitable habitat using the terra R package v1.7-71 (*93*). Temporal trends in global mean SST were extracted from the HadCM3 simulations and compared with the proportion of coastal grid cells classified as suitable for *Ahnfeltia* through time. Environmental novelty was assessed using MESS analyses.

Thermal niches were quantified using minimum volume ellipsoids (90%) based on mean annual SST and SST variability. Niche overlap was calculated using Jaccard and Sørensen indices. Further detailed information is described in Supplementary Methods S6.

## Data availability

*Ahnfeltia* genome assemblies were deposited on NCBI under PRJNA1447080. Gene models, annotations, and variant files were uploaded to the figshare under the doi link: https://doi.org/10.6084/m9.figshare.31859284. Raw data used for genome assemblies were deposited on NCBI under BioProjects PRJNA1446966.

## Code availability

All software and pipelines used in this study were executed according to the developers’ documentation and associated publications. Software versions and parameter settings are described in the Methods section. Default parameters were used unless otherwise specified.

## Acknowledgements

TM acknowledges partial support from the School of Environmental Sciences, University of East Anglia, Norwich Research Park, Norwich, UK.

## Funding

This study was supported by grants from the National Research Foundation of Korea (RS-2022-NR068987, RS-2022-NR070837, RS-2025-25434447), the Bio&Medical Technology Development Program of the National Research Foundation (NRF) funded by the Korean government (MSIT) (No. RS-2024-00411768), the Korea Institute of Marine Science and Technology Promotion (KIMST) funded to the Ministry of Oceans and Fisheries (RS-2025-02304428). AM was supported by FONDECYT Regular 1241697 and ANID BASAL FB210018.

## Authors’ contributions

HSY organized the study. HSY and TM supervised the project. HCK, DEB, MSC, AM, GIH, AVS, AN, KAM, BK, DK, YL, DL, GWS, and CAM collected samples and provided specimens. RM and JB generated molecular marker sequence data. HCK performed sequencing, assembled the reference genomes, and conducted comparative genomic, population genomic, and phylogenomic analyses. CHC, JJ, YL, and LG contributed to data interpretation, and DL, CJ and CVO supervised population genomic analyses. HCK and SC performed divergence time estimation. HCK and JHK examined agar properties. RD conducted palaeogeographic niche modelling under the supervision of ES. AF, PV, and DL contributed palaeoclimate modelling resources and methodology. CAM, GWS, AVS, JB, DEB, MSC, and AM confirmed *Ahnfeltia* occurrence and ecological information. SX provided insights into the palaeobiology of red algae. HCK, TM, and HSY wrote the manuscript. All authors reviewed and approved the final version.

## Ethics declarations

### Competing interests

The authors declare no competing interests.

## Notes

### Competing Interest Statement

The authors have declared no competing interest.

