## Supplementary Information for "The genomic basis of evolutionary stasis in the 500-million-year-old red seaweed genus *Ahnfeltia*"

**Supplementary information contents:**

### Supplementary Methods S1: Construction of *Ahnfeltia* genomes

#### Contaminant evaluation and genome size estimation

As *Ahnfeltia* samples were collected from natural habitats, it was difficult to completely eliminate bacterial contamination prior to sequencing. Therefore, contaminant reads were identified and removed using Kraken2 (*1*) and the BlobTools pipeline (*2*) before genome size estimation. Short reads from selected individuals for nuclear genome assembly were preliminarily assembled using SPAdes v3.15.2 (*3*). The taxonomy of assembled contigs was evaluated using DIAMOND BLASTX (*4*). Short reads were then mapped back to the assembled contigs to obtain coverage information for each contig using BWA v0.7.17 (*5*) and SAMtools v1.7 (*6*). Based on the DIAMOND results and BAM files containing mapping information, BlobTools was used to generate a taxonomic assignment table. From these results, contigs assigned to Rhodophyta were selected as candidate sequences. Raw reads were subsequently remapped to the Rhodophyta candidate contigs and sorted using BWA v0.7.17 and SAMtools v1.7. Using the sorted reads, *k*-mer base genome size estimation was performed with Jellyfish (*7*) and visualized using GenomeScope 2.0 (*8*).

#### *De novo* nuclear genome assembly and assessment

*De novo* nuclear genome assembly was conducted using Falcon (*9*) as well as NextDenovo (*10*). For Falcon assembly, most default parameters were used, yet the following parameters were modified: pa_daligner_option=-e.7 -l2000 -k14 -h480 -w8 -s100, falcon_sense_option=--output-multi --min-idt 0.70 --min-cov 2 --max-n-read 500 in pre-assembly step, ovlp_daligner_option=-e.93 -k18 -h1024 -w6 -s100 -l6000, ovlp_HPCdaligner_option=-v -B128 -M24 in pread overlapping step, overlap_filtering_setting=--min-cov 2 --max-cov 500 --max-diff 200, fc_ovlp_to_graph_option= --min-len 1000, length_cutoff_pr=1000. The assembled genomes from Falcon were polished using Falcon-Unzip in a series of the process. The NextDenovo assembly was conducted with default parameters. The primary assembly results from both tools were compared and improved using Geneious prime 2022.2.2 (https://www.geneious.com). Bacterial contigs were also identified and removed using BLASTn and BLASTp (*11*) by comparing all contigs to the NCBI nt and nr databases. The remaining genomes were polished using pilon v1.24 (*12*) after aligning short reads to genomes using BWA v0.7.17 (*5*) and Samtools v1.7 (*6*). In addition, duplicated contigs in polished genomes were removed using Purge_haplotigs v1.1.2 (*13*) and Purge_dups v1.2.3 (*14*) after aligning long-reads to the genomes using Minimap2 v2.17 (*15*). The completeness of the final genomes was checked with their read coverage and alignments by using WGSCoveragePlotter (*16*; fig. S33).

**Detecting transposable elements (TEs) and repeat masking**

As there is no robust TE library for red algae, *de novo* repeat libraries for *Ahnfeltia* were constructed first. To make the repeat library, TEs were identified from not only *Ahnfeltia* genomes but also published red algal genomes using RepeatModeler v2.0.3 (*17*) with -LTRStruct option. There were still many unclassified repeats from the result of RepeatModeler. Thus, re-classification of them was carried out using DeepTE (*18*). Unknown repeats identified by RepeatModeler were manually reclassified using DeepTE, and the results were integrated using a custom Python script. *Ahnfeltia* genomes were masked using RepeatMasker v4.1.2 (*19*) with the newly constructed repeat library. Masked genomes were used for the next gene prediction step.

**Gene prediction**

The transcripts were trimmed using Trimmomatic v0.39 (*20*) and aligned to the final *Ahnfeltia* genomes using STAR v2.7.10a with default parameters (*21*). Gene prediction was conducted with three pipelines based on *ab initio* gene prediction, protein, and transcript alignments. For *ab initio* gene prediction, BRAKER2 was employed with --softmasking option (*22, 23*). GeMoMa v1.7.1 was used for protein alignment (*24*). Proteins from published high-quality genomes of the red algae (*Chondrus crispus, Cyanidioschyzon merolae, Gracilariopsis chorda, Porpyridium purpureum, Pyropia yezoensis,* and *Porphyra umbilicalis*) were aligned and compared to *Ahnfeltia* genomes (references are listed in table S5). PASA (*25*) was utilized for spliced alignments of transcripts based on the result of Trinity-v2.8.5 with --genome_guided_bam option (*26*). Finally, a consensus gene model ended up being constructed using EVidenceModeler v1.1.1 (*27*). The completeness of genomes and gene models was assessed using BUSCO v4.1.2 with eukaryote odb9 and eukaryote odb10 (*28*).

**Functional annotation**

Predicted genes were annotated by using MMseqs2 (*29*) against locally installed NCBI nr database (February 20th, 2023) with the cutoff e-value 1e-05, BlastP (*11*) against UniProt database (*30*), eggNOG-Mapper v2.1.3 (*31*) with the eggNOG database (*32*), InterProScan v5.52-86.0 (*33*), and KEGG Automatic Annotation Server (KAAS) (*34*).

### Supplementary Methods S2: Divergence time estimation for *Ahnfeltia*

To obtain robust divergence time estimation for *Ahnfeltia* lineage, Bayesian MCMCtree (PAML packagev4.9j) was performed (*35, 36*). MCMCtree is a Bayesian divergence time estimation method that uses an approximate likelihood based on maximum likelihood estimates to improve computational efficiency (*37*). Due to the limited availability of red algal genome sequences, nuclear single-copy BUSCO genes were used for divergence time estimation as conducted in the phylogenomic analysis.

Three fossil calibration points were adopted from Yang et al. (*38*): the *Bangiomorpha* fossil (1,198 Ma; 95% CI: 1,174-1,222 Ma), the Doushantuo Formation fossil (593 Ma; 95% CI: 551-635 Ma), and the split of Sporolithales (133 Ma; 95% CI: 130-136 Ma). Because genome sequences from Sporolithales are currently unavailable, an additional analysis using only two calibration points excluding "Sporolithales split" was performed to evaluate the robustness of divergence time estimation.

Given that *Ahnfeltia* species exhibit relatively low substitution rates compared to other red algae in the nuclear ML phylogeny (Fig. 2A), divergence time estimation based solely on their substitution rates could be biased toward recent divergence. Although low substitution rates can suggest recent diversification, fossil and phylogenetic evidence of red algae indicate that *Ahnfeltia* lineages represent an ancient lineage within Florideophyceae (*38-40*). Therefore, divergence time estimates based purely on molecular rates may underestimate the true divergence time.

To address this issue, two molecular clock models were tested: an independent rates clock model and a correlated rates clock model (*37*). Under the independent clock model, evolutionary rate variation among *Ahnfeltia* lineages was estimated independently. Under the correlated clock model, rate variation was estimated in the context of broader red algal evolutionary history. Each MCMC chain was run for at least five million generations. After running, convergence and parameter mixing were evaluated using Tracer v1.7.1 (*41*). All parameters showed effective sample size (ESS) values greater than 200, indicating adequate sampling and convergence. The first 20% of generations were discarded as burn-in.

Evolutionary rates (number of substitutions per site per unit time) and lineage accumulation patterns were inferred from the time-calibrated phylogenies obtained from MCMCtree. Posterior mean trees were generated from the MCMC output and used as representative ultrametric trees for downstream analyses. Lineage-through-time (LTT) plots were generated using R package ape to compare diversification patterns between trees (*42*).

Evolutionary rates per branch were estimated as the ratio of branch lengths from phylogram and chronogram trees. Rate variation among branches was visualized using phytools (*43*) and fields (*44*).

### Supplementary Methods S3: Comparative genome analysis of *Ahnfeltia* genomes

#### Comparison of gene inventory

The difference of gene inventories from five *Ahnfeltia* genomes were compared using OrthoVenn3 (*45*). To compare how conserved the genomes are, the ratio singletons of *Ahnfeltia* genomes were compared with four *Gracilaria* genomes (*46*). In addition, divergence between *Ahnfeltia* and *Gracilaria* was estimated based on dS values calculated using CODEML implemented in PAML v4.9j (*35*). Their functional differences of genes were compared based on the result of from eggNOG, the Clusters of Orthologous Groups (COG) category (*31, 32*). The gene synteny of the genomes was analyzed using the python version of MCscan (*47*). For general description of genomic features of *Ahnfeltia*, gene density, transposable element density, GC, AT content, and Kimura value were portraited using CIRCOS (*48*) with reference to codes on github.com/GoreLab/Sorghum-HapMap and BEDTools v2.25.0 (*49*).

Orthologous gene families (OGFs) were clustered using OrthoFinder V2.5.2 with -M msa -S blast options (*50*). Protein sequences from four *Ahnfeltia* and the sequences from published genomes of the red algae, *Rhodelphis marinus,* and the green lineage (*Arabidopsis thaliana*, *Chlamydomonas reinhardtii*, *Oryza sativa*, *Ostreococcus tauri*, *Physcomitrella patens*) were analyzed for the OGFs analysis (*51-54*). The gain and loss of genes were estimated using Count with the Dollo parsimony principle (*55*). Gain and loss were sorted using extract_dollop_output_sequences_v2-fast.pl script (github.com/guyleonard/orthomcl_tools). Gain genes were annotated using NCBI nr database and InterProScan v5.52-86.0 (*33*).

To evaluate evolutionary pressure on orthologous genes, non-synonymous/synonymous mutation rate (dN/dS) of each orthologous gene was calculated. Multiple codon alignment of comparable orthologous genes (6,897) between *Ahnfeltia* species was conducted using MAFFT v7.310 (*56*). PAL2NAL v14 (*57*) was used for CODEML program in PAML v4.9j (*35*) with -output paml -nogap option. IQTREE v1.6.8 (*58*) was used to construct a tree file. After writing a ctl file, dN/dS were calculated using CODEML.

#### Verification incomparable regions between *A. plicata* and *A. borealis/A. fastigiata* clade

The highly mutated regions between *A. plicata* and *A. borealis*/*A. fastigiata* clade appeared because they were completely diverged. To identify these incomparable regions between them, unmapped regions were investigated after mapping WGS data of individuals of one species to the other *Ahnfeltia* species genome (*e.g. A. borealis* and *A. fastigiata* individuals to *A. plicata* genome). GATK results including SNP and indel information described in the variant calling were used. We manually categorized variant regions that were present in one species but absent in individuals of the other species using a Python script. Additionally, we confirmed the presence of two inverted regions by comparing entire scaffolds using LASTZ v1.02 (*59*) involved in Geneious prime 2022.2.2 (https://www.geneious.com). Gene Ontology (GO) analysis was also conducted to estimate the function profiling of genes in differentiated regions. gprofiler2 R package (*60*) was used for the GO analysis. As gprofiler2 requires a GMT file, GMT files were established based on the results from InterProScan v5.52-86.0 (*33*) and the go.obo file from the GENEONTOLOGY web site (http://geneontology.org/docs/download-ontology/).

**TE insertion in *A. fastigiata***

In the divergence of *A. fastigiata* from other *Ahnfeltia* species, TE was identified as the important factor generating the compartment between *A. fastigata* and *A. borealis*. TE divergence based on Kimura-distance was calculated using a cat file from RepeatMasker v4.1.2 (*19*). The insertion sites in intergenic and intronic regions of TEs were pinpointed using gff files from RepeatMasker and genemodel. To facilitate a clear comparison among *Ahnfeltia* species, only comparable orthologous genes (6,897) were analyzed. Lastly, GO analysis was conducted to infer which function is related to genes found in TE insertion regions.

Kimura divergence values obtained from the transposable element (TE) landscape were interpreted under a neutral substitution model, where sequence divergence (K) accumulates as a function of evolutionary time according to (K = 2xμxT), with (K) representing Kimura two-parameter sequence divergence, (μ) the neutral substitution rate per site per year, and (T) the divergence time (*61*). We adopted a mean evolutionary rate of ($1.13 \times 10^{-8}$) substitutions per site per year estimated from an MCMC-based molecular clock analysis, assuming a generation time of one year. Under this model, the estimated divergence time between the two clades (~148 Ma) corresponds to a Kimura divergence of approximately 3.3%. This calibration was used to provide an approximate temporal interpretation of the TE divergence landscape.

However, sequence divergence measures are known to become increasingly affected by substitution saturation and multiple substitutions at the same site as divergence increases, resulting in a loss of linearity between sequence divergence and time (*62, 63*). Consequently, the relationship between Kimura divergence and evolutionary time is expected to remain approximately linear only within the lower divergence range. For this reason, divergence times inferred from Kimura values were interpreted only for relatively low divergence levels (*e.g.*, ≤10-15% Kimura), whereas higher divergence values were considered to reflect ancient TE insertions and long-term sequence persistence rather than a direct linear relationship with evolutionary time.

**Comparison of gene contents related to DNA repair systems**

We observed that *Ahnfeltia* lineages exhibit low substitution rates and conserved genomic evolution. To test if this evolutionary stasis may be associated with duplication of DNA repair systems, we compared gene content related to DNA repair pathways across five *Ahnfeltia* genomes and other seaweeds (11 Rhodophyta, two Phaeophyceae, and two Chlorophyta). DNA repair genes were categorized according to KEGG pathway classifications, including approximately 360 genes. KEGG Orthology (KO) numbers for the seaweed genomes were obtained using the KEGG Automatic Annotation Server (KAAS) (*34*).

**Supplementary Methods S4: Characterization of *Ahnfeltia* agar component**

The agar component of *A. plicata* Chile and Wales, UK and *A. fastigiata* Korea and Oregon, USA were characterized to compare a populational difference. *Gelidium elegans* were selected to compare the agar character of *Ahnfeltia* with red algae in Rhodyemeniophycidae. Attached eukaryote contaminants on fresh samples were manually removed with forceps as many as possible and washed with sterilized water. Samples were frozen using liquid nitrogen and lyophilized in a row (Ilshinbio, Gyeonggi-do, South Korea). Dried samples were fragmentized using the automill (Tokken, Japan). About 100 mg of each powder was put into a 20 mL vial. After adding about 10 mL distilled water to the vial, the samples were pre-treated by an ultrasonicator (Hwashin Instrument, South Korea) keeping 50 °C for 30 minutes. The agar from each sample was extracted by heating them using a heat bath (Hansol Tech, South Korea) and keeping them at 90 °C for 3 hours. About 10 mL of extracted agar product was collected in a 15 mL conical tube (SPL Life Sciences, South Korea). The extracted agar was lyophilized once again. For the sulfate quantification, a conditioning reagent was prepared; the conditioning reagent included glycerol 25 mL, hydrochloric acid 32% 15 mL, 2-propanol 50 mL (Merck, USA), 5M sodium chloride 12.5 mL (Merck, USA), and it was adjusted to 500 mL of the final volume by adding distilled water. A reacting solution was made of 12.5 µl of the conditioning reagent, 25 µl of 3% barium dichloride solution, and 87.5 µl of distilled water. The powder of the extracted agar was melted into distilled water with a 1 mg/mL concentration. 125 µl of the liquid agar was reacted with the reacting solution on a 96-well microplate (SPL Life Sciences, South Korea) by shaking them for more than a minute. A calibration curve was based on the result of the reaction between four concentrations (1 mg/mL, 500 µg/mL, 250 µg/mL, and 50 µg/mL) of potassium sulfate solution with the reacting solution. The absorbance of the reacted suspension was measured using a spectrophotometer at 420 nm wavelength (BMG LABTECH, Germany).

### Supplementary Methods S5: Analyses for populational divergence

#### Evaluation of *Ahnfeltia* variant datasets

Because *Ahnfeltia* exhibits a heteromorphic triphasic life cycle and carposporophytes (2n) are form the surfaces of gametophytes (n), ploidy levels of all samples were assessed prior to downstream population genomic analyses. To evaluate ploidy and data quality, we examined the distribution of homozygous and heterozygous genotype calls as well as per-site read depth across individuals. The proportions of heterozygous sites and depth distributions were used to distinguish whether *Ahnfeltia* samples were solely from haploid stages or from potential mixed stages.

*k*-mer frequency distributions were additionally analyzed to infer genome characteristics and confirm ploidy status. *k*-mer counting was performed using Jellyfish (*7*), and *k*-mer spectra were visualized using GenomeScope to estimate heterozygosity and ploidy peaks (*8*). To further quantify within-sample genetic diversity and assess clonality or multiplicity of infection–like patterns, the within-sample fixation index (FWS) was calculated using moimix (*64*). FWS values approaching 1 indicate highly homozygous (haploid-like or clonal) samples, whereas lower values suggest elevated heterozygosity consistent with diploidy.

#### Population structures and relationship

Principal component analysis (PCA) and admixture analysis were performed using the LEA R package (*65*) to assess population relationships. To obtain robust results, two different SNP datasets were analyzed: (i) a dataset including all SNPs shared across all individuals (*i.e.,* without missing data), and (ii) a linkage disequilibrium (LD)-pruned dataset generated using PLINK v1.90b6.24 (*66*) with the parameter --indep-pairwise 40 5 0.5 to reduce the effect of non-randomly associated SNPs. For admixture analysis, the *snmf* function of LEA was run with 20 repetitions and K values ranging from 1 to 14 (*65*). In addition, population relationships were further investigated using a phylogenetic network constructed from SNP data in SplitsTree6 (*67*). Gene flow was further evaluated using the Fbranch statistic implemented in Dsuite, which maps excess allele sharing (based on Patterson's D statistics) onto branches of the phylogeny (*68*).

#### Assess of local adaptation of *Ahnfeltia* populations

To identify differentiated genomic regions and selection signals associated with local adaptation in *Ahnfeltia* populations, fixation index ($F_{ST}$) and genetic divergence between two populations (Dxy) were calculated based on 20 kb sliding windows and 10kb steps. Weir and Cockerham's $F_{ST}$ was calculated using vcftools v0.1.17 (*69*) and Dxy was calculated using the genomics_general pipline (github.com/simonhmartin/genomics_general). All manhattan plots were generated using qqman R package (*70*). To investigate genetic functions of highly divergent regions, Gene Ontology (GO) enrichment analysis was conducted using gprofiler2 in R (*60*). Gene annotations were obtained from InterProScan (*33*), and GO terms were retrieved from the Gene Ontology database (http://geneontology.org/docs/download-ontology/). To investigate the selective pressure on protein gene sets at the coding level, dN/dS ratio between two *Ahnfeltia* populations were also calculated. Population-specific nuclear genomes were updated by incorporating SNP information into the reference genome using bcftools (*71*). Coding sequences (CDSs) were extracted using GFF annotation files and gffread (*72*). Pairwsie dN/dS values were then calculated using KaKs_Calculator 2.0 (*73*).

**Genetic diversity and homozygosity of *Ahnfeltia* populations**

Nucleotide diversity ($\pi$) within species and populations was calculated per 20 kb nonoverlapping windows using vcftools v0.1.17 with --window-pi 20000 and --keep options (*69*). To calculate homozygosity and inbreeding levels in *Ahnfeltia* populations, Individual inbreeding coefficients (F) and observed heterozygosity (Ho) were calculated using plink v1.90b6.24 with --het and --hardy option, respectively (*66*). Wright's inbreeding coefficient within populations ($F_{IS}$) were calculated following 1-Ho/He, where He represents expected heterozygosity under Hardy-Weinberg equilibrium. Because many loci were monomorphic within populations (both Ho and He were 0), $F_{IS}$ values were undefined at those sites due to division by zero. If all these sites are ignored, the very small number of valid $F_{IS}$ values can represent a whole window and misleadingly suggest outbreeding, which seemed unreasonable. Thus, we treated monomorphic sites as fully homozygous ($F_{IS}$ = 1) by assuming that extremely low He and Ho = 0 (as we observed).  Window-based $F_{IS}$ values were then calculated as the mean $F_{IS}$ across SNPs within each 20-kb window. Runs of homozygosity (ROH) were identified using PLINK with --homozyg --homozyg-kb 20 --homozyg-snp 20 --homozyg-density 5 --homozyg-gap 20 --homozyg-window-snp 50 --homozyg-window-het 0 --homozyg-window-threshold 0.05 parameters. This result was also compared by allowing one heterozygous individual. The genomic inbreeding coefficient based on ROH (FROH) was calculated as the proportion of the genome contained within ROH for each individual (total ROH length divided by total autosomal genome length).

**Evaluation of isolation-by-distance (IBD)**

To evaluate the impact of geographic barriers on *Ahnfeltia* populations, isolation-by-distance (IBD) was assessed by examining the relationship between genetic differentiation ($F_{ST}$) and geographic distance. Pairwise weir and Cockerham's $F_{ST}$ was calculated using vcftools v0.1.17 (*69*). Geographic distances were calculated as great-circle distances (km) based on latitude and longitude coordinates of each sampling site. The relationship between geographic and genetic distances was assessed using Pearson's correlation test, and linear regression was used for visualization.

To place the observed levels of genetic divergence in a broader context, we compared our $F_{ST}$ estimates with values reported for other seaweeds in Durrant et al. (*74*) since there are no other genome-scale population studies on red seaweeds to date. Although methodological differences and marker types limit direct quantitative comparisons, this comparison provides a general framework for interpreting the magnitude of divergence observed in *Ahnfeltia*. A more rigorous cross-species comparison will require additional genome-wide population studies in red algae and other seaweeds in further studies.

**Linkage Disequilibrium (LD) decay**

LD decay of each *Ahnfeltia* population was calculated to estimate the level of recombination within populations and compare evolutionary history of them. Mean $r^2$ was calculated using PopLDdecay v3.41 (*75*) with a default option.

**Estimation of historical effective population size (Ne) trajectories**

To infer historical changes in effective population size ($N_e$), we applied the stairway plot v2 (*76*) based on the folded site frequency spectrum (SFS). The folded SFS was generated from high-quality biallelic SNPs without missing data. The folded SFS was used as input, and multiple independent runs were performed to ensure convergence. Bootstrapping (200 replicates) was applied to estimate confidence intervals around $N_e$ trajectories. Mutation rate ($\mu$) and generation time (g) were specified to convert scaled time and population size estimates into absolute values. Because mutation rates are not well characterized in *Ahnfeltia*, we used an estimated mutation rate of $1x10^{-9}$ per site per generation. Generation time was assumed to be three years based on Brodie et al. (*77*). The resulting $N_e$ trajectories were visualized as changes in effective population size through time, with confidence intervals derived from bootstrap replicates.

### Supplementary Method S6: Ecological Niche Modelling of *Ahnfeltia* and *Gracilaria* through the Phanerozoic

#### Verification of *Ahnfeltia* occurrences

Although 10 *Ahnfeltia* species are currently listed in AlgaeBase (*78*), many of them have not yet been evaluated using molecular data. To date, only three species (*i.e. A. plicata, A. borealis*, and *A. fastigiata*) are supported by genetic evidence (*79-81*), based on worldwide sampling. These records are also reflected in the Barcode of Life Data System (*82*), which we used as a reference for subsequent analyses.

We re-examined historical herbarium specimens and added the occurrence information of newly-collected material. In particular, we carefully checked records from tropical regions, including the Galápagos Archipelago, because these occurrences were important for ecological niche modeling. We found that many of the putative *Ahnfeltia* specimens belonged to Gigartinales, such as *Gymnogongrus* sp. In addition, two type species (*A. svensonii* and *A. gigartinoides*) were Gigartinales species (table S2 and Fig. S53).

These results suggest that the remaining listed *Ahnfeltia* species need further taxonomic confirmation, especially those originally identified only based on morphology. We cannot draw a definitive conclusion about their actual occurrence without additional molecular evidence. Therefore, in this study, we used only *Ahnfeltia* occurrence records supported by reliable morphological and/or molecular information to keep our dataset conservative.

Occurrence records for the red algal genera *Ahnfeltia* and *Gracilaria* were compiled from the Barcode of Life Data System (*82*), which contains records based on genetically identified specimens (table S1). In addition, the occurrence of the confirmed old herbarium specimens of *Ahnfeltia* were added (table S2). Records were then cleaned using the CoordinateCleaner v3.0.1 R package (*83*), and entries with missing or invalid coordinates, duplicate records, or locations falling on land were excluded. We also included an *Ahnfeltia* dataset from the Global Biodiversity Information Facility (GBIF; GBIF.org, 17 September 2025, Occurrence Download, doi:10.15468/dl.5a4pgw), which was not genetically verified, to evaluate how incorporating additional but less certain records might affect our projections. The number of occurrence records decreased from 342 to 39 for *Ahnfeltia* and from 419 to 81 for *Gracilaria* after quality and spatial filtering (Supplementary Table 2).

#### Palaeoclimate Data

We used a recently updated version of the UK Met Office's HadCM3 palaeoclimate model (*84*), with a resolution of 3.75° longitude (~300 km in the tropics) and 2.5° latitude for both the atmosphere and the ocean, following Valdes et al. (*84*) and Valdes et al. (*85*), with additional model development described in Judd et al. (*86*). Environmental predictors were derived from two palaeogeographic frameworks: (1) the Getech palaeogeographies (286.8 Ma to Present), and (2) the Scotese climate reconstructions (*87*). The Scotese framework included two variants to represent uncertainty in Phanerozoic $CO_2$ evolution: (i) a temperature-based simulation ("Scotese_Temperature"), in which the evolution of atmospheric CO2 concentration was chosen such that the modelled global mean surface temperatures follow those of Scotese et al. (*88*) derived from geological temperature proxies (identical to simulation 'Scotese_08' in Judd et al. (*86*)), and (ii) a $CO_2$-based simulation ("Scotese_Foster-$CO_2$"), in which the evolution of atmospheric $CO_2$ followed the proxy-based trajectory of Foster et al. (*89*) (identical to simulation 'Scotese_07' in Judd et al. (*86*)). A comparable $CO_2$-derived reconstruction based on the Getech palaeogeographies ("Getech_Foster-$CO_2$") was also used, applying the same Foster et al. (*89*) trajectory. For all simulations, the native climate data were bilinearly interpolated to a uniform grid with an approximate resolution of 100 km in the tropics.

**Environmental variables**

From each palaeoclimate reconstruction, monthly sea surface temperature (SST), salinity, and photosynthetically active radiation (PAR) at 5 m depth were extracted to represent the intertidal and subtidal habitats of *Ahnfeltia* and *Gracilaria*. For each climate variable (Supplementary Table 1), the mean, minimum, maximum, seasonal range, and standard deviation (SD) were calculated from twelve monthly layers to capture both average conditions and seasonal variability. To minimize multicollinearity, pairwise correlations were assessed using the Variance Inflation Factor (VIF) approach implemented in the usdm v2.1-7 R package (*90*). Climatic variables with VIF values greater than 10 were iteratively excluded until all retained predictors met this threshold. The final set of predictors for both genera included salinity (SD and maximum), SST (range and maximum), and PAR (SD and minimum).

**Background sampling**

The calibration region was defined by buffering all occurrence localities (*91*). The buffer radius was set to the mean pairwise geographic distance among records, following previous studies that used inter-occurrence distances to approximate accessible areas (*92-95*). The buffered regions represented the accessible area for each genus and the restricted background sampling to realistically reachable locations (*91, 96, 97*). All environmental layers were masked to these genus-specific buffered areas to define the extent of background sampling. A target-group background approach was then used to account for spatial variation in sampling effort (*97, 98*). Material records for Florideophyceae were obtained from GBIF (*99*) to approximate the spatial distribution of collection effort and generate a sampling-bias surface. This surface was log-transformed and normalized to create a probability layer from which background points were drawn in proportion to sampling intensity, thereby weighting selection toward coastal regions with greater sampling effort. To reduce the effects of spatial clustering and sampling bias (*100*), presence records were then spatially thinned using the spThin R v0.2.0 package (*101*). The thinning threshold was set to the first distance at which Moran's I of scaled environmental variables crossed zero, computed with the ncf v1.3-2 (*102*). Background points were subsequently thinned using the same distance threshold, with the final background set containing ten times the number of retained presences (*103*).

**Model calibration and evaluation**

ENMs were fitted using the Maxent algorithm, a maximum entropy method (*104, 105*) that has demonstrated high predictive accuracy in palaeo-ENM contexts (*106, 107*). Maxent is particularly well-suited to small, presence-only datasets that are spatially biased, such as those used in this study, and addresses the problem of unknown true absences by sampling the environmental background (*108, 109*).

Limiting model complexity is critical for estimating the fundamental niche and projecting models through deep time because overly complex models tend to overfit present-day conditions, reduce temporal transferability, and perform poorly when extrapolated to novel or climatically dissimilar environments (*107, 110*). Accordingly, we restricted models to linear and quadratic feature classes and tested 21 regularization multipliers, ranging from 1 to 3 in 0.1 increments, to identify the optimal level of penalization. Model calibration and evaluation were performed using the blockCV v3.2-0 R package (*111*), applying both environmental clustering and spatial block partitioning as complementary four-fold cross-validation approaches. These approaches maintain spatial independence between training and testing data, thereby reducing inflated accuracy estimates (*112, 113*). Four folds were used to prevent over-fragmented partitions in the small occurrence datasets.

The resulting partitions were supplied to ENMeval v2.0 (*114*) for model tuning and validation. Candidate model performance was evaluated using the small sample corrected Akaike Information Criterion (AICc), spatial validation area under the receiver operating characteristic curve (AUC), and the Continuous Boyce Index (*115*). The best-performing model for each genus was defined as the configuration with the lowest AICc value. When multiple models had $\Delta$AICc < 2, the model with the lower regularization multiplier was selected to favor parsimony (*116*). The selected configuration was then rerun ten times using alternative random background samples to account for stochastic variation in background selection.

**Model projection through the Phanerozoic**

For each genus, the ten final model replicates were projected onto palaeoclimate time slices spanning 108 intervals for *Ahnfeltia* (0 to 505 Ma) and 60 intervals for *Gracilaria* (0 to 300 Ma). To visualize broad temporal patterns, we present a subset of projections at approximately 100 Mya intervals: 0, 102.6, 201.3, 301.3, 400, and 505 Mya within the Scotese framework, and the closest available time slices (0, 102.6, 201.3, and 286.8 Mya) within the Getech framework. Projections were generated on the cloglog scale, representing estimated probabilities of occurrence ranging from 0 to 1 (*117*). For each time slice, suitability maps from the ten replicates were combined using a weighted median approach to produce a single final map, with weights based on each replicate's mean environmental-fold AUC. To restrict projections to potential coastal regions accessible to *Ahnfeltia* and *Gracilaria*, which are intertidal and upper subtidal taxa (*118-120*), outputs were masked to include only grid cells adjacent to land. Continuous suitability values were converted to binary presence-absence maps using the sensitivity-specificity threshold, which maximizes the sum of sensitivity and specificity (*121*). This threshold balances omission and commission error and is widely used in species distribution modelling (*122*). The mean absolute latitude and latitudinal range of projected presence cells at each time slice were then calculated to track latitudinal changes in potential suitability through time. Minimum and maximum limits were defined using the 0.05 and 0.95 quantiles of the distribution of suitable cells to reduce sensitivity to outliers. Spatial continuity of environmentally-suitable areas for each genus was quantified by calculating the number of contiguous patches of binary suitable habitat using the terra R package v1.7-71 (*123*).

**Novel environments**

ENM projections may encounter novel environmental conditions that were not present in the model calibration region, which can affect model accuracy (*124*). To evaluate model extrapolation through deep time, multivariate environmental similarity surface (MESS) analyses were conducted for each geological time slice using the mess function in the dismo v1.3-16 R package (*125*). For each slice, we calculated the proportion of climatically suitable area with negative MESS values to quantify the extent of environmental novelty relative to modern training conditions. Negative MESS values indicate environmental conditions outside the range of those used to train the model, and thus represent areas of extrapolation (*126*). In addition, we performed a sensitivity analysis in which all grid cells with negative MESS values were excluded from the suitability maps prior to summarizing latitudinal patterns, allowing us to assess how our main results were influenced by the inclusion of novel suitable regions.

**Limitations and caveats of projections**

Before interpreting the results, it is important to recognize that the palaeoclimatic projections presented here inevitably involve uncertainties. ENMs estimate environmental suitability rather than realized distribution, and projections should therefore be interpreted as highlighting regions with broadly favorable conditions, and not precise reconstructions of occupied range. Additional environmental factors, such as pH, nutrient availability, and substrate type also influence the suitability of a region for these intertidal and subtidal macroalgae (*127*). For example, *Ahnfeltia* species are closely associated with rocky-substrate habitats (*120*), but this variable could not be included because such data are unavailable in global palaeoclimate reconstructions.

Our analyses were conducted at the genus rather than species level, which can obscure meaningful variation in environmental tolerances among constituent taxa (*128, 129*). Species-level modelling was constrained by limited occurrence sampling and turnover across Phanerozoic timescales, so we employed genus-level ENMs to provide conservative estimates of the broader climatic envelopes occupied by these lineages. However, genera are not biologically coherent or evolutionarily functional units, but rather arbitrarily circumscribed, often paraphyletic or polyphyletic groupings of species assembled largely for taxonomic convenience (*129*). Environmental tolerance can vary substantially among species within the same genus (*130*) and thus modelling the niche at a higher taxonomic level may produce inflated or misleading representations of true ecological constraints.

These concerns are particularly relevant when projecting a genus-level estimate of climatic suitability back hundreds of millions of years. Whilst this approach may capture large-scale patterns, it also compounds the broader challenges of projecting ENMs into deep time, where stable ecological relationships cannot be assumed due to evolutionary change over macroevolutionary timescales. For instance, ecological niches may shift during or following speciation events or along lineages, violating the assumption of niche conservatism that underpins ENM transferability (*131, 132*). Consequently, even if genus-level models capture broad climatic envelopes, modern niches likely represent only the most recent ecological expressions of lineages that have undergone repeated adaptive transformations throughout the Phanerozoic. These uncertainties are further exacerbated by the magnitude of climatic change that occurred throughout the Phanerozoic (*110*).

Finally, palaeoclimate reconstructions may include regions with non-analogue conditions, where extrapolation beyond the present-day calibration domain introduces additional uncertainty (*133, 134*). The generated maps should therefore be regarded as hypotheses of potential environmental suitability rather than definitive reconstructions of their past geographic distributions. The climate model simulations themselves are associated with uncertainty. Although we have explored climatic uncertainty associated with the uncertainty in $CO_2$ ("Scotese_Temperature" versus "Scotese_Foster-$CO_2$") and palaeogeography ("Scotese_Foster-$CO_2$" versus "Getech_Foster-$CO_2$"), there is still uncertainty associated with the model itself. Future work could explore this further by comparison with similar simulations carried out with another climate model (*e.g.* Li et al (*135*)).

**Climatic hypervolume**

To compare the thermal niches of *Gracilaria* and *Ahnfeltia*, which are hypothesized to be ecologically distinct, we used multivariate hypervolume modelling based on two key temperature variables: mean annual SST and its variability (SD of SST). We focused on temperature because differences in thermal tolerance likely underpin the contrasting biogeographic distributions of *Gracilaria* and *Ahnfeltia (79, 119, 120)*. For each genus, modern baseline marine temperature variables (Supplementary Table 1) were extracted at all occurrence locations retained after data cleaning. These two variables were used directly as the environmental axes for hypervolume construction. For each genus, a minimum volume ellipsoid (MVE) was fitted to the environmental points to delineate the smallest convex ellipsoid encompassing 90% of the most central observations (*136*). This approach aims to estimate the realized thermal niche of each genus under the assumption that ecological niches are approximately convex in environmental space, for which ellipsoidal representations are an appropriate approximation (*137, 138*). Specifically, a MVE was selected because it performs robustly with small sample sizes (*139*). Thermal niche overlap was quantified using Jaccard and Sørensen similarity indices, together with measures of shared and unique hypervolume area for each genus.

### Supplementary Text S1: Assessment of ploidy in *Ahnfeltia* individuals

For population analyses, it is important to determine the ploidy of the target organisms, as many analyses and their interpretations differ depending on ploidy level (*140, 141*). *Ahnfeltia* has a heteromorphic triphasic life cycle as do other florideophycean algae. In natural populations, the conspicuous thalli typically represent dioecious haploid gametophytes (n). Spermatia in male gametophytes produce sperm (n) that fertilize carpogonia on female gametophytes, after which diploid cells develop in the upper cortical layers and give rise to a globose diploid carposporophyte (2n). Carposporophytes release diploid carpospores (2n), which develop into a diploid crustose tetrasporophyte (2n). Meiosis within the tetrasporophyte produces haploid tetraspores (n), which subsequently develop into new gametophytes, completing the life cycle (*118, 142*).

In this study, erect gametophytes were re-sequenced for population analyses, and visible diploid lump tissues were carefully excluded during sampling. Therefore, only homozygous variants were expected in the dataset. However, heterozygous variants were consistently detected in all individuals, and their frequency exceeded expectations under minor contamination from diploid tissues (fig. S34 to S37, table S13).

To investigate this unexpected signal, we assessed ploidy using multiple complementary approaches. First, ploidy inference was performed using momix (*64*). Second, k-mer frequency distributions were examined using jellyfish and GenomeScope (*7, 8*). Third, we evaluated allele depth ratios at heterozygous sites. In strictly haploid individuals, alternative allele depth ratios are expected to cluster near 0 or 1, whereas diploid individuals should show a peak near 0.5. If low-level diploid contamination were present within predominantly haploid tissues, intermediate peaks around 0.25 and 0.75 would be expected.

Across individuals, results did not show clear clustering consistent with exclusively haploid signals. Instead, many individuals exhibited patterns suggestive of diploidy or mixed ploidy states; however, the allele frequency distributions were broadly dispersed rather than forming well-defined peaks (fig. S2 and S38). In this study, stringent filtering was applied (described in methods, main text) to minimize potential technical issues; however, the heterozygous signal persisted across coding and non-coding regions and was consistently observed across independent samples having similar read depth (fig. S39 to S41).

To further clarify the ploidy issue, we examined the difference between lump (putative 2n) and clean tip (putative n) of female gametophytes once again by separating the expected haploid and diploid areas and re-sequencing. In this experiment, a small number of heterozygous variants were still detected in branches of female gametophytes and read depth distributions did not show clear clustering patterns (fig. S42). While technical artifacts cannot be completely excluded, the repeated detection of heterozygosity across individuals and tissue types suggests that a biological explanation cannot be ruled out.

Currently, there is limited information available on the life cycle of *Ahnfeltia*. The unexpected distribution of variants may suggest a more complex life cycle than the typical triphasic pattern, potentially involving somatic diploid sectors, cryptic diploid phases, or nuclear heterogeneity, particularly regarding the diploid stage in *Ahnfeltia* and even red algae. Although the haploid–diploid structure of red algal life cycles has been described (*143*), none of the genome-wide population studies have been conducted to evaluate ploidy rigorously. Thus, further integrative cytological and population genomic investigations will be necessary to clarify the extent and evolutionary significance of diploid stages in red algae.

Given the consistent detection of heterozygous variants and the lack of clear haploid patterns, we treated individuals as diploid in downstream population genomic analyses. When ploidy cannot be clearly determined or mixed ploidy cannot be excluded, assuming diploidy provides a practical and cautious analytical approach and reduces the risk of underestimating genetic variation. Although overall heterozygosity levels were low in the re-sequencing data, AMOVA revealed that covariance within individuals (3.06%) was comparable to that among individuals within populations (4.15%, table S9). This indicates that heterozygous variants are not negligible and contribute meaningfully to the total genetic variance. Although we did not separately re-sequence all putative diploid tissues, these detected variants still represent meaningful population-level diversity and evolutionary signals of *Ahnfeltia*.

**Supplementary Text S2: Time estimation using MCMCtree**

To estimate *Ahnfeltia* diversification time, we conducted Bayesian MCMCtree time estimation using nuclear BUSCO genes shared in current available red algal genomes. Because *Ahnfeltia* exhibits relatively low substitution rates despite its deep phylogenetic position, its low heterogeneity from ancestors may induce uncertainty in divergence time estimation. This challenge is reflected in previous studies, which reported conflicting speciation timelines: Bringloe and Saunders (*144*) suggested a speciation timeframe of approximately 5.3 Ma, while Yang et al. (*38*) proposed a notably earlier timeframe of 112 Ma. These results highlight the sensitivity of time estimates to model assumptions.

To account for potential rate variation among lineages, we applied both independent (calculating evolutionary rates independently in each clade) and correlated clock models (following other species' evolutionary patterns). Under the correlated model, *Ahnfeltia* speciation began approximately 190 Ma and around 20 Ma under the independent model (Fig. 2E and fig. S43). The two models imply contrasting evolutionary scenarios. Under the correlated model, evolutionary rates change gradually across lineages, resulting in divergence times that fit well within the broader timeline of red algal diversification. In contrast, the independent model implies a recent and substantial increase in evolutionary rate along the *Ahnfeltia* crown lineage in order to reconcile its deep phylogenetic placement.

The older estimate (ca. 190 Ma) overlaps temporally with major Mesozoic geological transitions, including the fragmentation of Pangea, which has been associated with large-scale marine environmental restructuring. During this interval, diversification of several macroalgal lineages has been inferred in previous studies (*145, 146*). Direct causal links cannot be established. However, given that *Ahnfeltia*'s ecology and physiology are not markedly different from other red seaweeds, this temporal correspondence is compatible with an early diversification scenario for *Ahnfeltia*.

By contrast, the recent estimate (ca. 20 Ma) implies prolonged evolutionary stasis followed by comparatively late crown diversification. While such patterns are not impossible, they would require substantial rate shifts relative to other red algal lineages. Thus, it might be challenging to explain a 480-million-year gap before speciation and a recent trigger that affected *Ahnfeltia* speciation, distinct from other red seaweeds.

This argument is further supported by the evolution rate patterns inferred from both models and the lineage-through-time (LTT) plot (fig. S43 and S44). The most abundant red algal group, Rhodymeniophycidae, exhibits a gradual decline in evolutionary rates from ancestral nodes. *Ahnfeltia* is expected to follow this pattern (the correlated model) rather than the abrupt increase of evolution rate in the crown group (the independent model, fig. S44). Moreover, most red algal speciation likely occurred in past periods. Unlike land plants, there are no known cases of recent and rapid speciation in red seaweeds driven by mechanisms such as polyploidization or hybridization (*38, 147*).

Taken together, the correlated clock model appears more consistent with the broader evolutionary rate patterns observed across red algae. Under this framework, *Ahnfeltia* is inferred to have originated in the Mesozoic and subsequently maintained a relatively low substitution rate compared with other red algal genera (Figs. 2D and 2E). Nevertheless, given model sensitivity and rate heterogeneity, divergence time estimates should be interpreted cautiously.

### Supplementary Text S3: Key genomic variation between *Ahnfeltia* species

#### Divergence between *A. plicata* and the *A. borealis*/*A. fastigiata* clade

*Ahnfeltia* species first diverged into two main clades: *A. plicata* and the clade comprising *A. borealis* and *A. fastigiata*, with the latter two diverging more recently (*79, 81*). *Ahnfeltia* species exhibit generally simple and similar morphologies. In particular, most morphological traits of *A. borealis* closely resemble those of *A. plicata* rather than *A. fastigiata* (*79*). *A. plicata* and *A. borealis* are taller (up to 12 cm) than *A. fastigiata* (up to 5 cm, see Fig. 1B). Additionally, cortical growth rings are absent in *A. fastigiata*.

*A. borealis* and *A. fastigiata*, phylogenetically closer, shares reproductive features, which includes the positioning of carposporophytes near the thallus apex and similar patterns of gonimoblast development (*79, 81, 118*). Although we detected genetic differentiation among the three species (Fig. 5A), the connection between these genomic differences and the observed morphological traits remains unclear. Therefore, we further investigated how genetic and genomic variation may have contributed to the evolution of species-specific morphological characteristics.

To identify lineage-specific genomic changes between *A. plicata* and the *A. borealis*/*A. fastigiata* clade, we initially identified highly mutated regions, using short read sequencing data from each individual to the reference genomes to detect unmapped regions (UMRs), where long insertions/deletions happened (fig. S45A and S46). Highlighting UMRs can find notable disparity between the two species (*148*). We identified these UMRs between *A. plicata* and *A. borealis*/*A. fastigiata* clade based on GATK results after short read mapping. The functions of genes within these UMRs were associated with protein binding, catalytic activity, carbohydrate binding (fig. S45D).

Secondly, we found that the inversion of a segment in scaffold 17 and scaffold 26 commonly occurred in *A. borealis* and *A. fastigiata* genomes compared to *A. plicata* genome (fig. S45B). The gene ontologies (GOs) of genes within these inverted regions were associated with ribosome function, and proteolysis (fig. S45E). Furthermore, duplicated genes between *A. plicata* and *A. borealis*/*A. fastigiata* clade encompassed GOs terms of protein binding, phosphatase activator activity, and mismatch repair (fig. S45C and 45F, table S14). Environmental stresses may have played an important role to initiate divergence of *A. plicata* and the common ancestor of *A. borealis*/*A. fastigiata* as GO terms of oxidoreductase activity were identified (fig. S45E and S45F).

#### Role of TEs in *A. fastigiata* evolution

The burst insertion of TEs can lead to genome size expansion or chromosomal rearrangements, influencing both transcriptional and epigenetic regulation (*149-151*). These changes are significantly associated with the species incompatibility in *Drosophila*, fungal species, and plants (*152-154*). The abrupt changes of TEs are attributed to interspecific and intergeneric hybridization, as well as response to environmental stress (*155-157*). Numerous studies have highlighted the involvement of TEs in adaptive evolution, affecting fitness and resulting in phenotypic consequences (*158-160*).

The assembled genomes size of *A. fastigiata* were an approximately 2 Mb larger than those of *A. plicata* and *A. borealis*. This increase in genome size was primarily attributed to the expansion of transposable elements (TEs) in the *A. fastigiata* genomes (table S7). From this result, it was postulated that *A. fastigiata* diverged from the common ancestor of the *A. borealis*/*A. fastigiata* with an introduction of TE. To more specify the locations of TE insertions, we examined the number of TEs around comparable orthologue genes across the five *Ahnfeltia* genomes. Significant TE insertions were identified in the upstream regions of 537 orthologue genes, and in the downstream of 488 orthologue genes (fig. S47A and 47B). Some repeats were also found in intronic and genic regions of three and five genes, respectively. One of the LTRs, copia was identified in the intronic region of one gene (OG0007094; fig. S47C). Interestingly, the GO of this gene was associated with gamma-tubulin ring complex (fig. S47D).

*cis*-Regulatory elements, including promoters and intronic enhancers, play an essential role in gene expression, which affects plant development and environmental responses (*161, 162*). Some studies verified that TE insertions could prevent this gene regulatory elements, leading to phenotypical changes such as downsizing of plants (*163, 164*). Given that one notable morphological distinction of *A. fastigiata* was downsized thallus, the changes in tubulin proteins may be significant. Gamma tubulins are crucial proteins for microtubule nucleation, and gamma tubulin ring complex is involved in microtubule anchoring as a template (*165, 166*). Thus, genetic changes on this protein likely implicated in structural variation in microtubules, which affected the downsizing of *A. fastigiata*. Interestingly, another gene (OG0004475) involved in microtubule interaction and transport protein exhibited high dN/dS (1.34) indicating the positive selection in this gene (fig. S47E, S48, and table S15), which also supported that the genetic variation in tubulin may have affected morphological variation of *A. fastigiata*.

**Limitations and interpretation**

Establishing direct causal links between genomic changes and phenotypic differences in ancient lineages with relatively simple morphologies is challenging. In red algae, the genetic basis of key morphological traits is still poorly understood. Therefore, the genomic differences identified here may not represent the direct drivers of speciation in *Ahnfeltia*. Nevertheless, our analyses reveal distinct lineage-specific genomic modifications, providing candidate loci and pathways for future functional investigation. Experimental validation, including functional genomic approaches, will be necessary to clarify the mechanistic basis of species divergence.

**Supplementary Text S4: Additional analyses and inference to identify the causes of limited divergence in *Ahnfeltia***

We observed strong geographic isolation among *Ahnfeltia* lineages, with little evidence of recent gene flow following their migration (fig. S13 to S17). Although a signal of gene flow was detected from Brunswick to Ireland and Wales (fig. S49), this event was not recent based on our time estimation results (Fig. 4B). The overall pattern of limited diversification in *Ahnfeltia* implies that, after geographic isolation, these lineages did not undergo significant genomic or genetic changes. Rather than experiencing rapid secondary radiation, they appear to have persisted with relatively modest divergence. In this context, *Ahnfeltia* species may represent long-standing evolutionary lineages that have maintained genetic continuity despite prolonged geographic separation.

To explore potential genetic explanations for this pattern, we examined genome-wide variation among populations. However, we detected few lineage-specific differences, indicating limited accumulation of novel mutations and gene contents (Figs. 4 to 6). In other taxa, such as gar, reduced evolutionary rates have been linked to enhanced DNA repair mechanisms (*167*). We therefore tested whether a similar mechanism might underlie the slow evolutionary dynamics in *Ahnfeltia*, but there was no significant association (fig. S50 and table S16). These results suggest that alternative factors may better explain the limited evolution.

The predominance of homozygosity and population isolation suggests that *Ahnfeltia* populations may have had limited opportunities for genetic exchange with diverse individuals and genomic reshuffling over time. Unlike brown and green seaweeds, red algae lack flagellated gametes, which constrains long-distance gamete dispersal. To compensate, many red algae, including *Ahnfeltia*, exhibit a triphasic life cycle that enhances reproductive success (*168*). Nevertheless, successful fertilization still depends largely on local water movement, potentially limiting gene flow among distant populations.

In addition, red algae do not always complete the full sexual cycle. Under certain environmental conditions, they may rely more on vegetative growth or asexual reproduction (*143, 169*). Environmental factors such as substrate stability and habitat disturbance can influence this shift (*170, 171*). In *Ahnfeltia*, apomeiosis has been reported, in which gametophytes derived from tetraspores retain the same genetic composition as the parental thallus (*142, 172*). Such reproductive modes could further reduce effective recombination and slow the accumulation of genomic divergence.

We also observed this ecological persistence and slow vegetative growth in a simple long-term observation experiment. Over a four-year period, *Ahnfeltia* individuals remained viable and survived under the same conditions (4 °C) where co-occurring coralline algae eventually died (fig. S51). Although this is preliminary, this observation suggests that *Ahnfeltia* may possess considerable ecological resilience. Such resilience could buffer populations against environmental fluctuations, potentially reducing the strength of selective pressures that drive rapid diversification.

Taken together, these findings suggest that *Ahnfeltia* lineages may have experienced long-term evolutionary constraint. While it remains unclear whether ancestral diversity within *Ahnfeltia* was once greater than at present, contemporary populations show limited genetic differentiation and little evidence of ongoing speciation. This contrasts with genera such as *Gelidium* and *Gracilaria* (Rhodymeniophycidae), where diversification has been associated with ecological expansion and habitat shifts (*173, 174*). Similarly, compared with commercially exploited and anthropogenically influenced seaweeds such as *Undaria pinnatifida* (*175*), *Ahnfeltia* exhibits relatively high homozygosity and inbreeding coefficients (fig. S52). Overall, these patterns suggest that limited gene flow, life-history characteristics, and ecological resilience may together have contributed to the relatively low species richness observed in *Ahnfeltia*.

### Supplementary Text S5 : Ecological niche of *Ahnfeltia*

#### Ecological Niche Model Performance

The *Ahnfeltia* models performed well spatially (mean AUC = 0.87, SD = 0.08) and maintained robust environmental generality (AUC = 0.76, SD = 0.14), achieving high final accuracy (AUC = 0.92 and CBI = 0.93). In contrast, *Gracilaria* models showed lower discrimination ability (spatial AUC = 0.68, SD = 0.12; environmental AUC = 0.67, SD = 0.24), though their final models still fit the training data well (AUC = 0.73 and CBI = 0.96; Supplementary Table 2). When using the GBIF occurrence dataset for *Ahnfeltia*, 277 occurrence records were retained after filtering, and we found that model performance across spatial and environmental folds was similar to that of the genetically verified dataset (Supplementary Table 2).

#### Suitability Projections

Through time, palaeoclimate reconstructions revealed contrasting patterns of environmental suitability between *Ahnfeltia* and *Gracilaria* (Fig. 7A, fig. S25, and S26). Environmental suitability for *Ahnfeltia* was largely restricted to mid- to high-latitude coastal margins. In contrast, *Gracilaria* exhibited broader environmental suitability, with suitable areas extending into equatorial to subtropical coastal regions globally (fig. S27).

Reconstructed patterns of suitability by latitude further emphasized this long-term separation (Fig. 7B and 7C). *Ahnfeltia* maintained predominantly high-latitude environmental suitability (mean absolute latitude ~60°), whereas *Gracilaria* exhibited suitability concentrated at lower latitudes (~30–40°). Projections extended across the ocean and not limited to coastal grid cells reflected these same spatial patterns (fig. S28 and S29). In addition, projections from the GBIF-trained *Ahnfeltia* model closely resembled those generated using the genetically verified dataset (fig. S30).

Projections differed more between the Getech and Scotese-Temperature models and Getech and Scotese-$CO_2$ models than they did between Scotese-Temperature and Scotese-$CO_2$ across all model projections (n = 108 for *Ahnfeltia*, n = 60 for *Gracilaria*).

Analyses examining model extrapolation indicated temporal variation in climatic novelty across the Phanerozoic for both *Ahnfeltia* and *Gracilaria* ENMs (fig. S31). The proportion of suitable area with novel environmental conditions (MESS < 0) remained modest (< 0.4) through most of the Phanerozoic. Peaks in climatic novelty occurred primarily during the late Palaeozoic (around 320-280 Ma), when values exceeded 0.35 for all *Ahnfeltia* models. Patterns were broadly consistent among datasets, with the *Ahnfeltia* model showing slightly higher and more variable negative MESS than *Gracilaria*. Importantly, excluding all grid cells with negative MESS values prior to summarising latitudinal distributions had little effect, producing slightly lower estimates for both genera (fig. S32).

**Supplementary Figures 1-53**

**Fig. S1. Genome sizes of three *Ahnfeltia* species were estimated using *k*-mer analysis based on short-read sequencing data.** The nuclear genome sizes were approximately 30-40 Mbp.

**Fig. S2. *k*-mer profiles used to identify haploid and diploid individuals.** Samples with higher levels of heterozygosity exhibited more pronounced duplicated peaks.

**Fig. S3. Assembled genomes of three *Ahnfeltia* species.** Gene density (a), transposable element (TE) density (b), GC and AT content (c), Kimura value of retroelement of TEs (d), and Kimura value of DNA element of TEs were shown in Circos plot.

**Fig. S4. Telomeric regions recovered in five assembled *Ahnfeltia* genomes.** Many telomeric regions were recovered across all genomes. Particularly, the *A. plicata* UK genome recovered nearly all telomeric regions.

**Fig. S5. Functional annotation results following gene prediction in five *Ahnfeltia* genomes.** Functional annotations were conducted using BlastP against Uniprot and NCBI nr databases, as well as KASS, eggNOG, and InterProScan.

**Fig. S6. Evidence of constrained evolution of *Ahnfeltia* lineage in Rhodophyta based on plastid and mitochondrial phylogenies**. (A) Plastid phylogeny using 46 concatenated genes from plastid genomes of 282 red algal taxa and 10 *Ahnfeltia* populations. (B) Mitochondrial phylogeny using five concatenated genes from mitochondrial genomes of 209 red algal taxa and 10 *Ahnfeltia* populations.

**Fig. S7. Evidence of constrained evolution of *Ahnfeltia* lineage in Rhodophyta based on nuclear gene phylogenies using all orthologous genes.**

**Fig. S8. Low genetic differentiation of mitochondrial genes of *Ahnfeltia* compared to other red seaweeds.** (A) Comparison of the number of reported species and genetic distance of protein sequence using in mitochondrial phylogeny. (B) Branch length of *Ahnfeltia* and red seaweeds species from ancestral node in mitochondrial phylogeny.

**Fig. S9. Frequency and distribution of Kimura distances of transposable elements (TEs) across all *Ahnfeltia* species.** *A. fastigiata* genomes show evidence of relatively recent TE expansion.

**Fig. S10. Frequency and distribution of Kimura distances among transposable element (TE) families.** DNA/hobo-Activator, LINEs, and LTR elements (including LTR/Copia and LTR/Gypsy/DIRS1) show relatively recent expansion in the *A. fastigiata* genomes.

**Fig. S11. Comparison of singleton genes between *Ahnfeltia* (Three species five populations) and *Gracilaria* genomes (Four species).** The proportion of singletons indicates higher conservation of gene content in *Ahnfeltia* genomes compared with *Gracilaria* genomes.

**Fig. S12. Evolution of orthologue gene family (OGFs) of five *Ahnfeltia* species, nine red algal species, *Rhodelphis marinus*, and four species of land plants and green algae.** Gene gain and loss events were inferred using the Dollo parsimony principle, and clusters of orthologous groups (COGs) were functionally annotated based on eggNOG results.

**Fig. S13. Principal component analysis (PCA) of three *Ahnfeltia* species across ten populations based on SNPs mapped to five different reference genomes, using all detected variants to assess population structure.** PCA plots revealed clear genetic segregation among populations.

**Fig. S14. Principal component analysis (PCA) of three *Ahnfeltia* species across ten populations based on SNPs mapped to five different reference genomes, using all pruned data to assess population structure.**

**Fig. S15. Genetic relationships among *Ahnfeltia* populations inferred from a phylogenetic network using all detected variants (1,559,257 SNPs).**

**Fig. SPopulation admixture structure inferred for K = 2 to K = 14 based on all SNPs mapped to five different reference genomes.** Although Northern Hemisphere populations (Brunswick, Ireland, and Wales) exhibited minor admixture signals at K = 8–11, most *Ahnfeltia* populations were clearly segregated at each K value. Overall, the results indicate limited evidence of recent gene flow among populations.

**Fig. S17. Population admixture structure inferred for K = 2 to K = 14 based on pruned SNPs mapped to five different reference genomes.**

**Fig. S18. Populational divergence time estimation based on SNAPP analysis.** Pruned SNP datasets mapped to five *Ahnfeltia* reference genomes were analyzed to assess the consistency of the inferred results.

**Fig. S19. Genomic landscape of fixation index (FST), genetic distance (Dxy), and nonsynonymous-to-synonymous substitution rate ratio (dN/dS) ratios with pairwise comparison of *Ahnfeltia plicata* and *A. fastigiata* populations.** All values were calculated based on the *A. plicata* Chile, *A. borealis* Sakhlain, and *A. fastigiata* Oregon genomes.

**Fig. S20. Phylogeny of sulfatase genes founded in *Ahnfeltia* genomes.** This result suggest that *Ahnfeltia* likely acquired sulfatase genes through horizontal gene transfer (HGT).

**Fig. S21. Inbreeding coefficient ($F_{IS}$) and observed homozygosity (Ho) across the Ahnfeltia genome in each population.** These results indicate high levels of allelic homozygosity and inbreeding.

**Fig. S22. Distribution of runs of homozygosity (ROH) blocks across *Ahnfeltia* genomes, identified using PLINK v1.9 for each population.**

**Fig. S23. Changes in effective population size (Ne) inferred using Stairway Plot 2 based on SNP datasets mapped to five reference genomes to assess the consistency of the results.**

**Fig. S24. Linkage disequilibrium (LD) decay of *Ahnfeltia* populations based on each *Ahnfeltia* genome.**

**Fig. S25. Predicted climatic suitability for *Ahnfeltia* and *Gracilaria* in coastal grid cells under the HadCM3 palaeoclimate reconstructions (using the Scotese-Foster $CO_2$ model; Valdes et al. (*85*)) at representative ~100 million-year intervals through the Phanerozoic (0, 102.6, 201.3, 301.3, 400 and 505 Ma**). Projections for *Gracilaria* are restricted to ≤300 Ma in accordance with its estimated crown lineage age. Suitability values are shown on a cloglog scale (0-1), where higher values indicate greater suitable environmental conditions for the genus. Coastal cells were enlarged for clearer visualisation but still represent suitability values from the 100-km coastal buffer (one grid cell adjacent to land).

**Fig. S26. Predicted climatic suitability for *Ahnfeltia* and *Gracilaria* in coastal grid cells under the HadCM3 palaeoclimate reconstructions (using the Getech-Foster $CO_2$ model; Valdes et al. (*85*)) at representative ~100 million-year intervals (0, 102.6, 201.3 and 286.8 Ma) spanning the late Palaeozoic to the present.** Getech-Foster $CO_2$ reconstructions are available only from 286 Ma onwards and therefore omit the earliest time slices shown in other figures. Suitability values are shown on a cloglog scale (0-1), where higher values indicate greater suitable environmental conditions for the genus. Coastal cells were enlarged for clearer visualisation but still represent suitability values from the 100-km coastal buffer (one grid cell adjacent to land).

**Fig. S27. Binary climatic suitability projections for *Gracilaria* and *Ahnfeltia* under alternative palaeoclimate reconstructions across the Phanerozoic**. Projections for *Gracilaria* are restricted to ≤300 Ma in accordance with its estimated crown lineage age. Continuous suitability values were converted to presence-absence using the sensitivity-specificity threshold (see Methods). Each row represents selected projections at approximately 100 Ma intervals, and each column corresponds to one of the three palaeogeographic frameworks: Scotese-Temperature, Scotese-Foster $CO_2$, and Getech-Foster $CO_2$ (see Methods for details, Valdes et al. (*85*)). The Getech-Foster $CO_2$ reconstructions are available only for younger intervals and therefore omit the earliest time slices. Coloured cells indicate areas predicted as suitable for *Gracilaria* only (orange), *Ahnfeltia* only (navy), or both genera (green), whilst light grey cells indicate unsuitable conditions for both. Coastal cells were enlarged for clearer visualisation but still represent suitability values from the 100-km coastal buffer (one grid cell adjacent to land).

**Fig. S28. Predicted climatic suitability for *Ahnfeltia* worldwide under alternative HadCM3 palaeoclimate reconstructions (using the Scotese-Foster $CO_2$ model; Valdes et al. (*85*)) at representative ~100 million-year intervals through the Phanerozoic (0, 102.6, 201.3, 301.3, 400 and 505 Ma).** Each row represents suitability projections at approximately 100 Ma intervals, and each column corresponds to one of the three palaeogeographic frameworks: Scotese-Temperature, Scotese-Foster $CO_2$, and Getech-Foster $CO_2$ (see Methods for details, Valdes et al. (*85*)). The Getech-Foster $CO_2$ reconstructions are available only for younger intervals and therefore omit the earliest time slices. Suitability values are shown on the cloglog scale (0-1), where higher values indicate more suitable environmental conditions for the genus.

**Fig. S29. Predicted climatic suitability for *Gracilaria* worldwide under alternative HadCM3 palaeoclimate reconstructions (using the Scotese-Foster $CO_2$ model; Valdes et al. (*85*)) at representative ~100 million-year intervals through the Cenozoic to late Paleozoic (0, 102.6, 201.3, and 286.8 Ma).** Each row represents suitability projections at approximately 100 Ma intervals, and each column corresponds to one of the three palaeogeographic frameworks: Scotese-Temperature, Scotese-Foster $CO_2$, and Getech-Foster $CO_2$ (see Methods for details, Valdes et al. (*85*)). The Getech-Foster $CO_2$ reconstructions are available only for younger intervals and therefore omit the earliest time slices. Suitability values are shown on the cloglog scale (0-1), where higher values indicate more suitable environmental conditions for the genus.

**Fig. S30. Fig. S30. Predicted climatic suitability for *Ahnfeltia* worldwide based on the GBIF-trained model under alternative HadCM3 palaeoclimate reconstructions (using the Scotese-Foster $CO_2$ model; Valdes et al. (*85*)) at representative ~100 million-year intervals through the Phanerozoic (0, 102.6, 201.3, 301.3, 400 and 505 Ma).** The GBIF-trained model includes a larger but non-genetically verified set of occurrence records (N = 277). Each row represents suitability projections at approximately 100 Ma intervals, and each column corresponds to one of the three palaeogeographic frameworks: Scotese-Temperature, Scotese-Foster $CO_2$, and Getech-Foster $CO_2$ (see Methods for details, Valdes et al. (*85*)). The Getech-Foster $CO_2$ reconstructions are available only for younger intervals and therefore omit the earliest time slices. Suitability values are shown on the cloglog scale (0-1), where higher values indicate more suitable environmental conditions for the genus. Coastal cells were enlarged for clearer visualisation but still represent suitability values from the 100-km coastal buffer (one grid cell adjacent to land).

**Fig. S31. Proportion of predicted climatically suitable area exhibiting novel climatic conditions (MESS < 0) through geological time for *Ahnfeltia* and *Gracilaria*.** Lines show the mean across simulations. Shaded ribbons indicate ±1 standard deviation and background colours denote major geological eras. Higher values indicate a greater fraction of suitable habitat in extrapolated climate space and increased uncertainty in deep-time projections.

**Fig. S32. Mean absolute latitude of suitable climatic conditions through geological time for *Ahnfeltia* and *Gracilaria* after accounting for environmental extrapolation.** Lines show the mean latitude of predicted presence cells for each time slice after excluding all grid cells with negative MESS values (*i.e.* environments outside the range of modern training conditions), averaged across palaeoclimate simulations. Shaded ribbons indicate ±1 standard deviation and background colours denote major geological eras.

**Fig. S33. Long-read coverage of the assembled *Ahnfeltia* genomes.** This plot reflects genome completeness and assembly continuity.

**Fig. S34. Allele frequency of *Ahnfeltia* populations based on *A. plicata* Chile genome.**

**Fig. S35. Comparison of allele frequency between *Ahnfeltia* species based on *A. plicata* Chile genome.**

**Fig. S36. Comparison of allele frequency between *Ahnfeltia* populations based on *A. plicata* Chile genome.**

**Fig. S37. Continued.**

**Fig. S38. Results of the momix R package used to calculate FWS and to identify haploid and diploid signals in *Ahnfeltia* individuals.**

**Fig. S39. Read depth of variants depended on types of alleles and populations.**

**Fig. S40. Ratio of alternative reads per all reads of *A. plicata* populations.** 0.5 value indicated heterozygous variants had exact half of alternative reads.

**Fig. S41. Ratio of alternative reads per all reads of *A. borealis* and *A. fastigiata* populations.**

**Fig. S42. Re-examining data for clarifying types of variants from female gametophytes of *Ahnfeltia*.** (A) Photos of tested tip branches (expected as haploid) and carposporophytes (expected as diploid). (B) Number of heterozygous sites of SNPs from re-sequencing data of tip branches and carposporophytes, which indicates a higher abundance of heterozygous signals (diploid) in carposporophytes. (C) Ratio of depths of alternative/reference alleles. In purely haploid samples, allele ratios are expected to approach 0 or 1, whereas diploid samples are expected to center around 0.5. The presence of intermediate ratios indicates that diploid cells are present within gametophyte tip branches.

**Fig. S43. Evolutionary rates and divergence time of *Ahnfeltia* in independent and correlated models used in MCMC time estimation.** Correlated model indicates evolutionary rates of *Ahnfeltia* gradually decreased from their ancestor. However, independent model indicates evolutionary rates of *Ahnfeltia* surged within recent short period.

**Fig. S44. Lineage-through-time (LTT) plot to compare independent and correlated models used in MCMC time estimation**. Correlated model indicates species diversity of red algae increased in past. Independ models indicates species diversity of red algae increased recently.

**Fig. S45. Genomic difference between *A. plicata* and *A. borealis*/*A. fastigiata* clade.** (A) Highly mutated regions between *A. plicata* and *A. borealis*/*A. fastigiata* clade verified with the unmapped regions of short read data. (B) Duplicated genes between *A. plicata* and *A. borealis*/*A. fastigiata* clade. (C) Inverted regions that occurred in *A. borealis*/*A. fastigiata* clade. (D) Gene Ontology (GO) of genes in the unmapped regions. (E) GO of duplicated genes. (F) GO of gene in inverted regions.

**Fig. S46. High divergence regions in chromosomes when *A. plicata* and *A. borealis*/*A. fastigiata* clades diverged.**

**Fig. S47. Key genomic and genetic differences associated with the divergence of *A. fastigiata*.** (A) Distribution of TE insertion sites in upstream and downstream intergenic regions of orthologue genes. (B) Number of TE insertions surrounding comparable orthologous genes across *Ahnfeltia* genomes. (C) Insertion of an LTR Copia element within a gene associated with the γ-tubulin ring complex. (D) Gene Ontology (GO) of orthologue genes where TE insertion occurred. (E) dN/dS analysis indicating positive selection in genes involved in microtubule interaction and transport.

**Fig. S48. dN/dS of comparable orthologue genes (6,897) of *Ahnfeltia* species.**

**Fig. S49. D-statistics and populational admixture test using Dsuite.** Limited gene flow was detected among populations; however, relatively strong gene flow from the Brunswick to the Ireland and Wales populations was observed, as indicated by high *f* values.

**Fig. S50. DNA repair systems of *Ahnfeltia* and other seaweeds were examined using the full set of DNA replication and repair pathways provided by KEGG (approximately 360 genes).** These genes were conserved in *Ahnfeltia* genomes, with no evidence of significant gene duplication or insertion.

**Fig. S51. Simple experimental results demonstrating the persistence of *Ahnfeltia*.** *Ahnfeltia* and coralline algae were stored at 4 °C for four years with two seawater changes. During this period, the coralline algae died, whereas *Ahnfeltia* survived.

**Fig. S52. Comparison of nucleotide diversity (π) and inbreeding levels between *Ahnfeltia* species and *Undaria pinnatifida*.** (A) Differences in nucleotide diversity (π) between *Ahnfeltia* and *U. pinnatifida.* (B) Proportion of the genome in runs of homozygosity, used to estimate the inbreeding coefficient based on ROH (FROH), comparing *Ahnfeltia* and *U. pinnatifida*. (C) Inbreeding coefficient (F) of each *Ahnfeltia* population compared with that of *U. pinnatifida.*

**Fig. S53. Re-examination of historical *Ahnfeltia* voucher specimens from the Berkeley Herbarium.** Most samples, including the type specimens of *A. gigartinoides* and *A. svensonii*, were identified as species belonging to the order Gigartinales. Only the samples from Uruguay and Chile were identified as *A. plicata*.

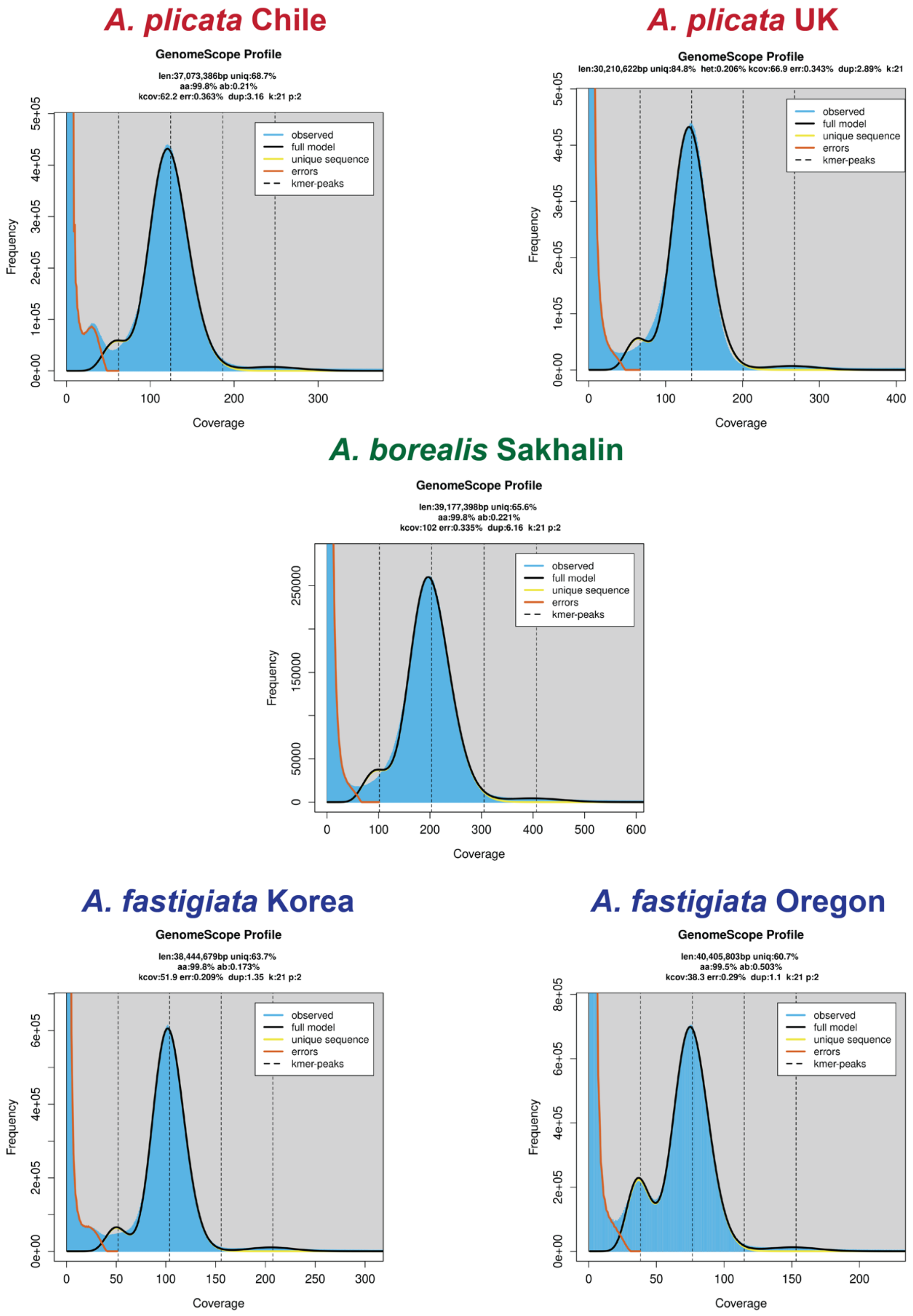

**Fig. S1. Genome sizes of three *Ahnfeltia* species were estimated using *k*-mer analysis based on short-read sequencing data.** The nuclear genome sizes were approximately 30-40 Mbp.

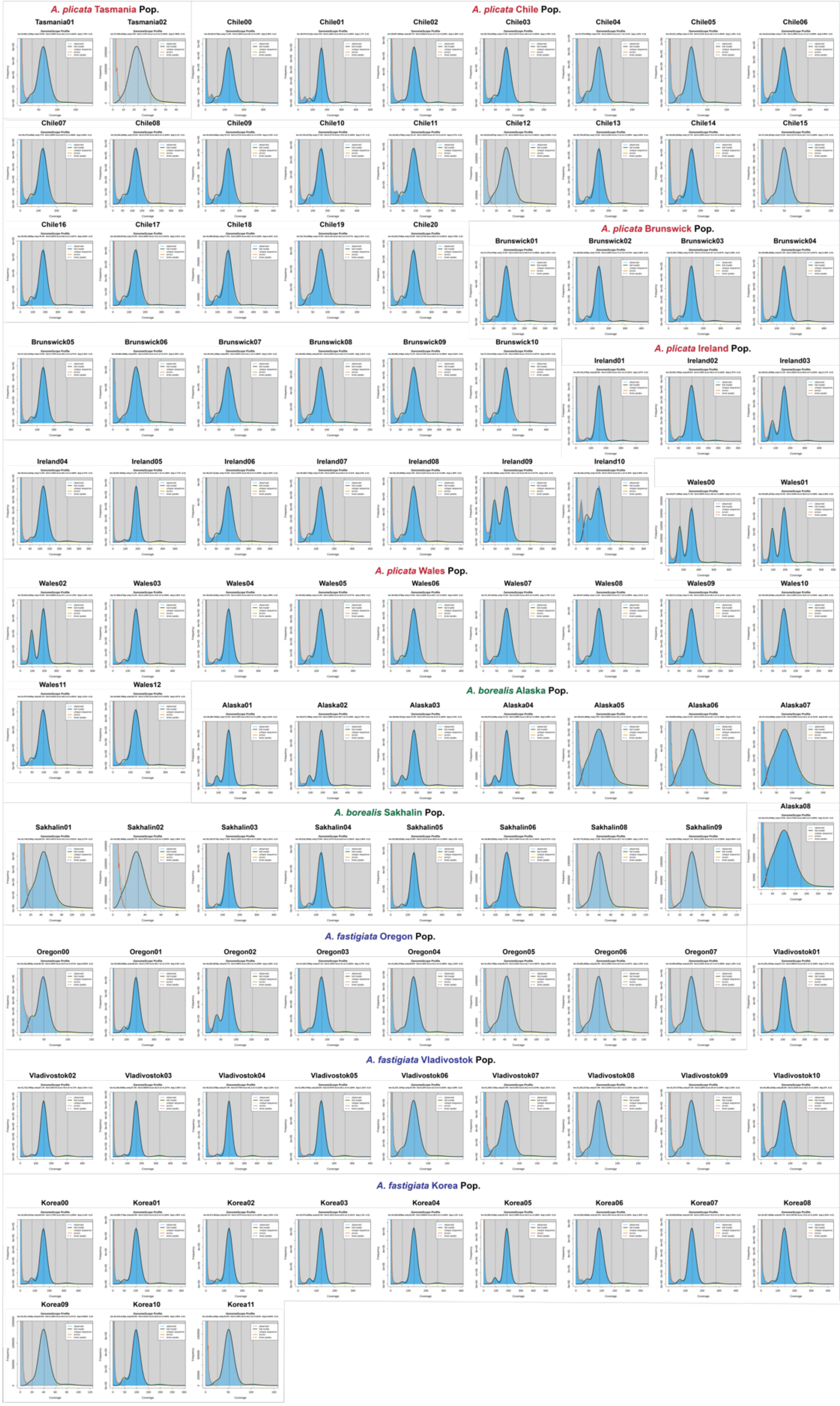

**Fig. S2. *k*-mer profiles used to identify haploid and diploid individuals.** Samples with higher levels of heterozygosity exhibited more pronounced duplicated peaks.

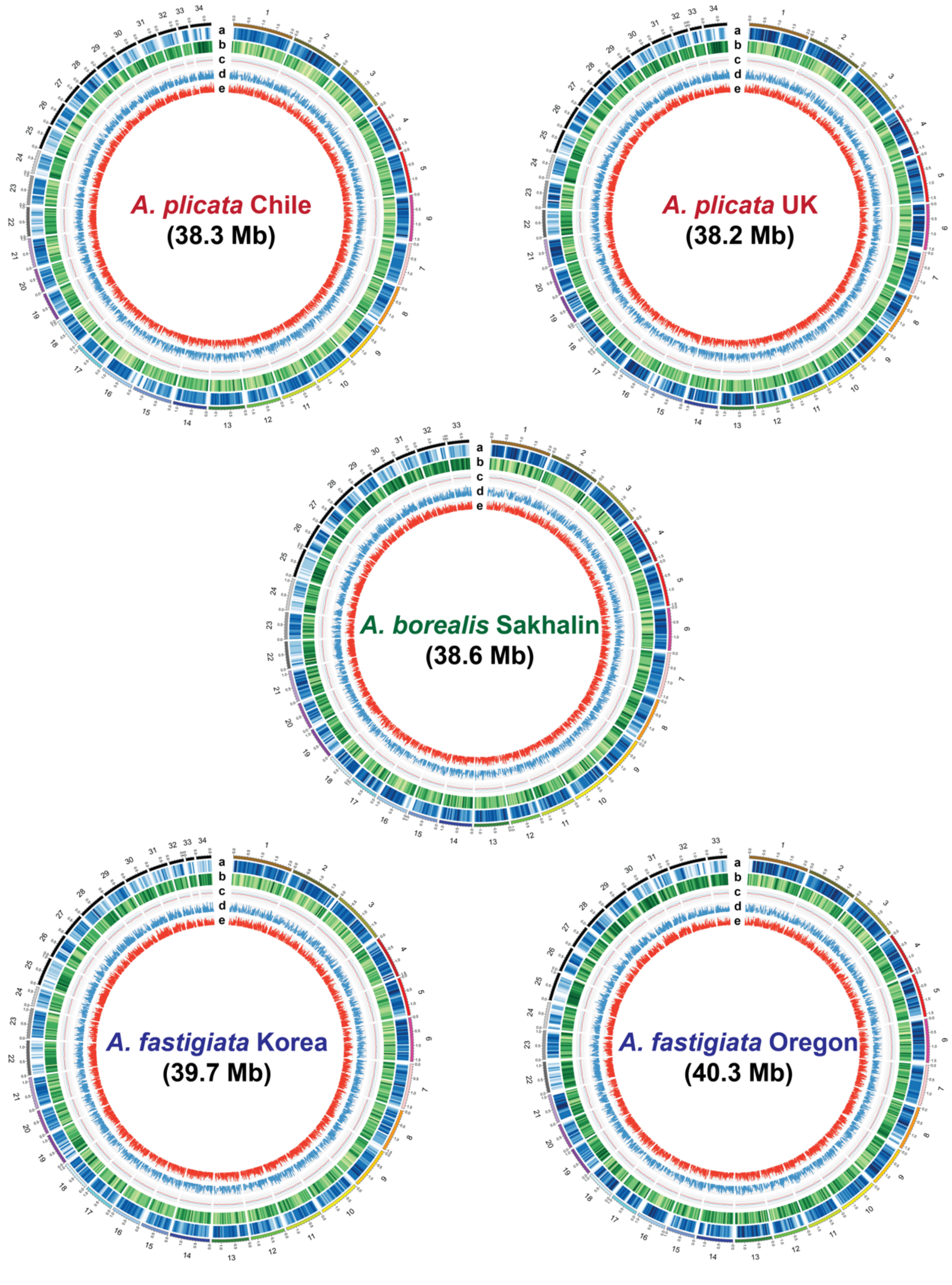

**Fig. S3. Assembled genomes of three *Ahnfeltia* species.** Gene density (a), transposable element (TE) density (b), GC and AT content (c), Kimura value of retroelement of TEs (d), and Kimura value of DNA element of TEs were shown in Circos plot.

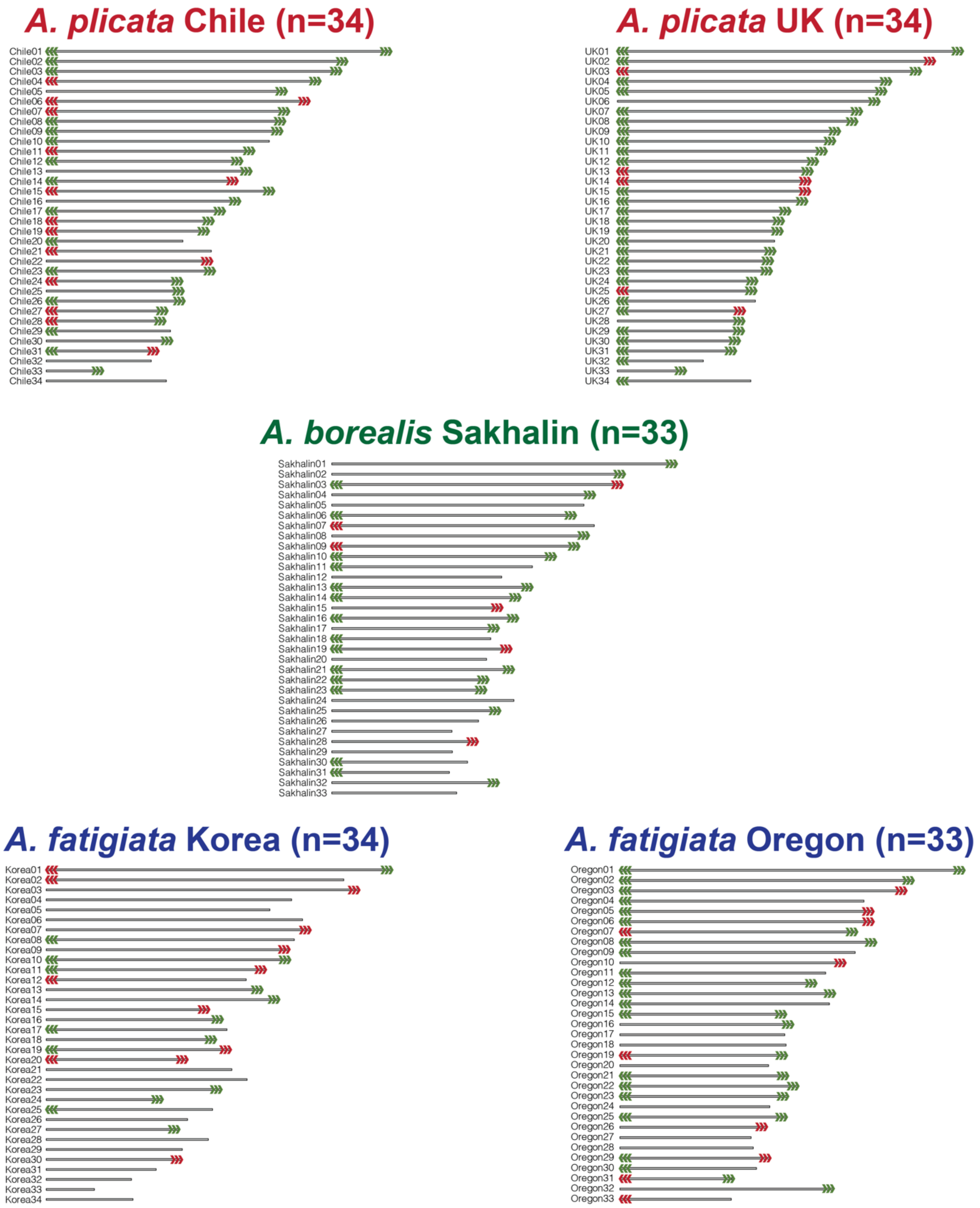

**❯❯❯ : Telomeric regions ❯❯❯ : Putative telomeric regions (including subteromeric regions)**

**Fig. S4. Telomeric regions recovered in five assembled *Ahnfeltia* genomes.** Many telomeric regions were recovered across all genomes. Particularly, the *A. plicata* UK genome recovered nearly all telomeric regions.

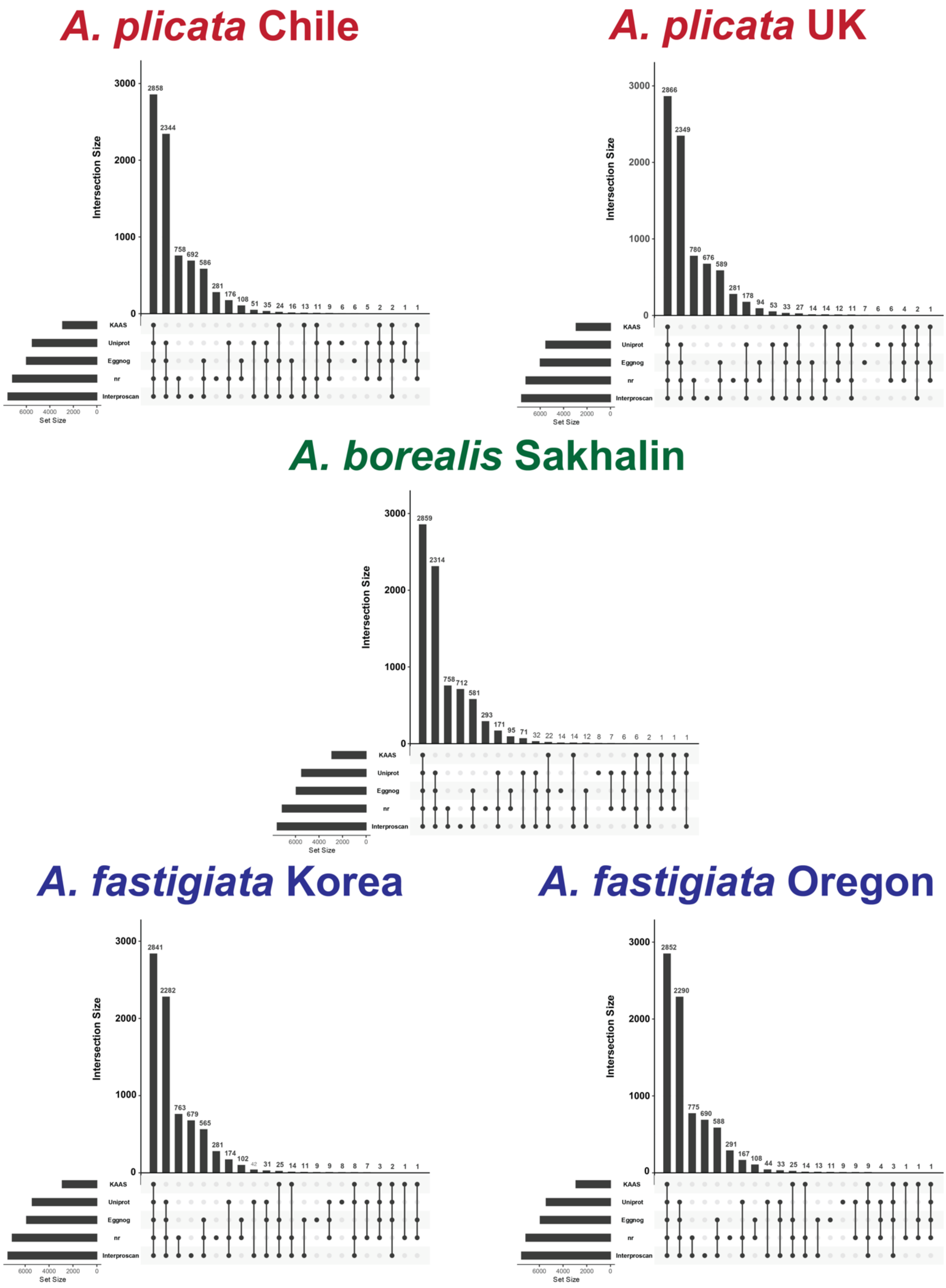

**Fig. S5. Functional annotation results following gene prediction in five *Ahnfeltia* genomes.** Functional annotations were conducted using BlastP against Uniprot and NCBI nr databases, as well as KASS, eggNOG, and InterProScan.

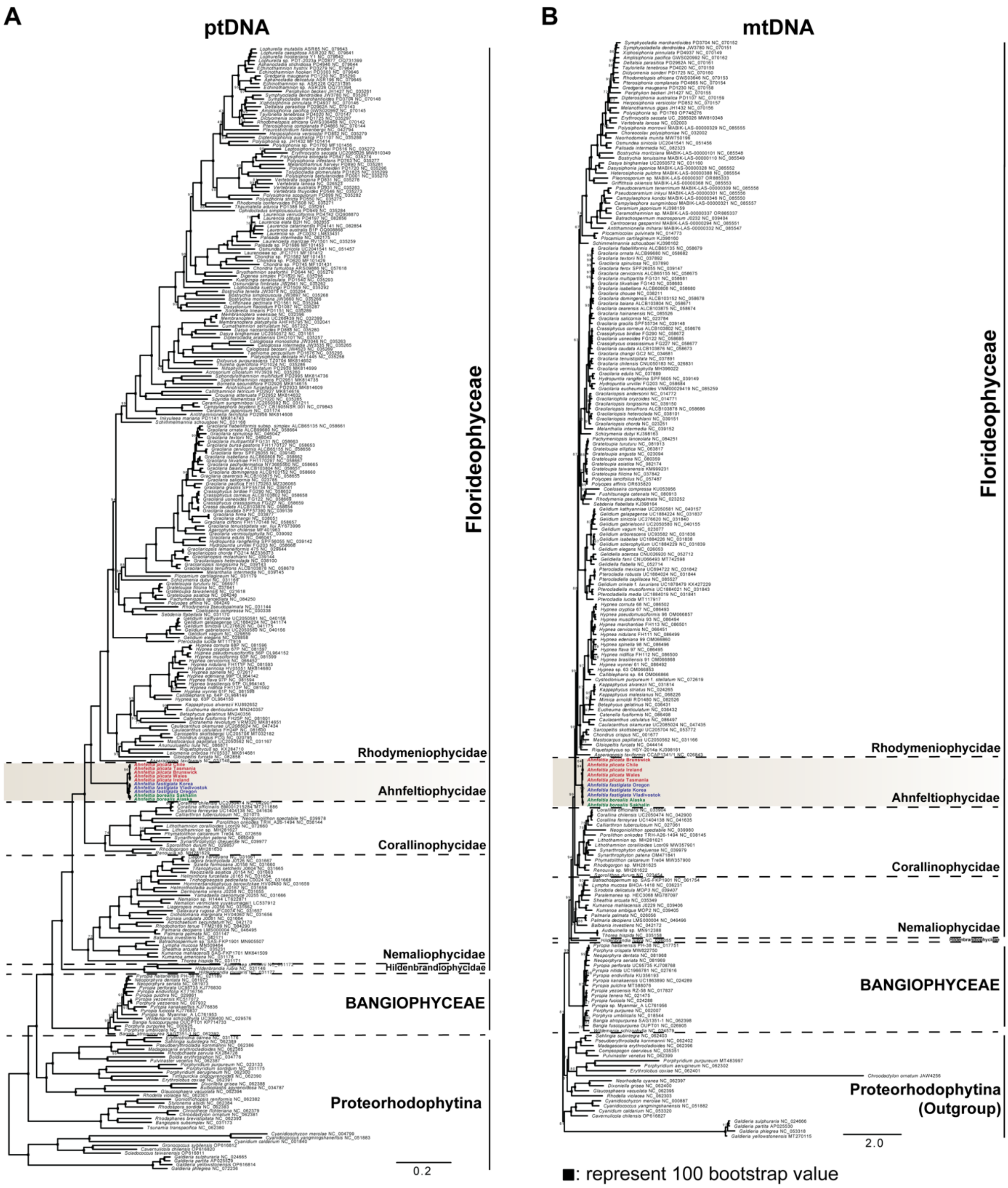

**Fig. S6. Evidence of constrained evolution of *Ahnfeltia* lineage in Rhodophyta based on plastid and mitochondrial phylogenies**. (A) Plastid phylogeny using 46 concatenated genes from plastid genomes of 282 red algal taxa and 10 *Ahnfeltia* populations. (B) Mitochondrial phylogeny using five concatenated genes from mitochondrial genomes of 209 red algal taxa and 10 *Ahnfeltia* populations.

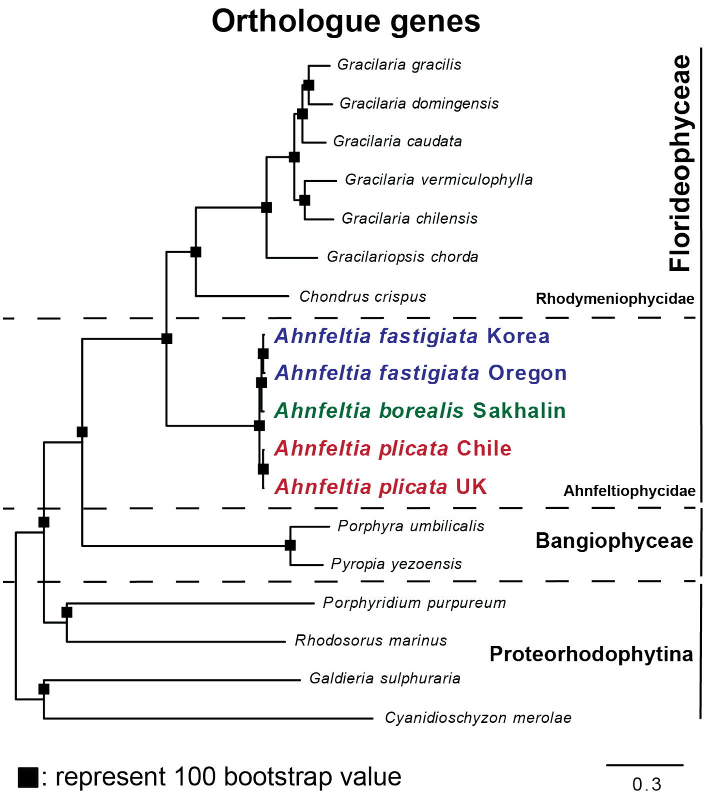

**Fig. S7. Evidence of constrained evolution of *Ahnfeltia* lineage in Rhodophyta based on nuclear gene phylogenies using all orthologous genes.**

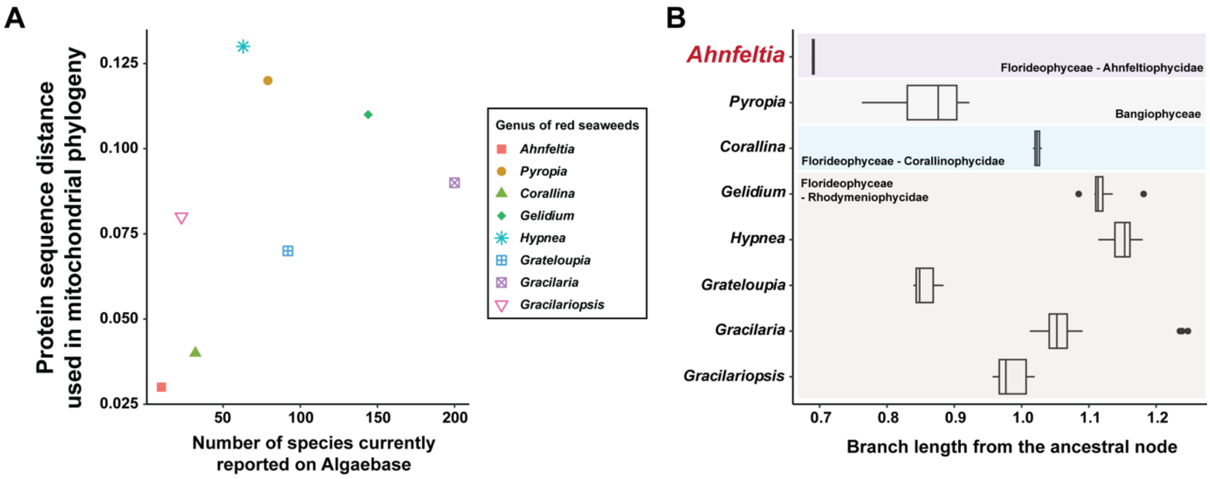

**Fig. S8. Low genetic differentiation of mitochondrial genes of *Ahnfeltia* compared to other red seaweeds.** (A) Comparison of the number of reported species and genetic distance of protein sequence using in mitochondrial phylogeny. (B) Branch length of *Ahnfeltia* and red seaweeds species from ancestral node in mitochondrial phylogeny.

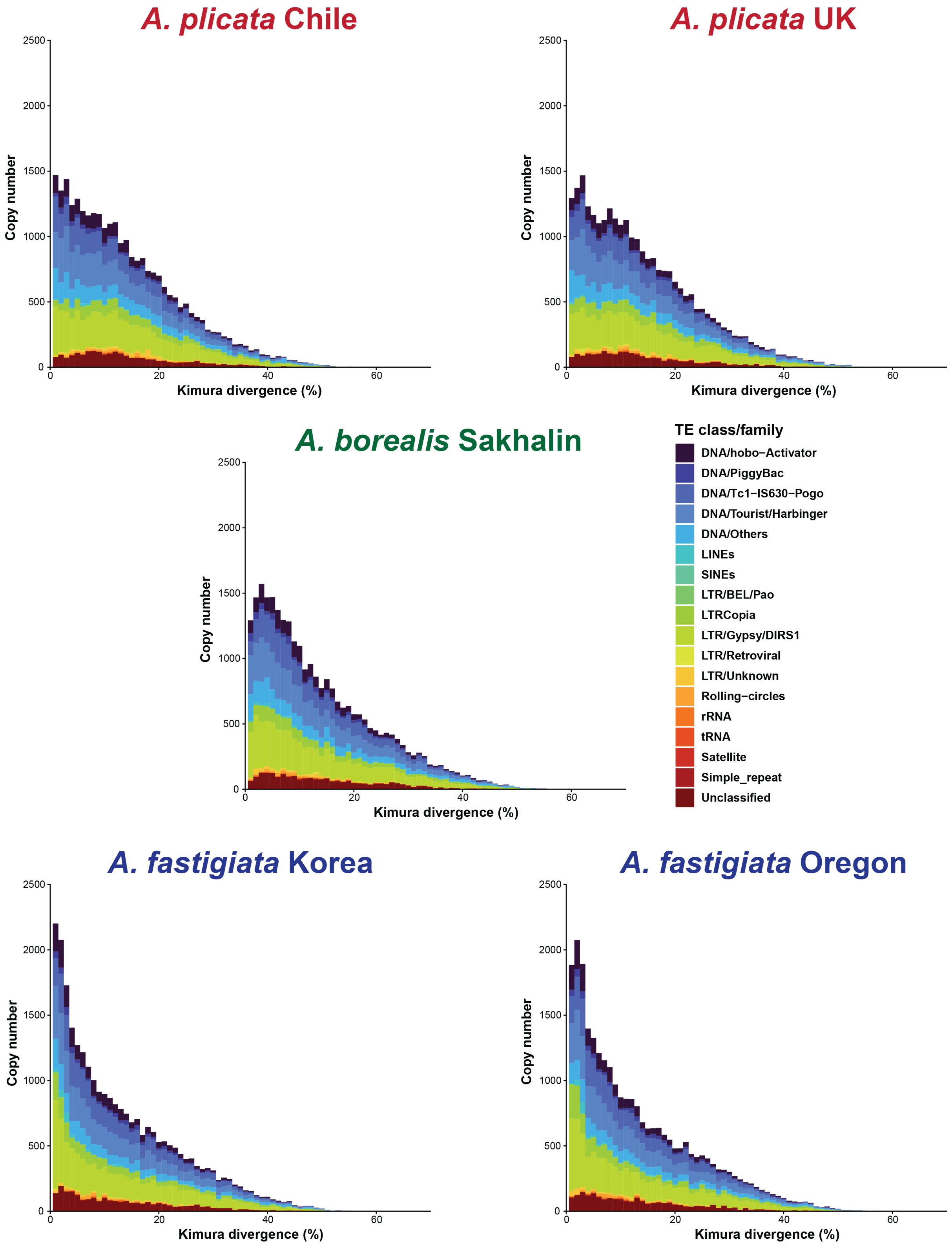

**Fig. S9. Frequency and distribution of Kimura distances of transposable elements (TEs) across all *Ahnfeltia* species.** *A. fastigiata* genomes show evidence of relatively recent TE expansion.

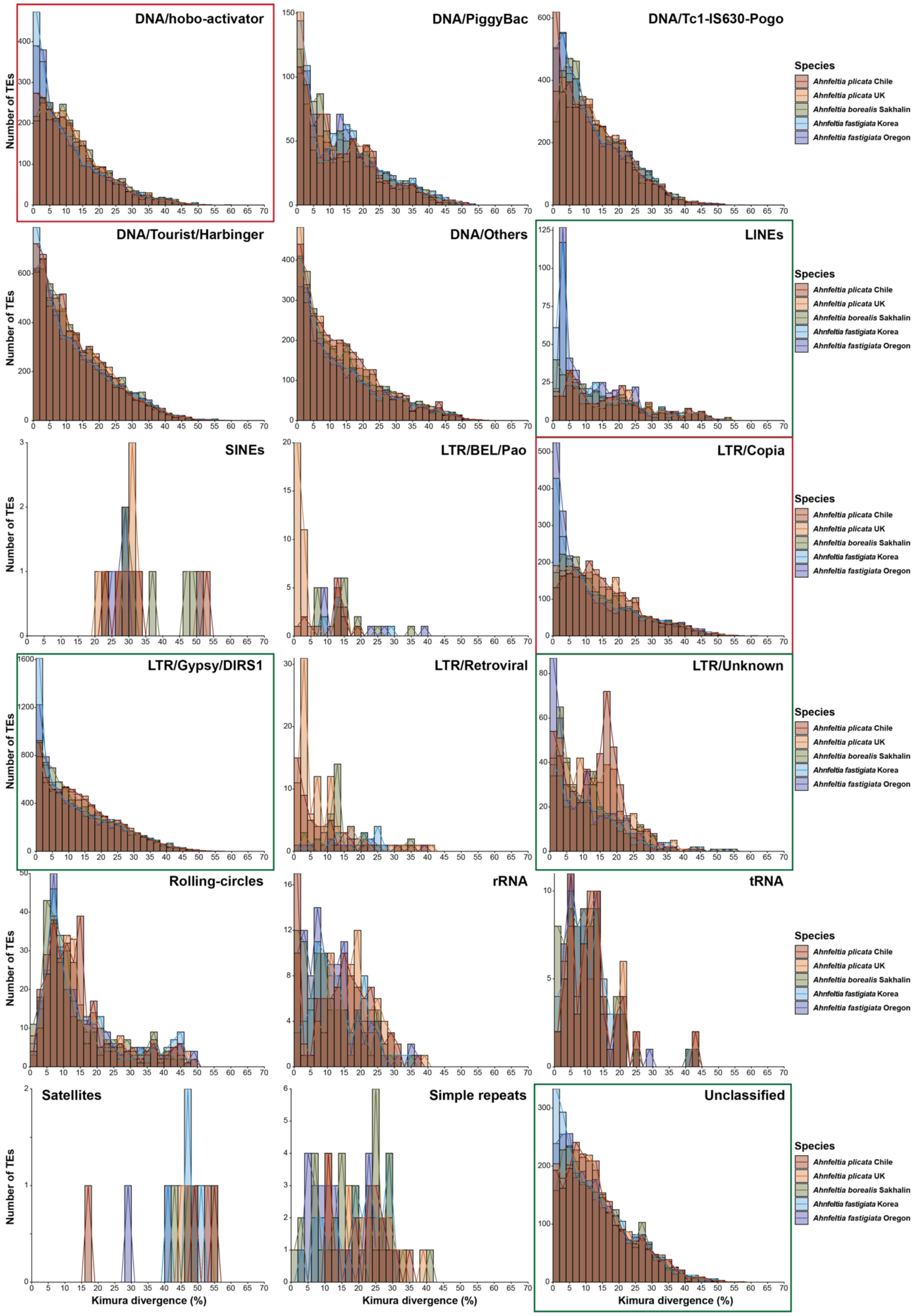

**Fig. S10. Frequency and distribution of Kimura distances among transposable element (TE) families.** DNA/hobo-Activator, LINEs, and LTR elements (including LTR/Copia and LTR/Gypsy/DIRS1) show relatively recent expansion in the *A. fastigiata* genomes.

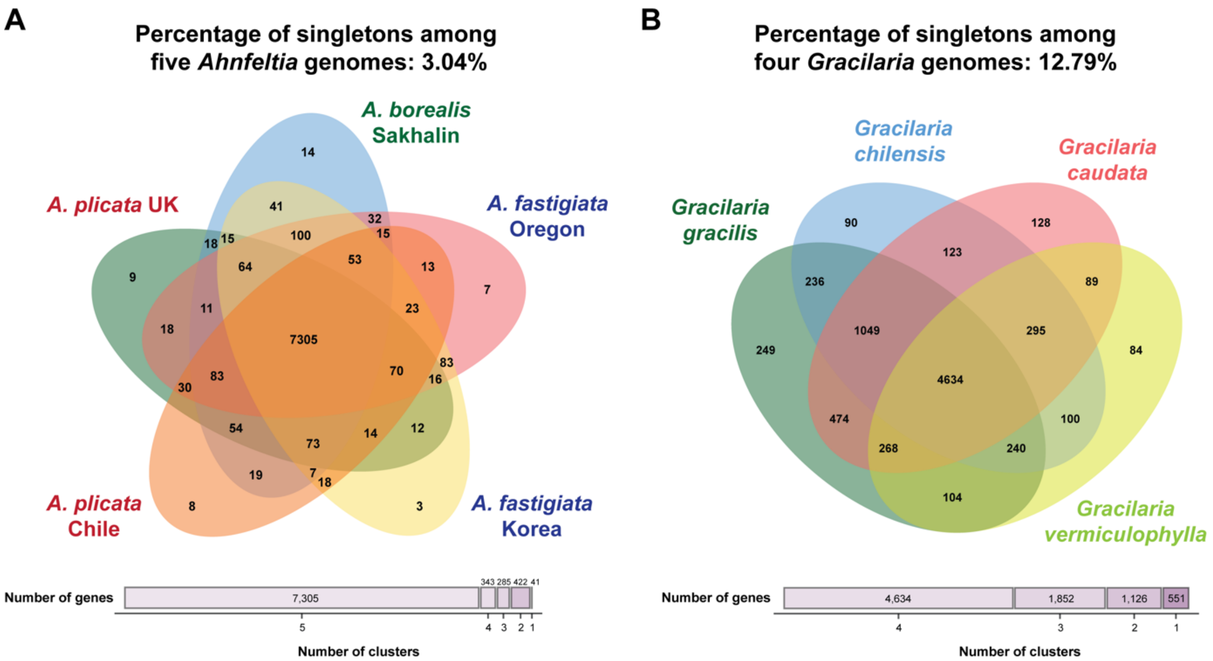

**Fig. S11. Comparison of singleton genes between *Ahnfeltia* (Three species five populations) and *Gracilaria* genomes (Four species).** The proportion of singletons indicates higher conservation of gene content in *Ahnfeltia* genomes compared with *Gracilaria* genomes.

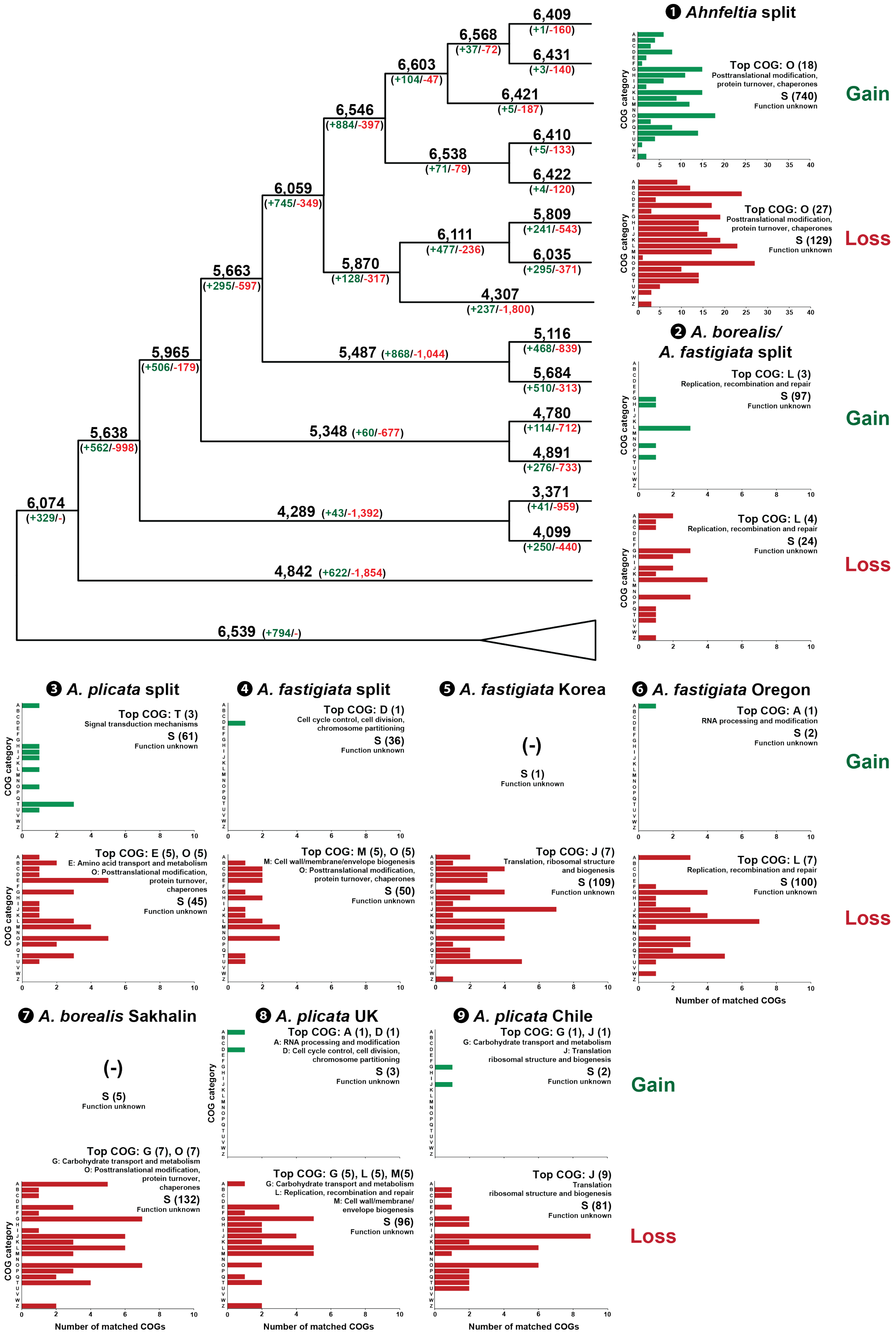

**Fig. S12. Evolution of orthologue gene family (OGFs) of five *Ahnfeltia* species, nine red algal species, *Rhodelphis marinus*, and four species of land plants and green algae.** Gene gain and loss events were inferred using the Dollo parsimony principle, and clusters of orthologous groups (COGs) were functionally annotated based on eggNOG results.

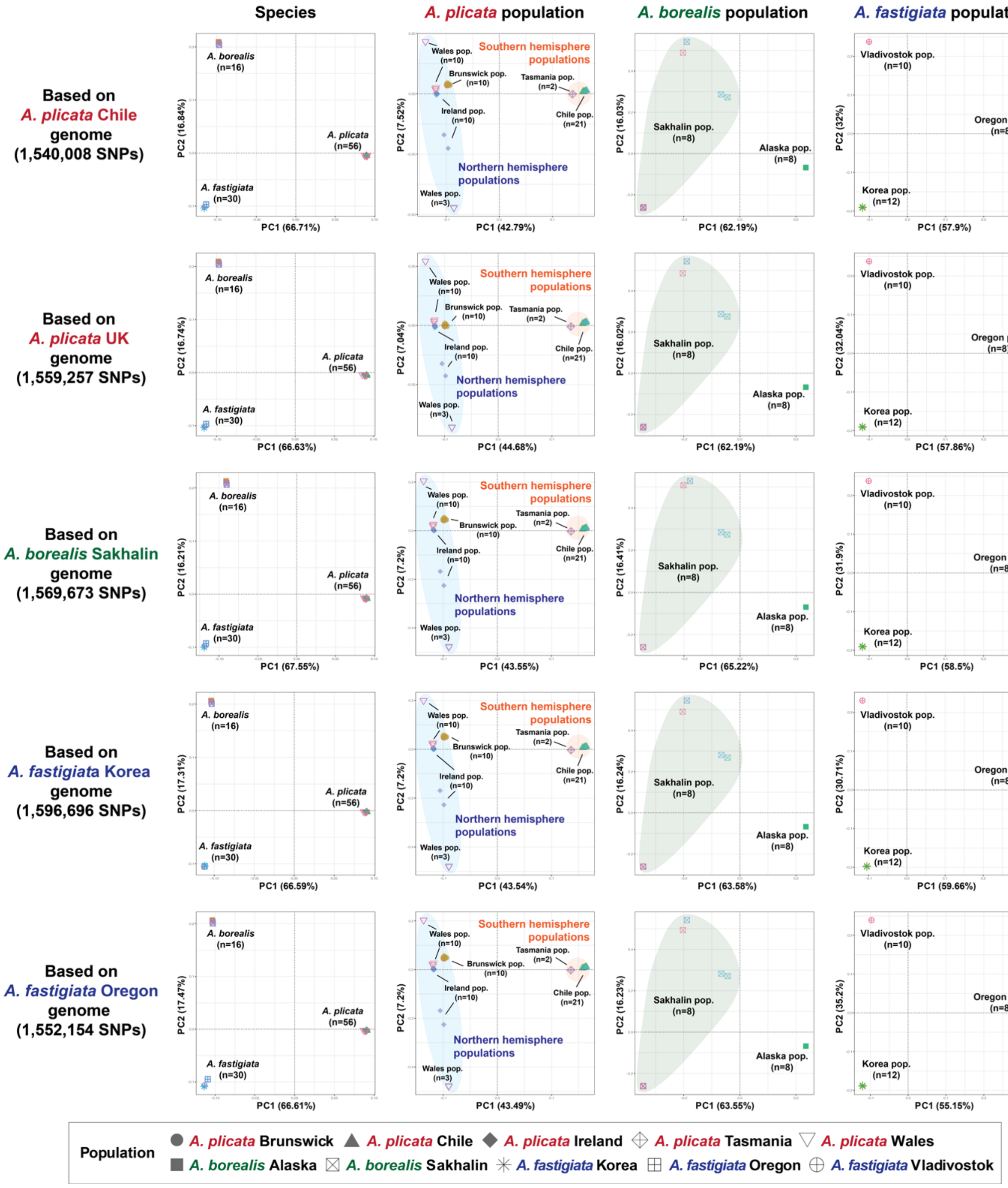

**Fig. S13. Principal component analysis (PCA) of three *Ahnfeltia* species across ten populations based on SNPs mapped to five different reference genomes, using all detected variants to assess population structure.** PCA plots revealed clear genetic segregation among populations.

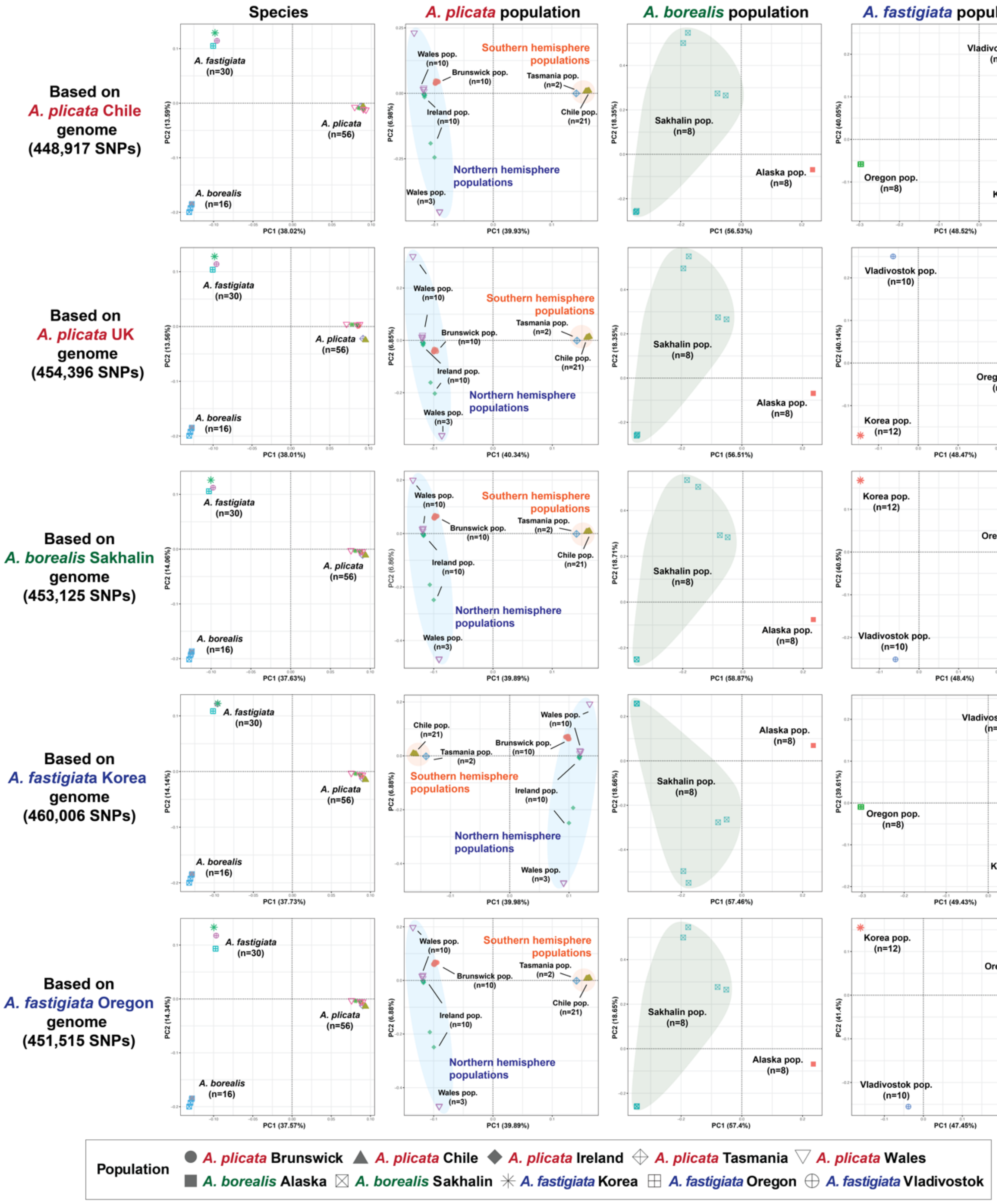

**Fig. S14. Principal component analysis (PCA) of three *Ahnfeltia* species across ten populations based on SNPs mapped to five different reference genomes, using all pruned data to assess population structure.**

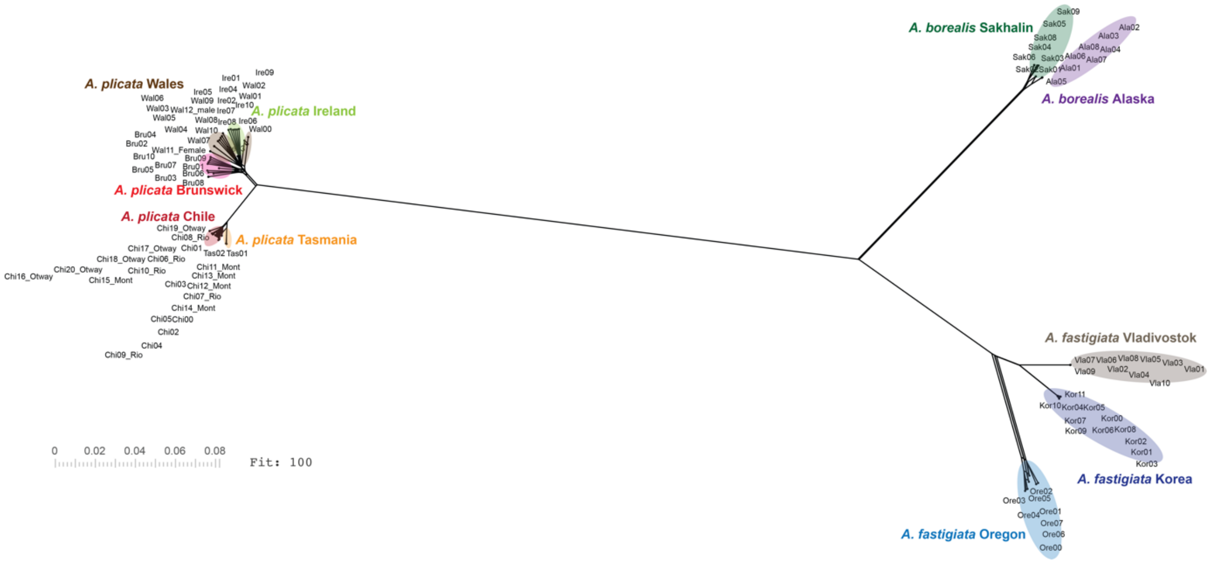

**Fig. S15. Genetic relationships among *Ahnfeltia* populations inferred from a phylogenetic network using all detected variants (1,559,257 SNPs).**

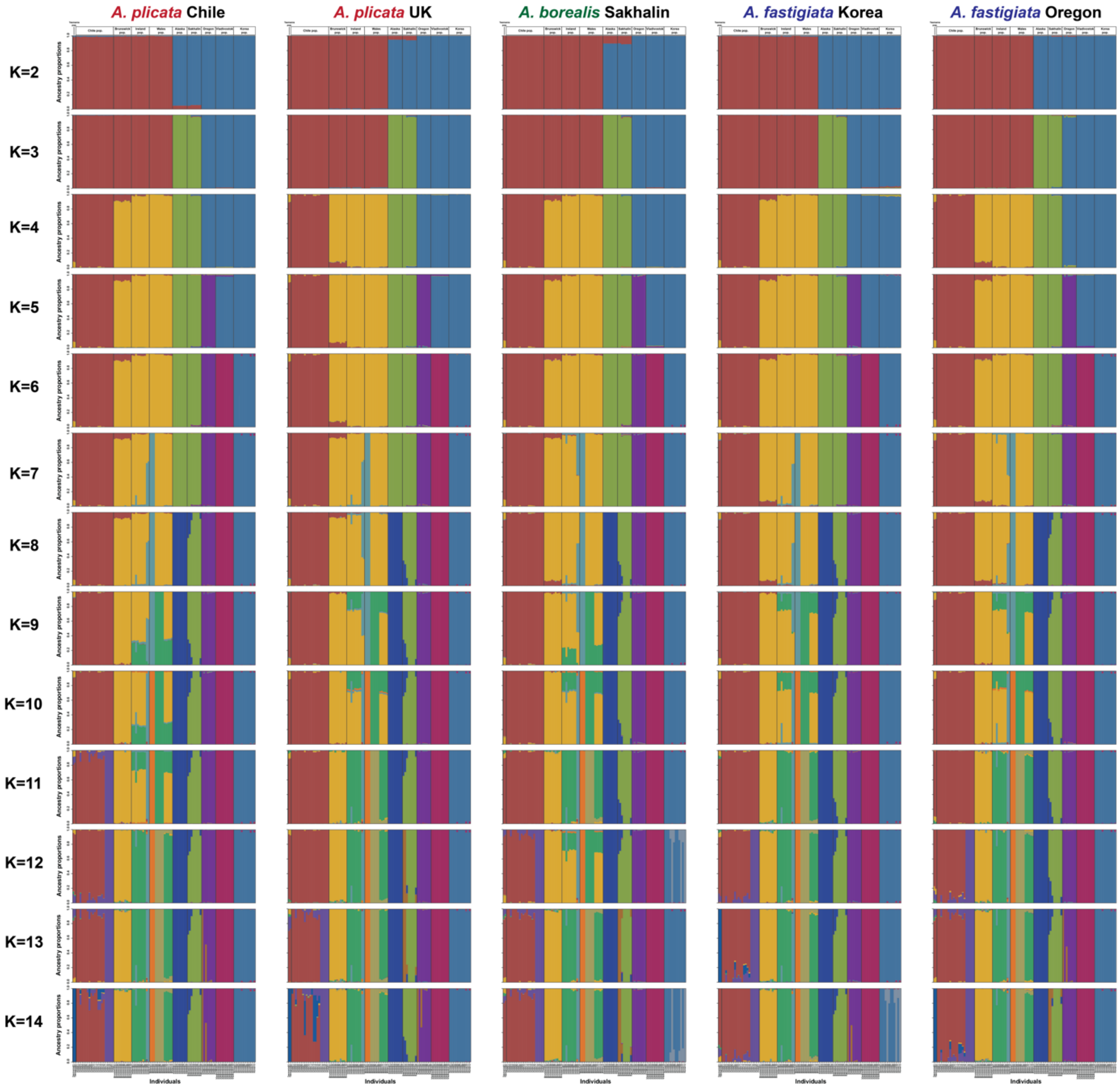

**Fig. S16. Population admixture structure inferred for K = 2 to K = 14 based on all SNPs mapped to five different reference genomes.** Although Northern Hemisphere populations (Brunswick, Ireland, and Wales) exhibited minor admixture signals at K = 8–11, most *Ahnfeltia* populations were clearly segregated at each K value. Overall, the results indicate limited evidence of recent gene flow among populations.

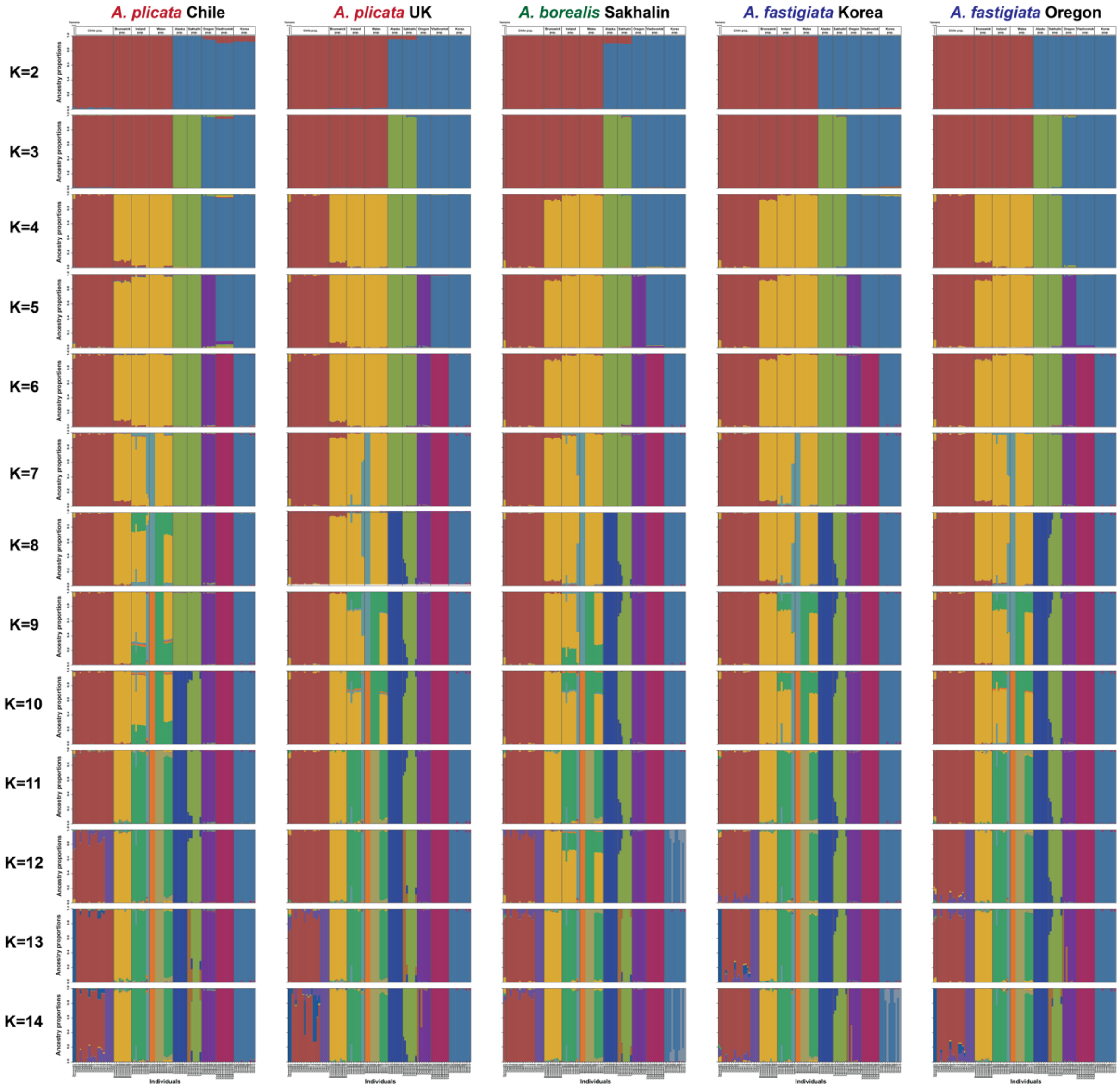

**Fig. S17. Population admixture structure inferred for K = 2 to K = 14 based on pruned SNPs mapped to five different reference genomes.**

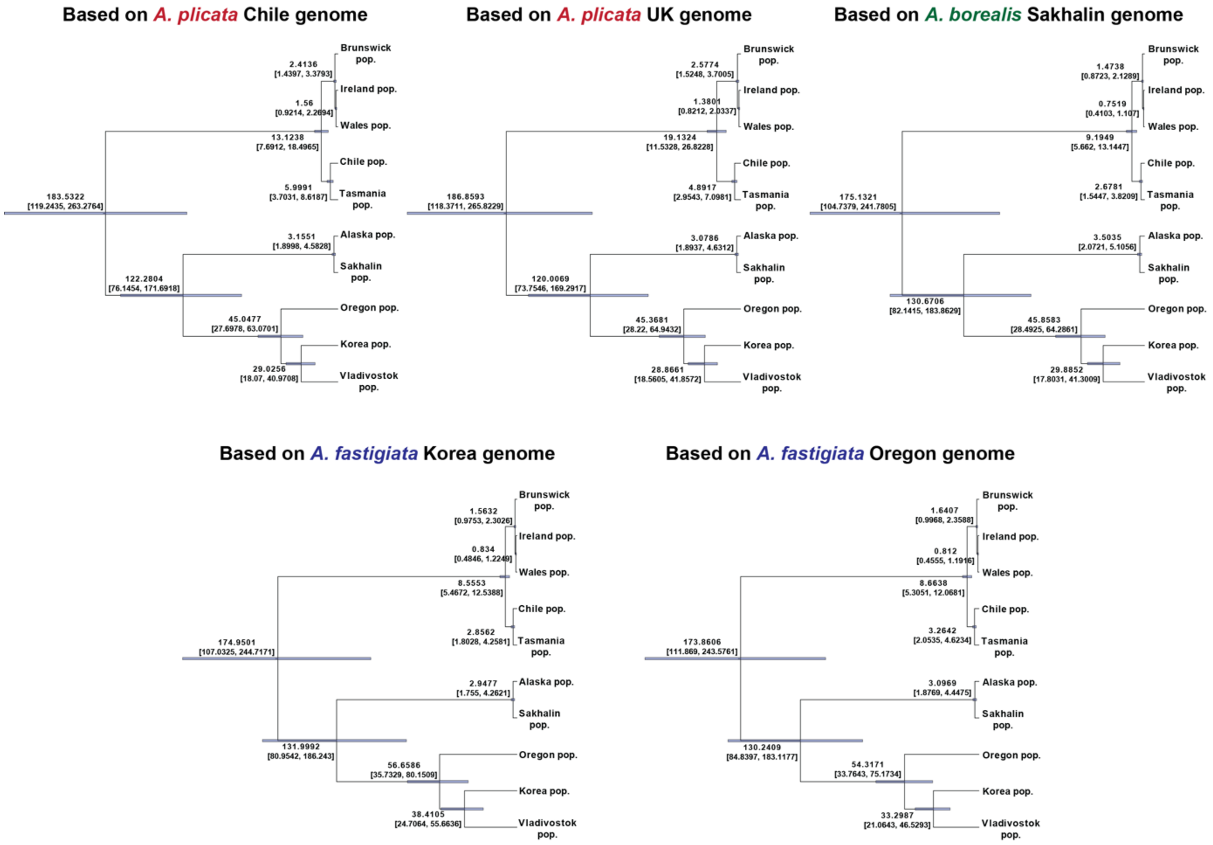

**Fig. S18. Populational divergence time estimation based on SNAPP analysis.** Pruned SNP datasets mapped to five *Ahnfeltia* reference genomes were analyzed to assess the consistency of the inferred results.

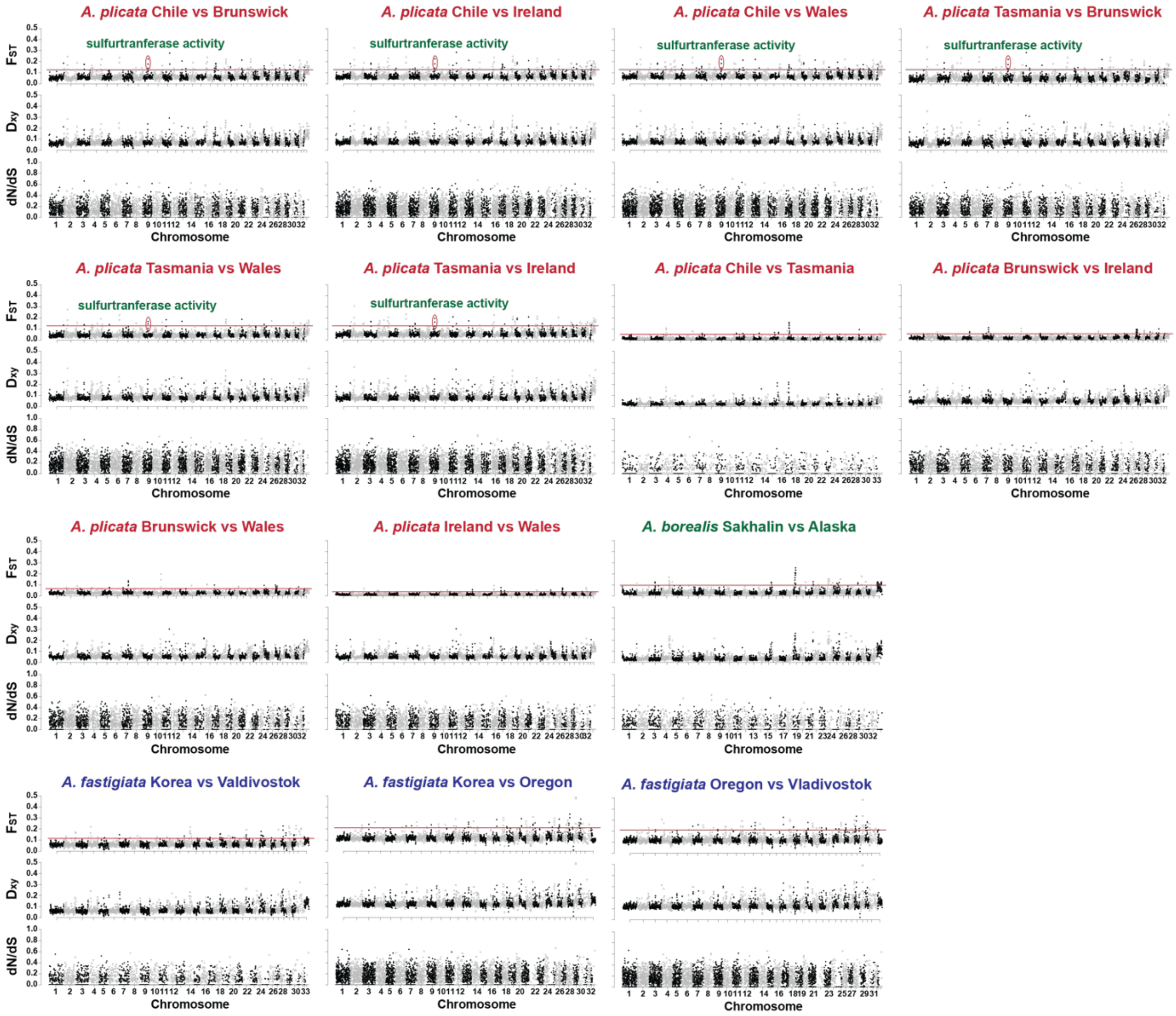

**Fig. S19. Genomic landscape of fixation index (FST), genetic distance (Dxy), and nonsynonymous-to-synonymous substitution rate ratio (dN/dS) ratios with pairwise comparison of *Ahnfeltia plicata* and *A. fastigiata* populations.** All values were calculated based on the *A. plicata* Chile, *A. borealis* Sakhlain, and *A. fastigiata* Oregon genomes.

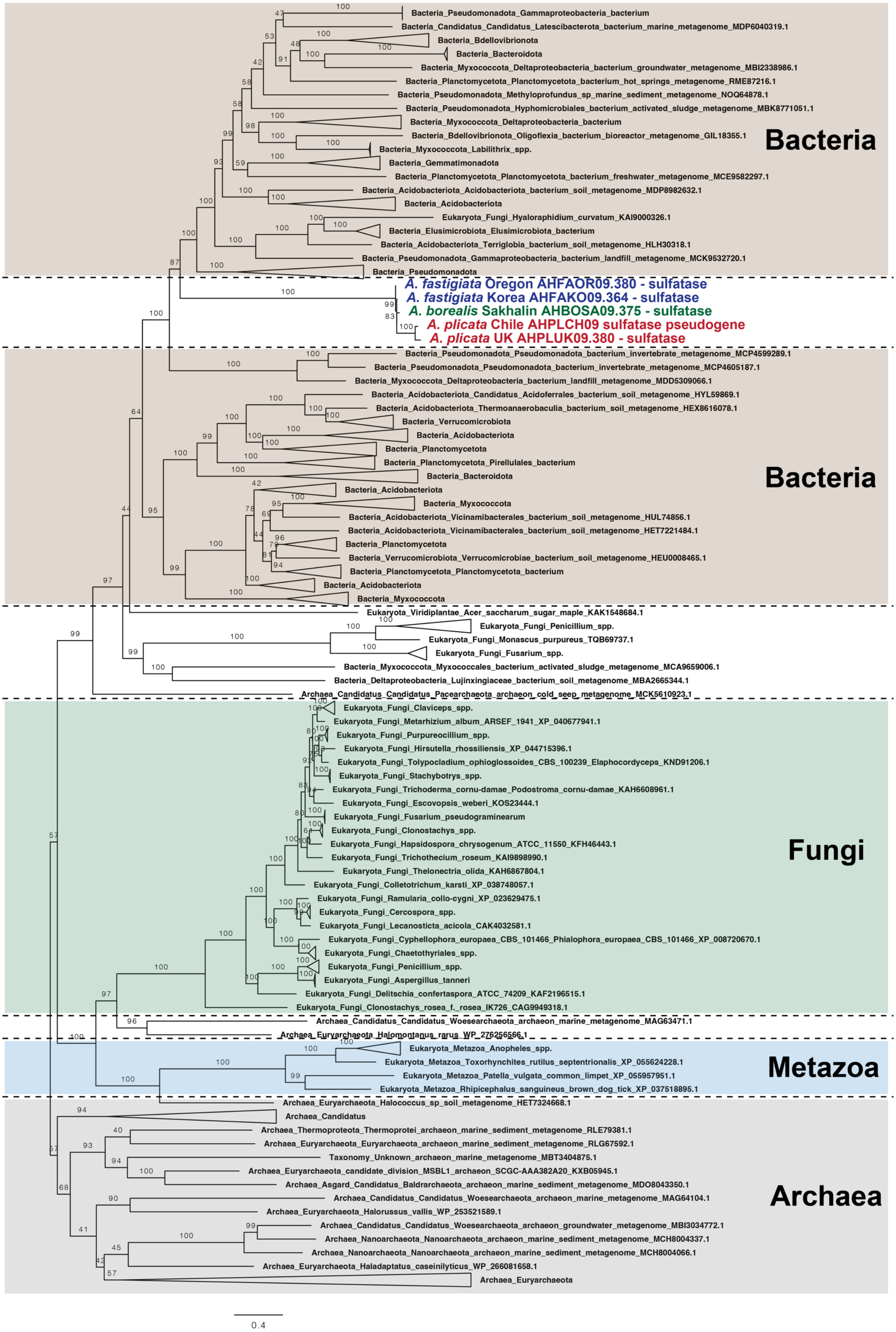

**Fig. S20. Phylogeny of sulfatase genes founded in *Ahnfeltia* genomes.** This result suggest that *Ahnfeltia* likely acquired sulfatase genes through horizontal gene transfer (HGT).

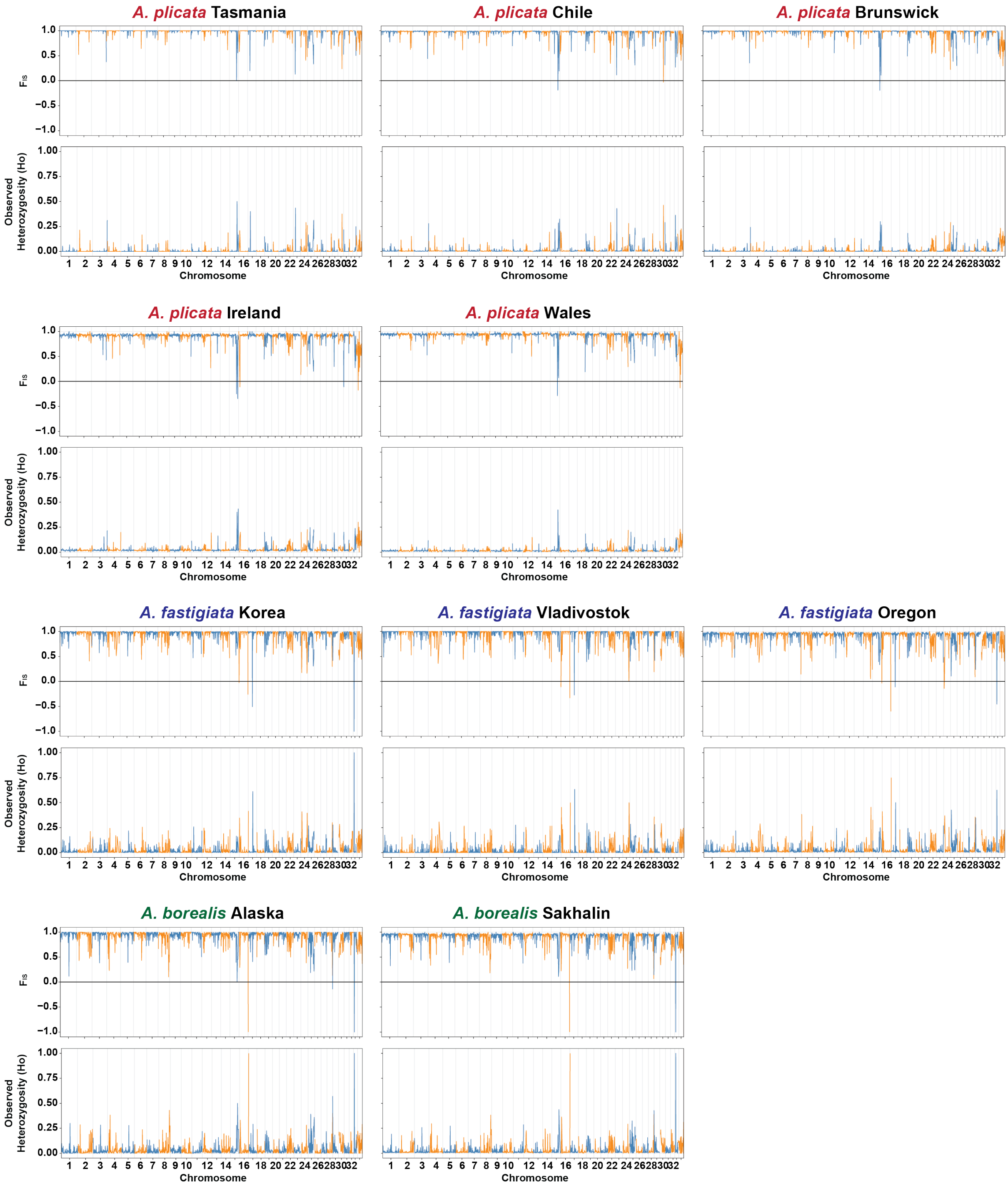

**Fig. S21. Inbreeding coefficient ($F_{IS}$) and observed homozygosity (Ho) across the Ahnfeltia genome in each population.** These results indicate high levels of allelic homozygosity and inbreeding.

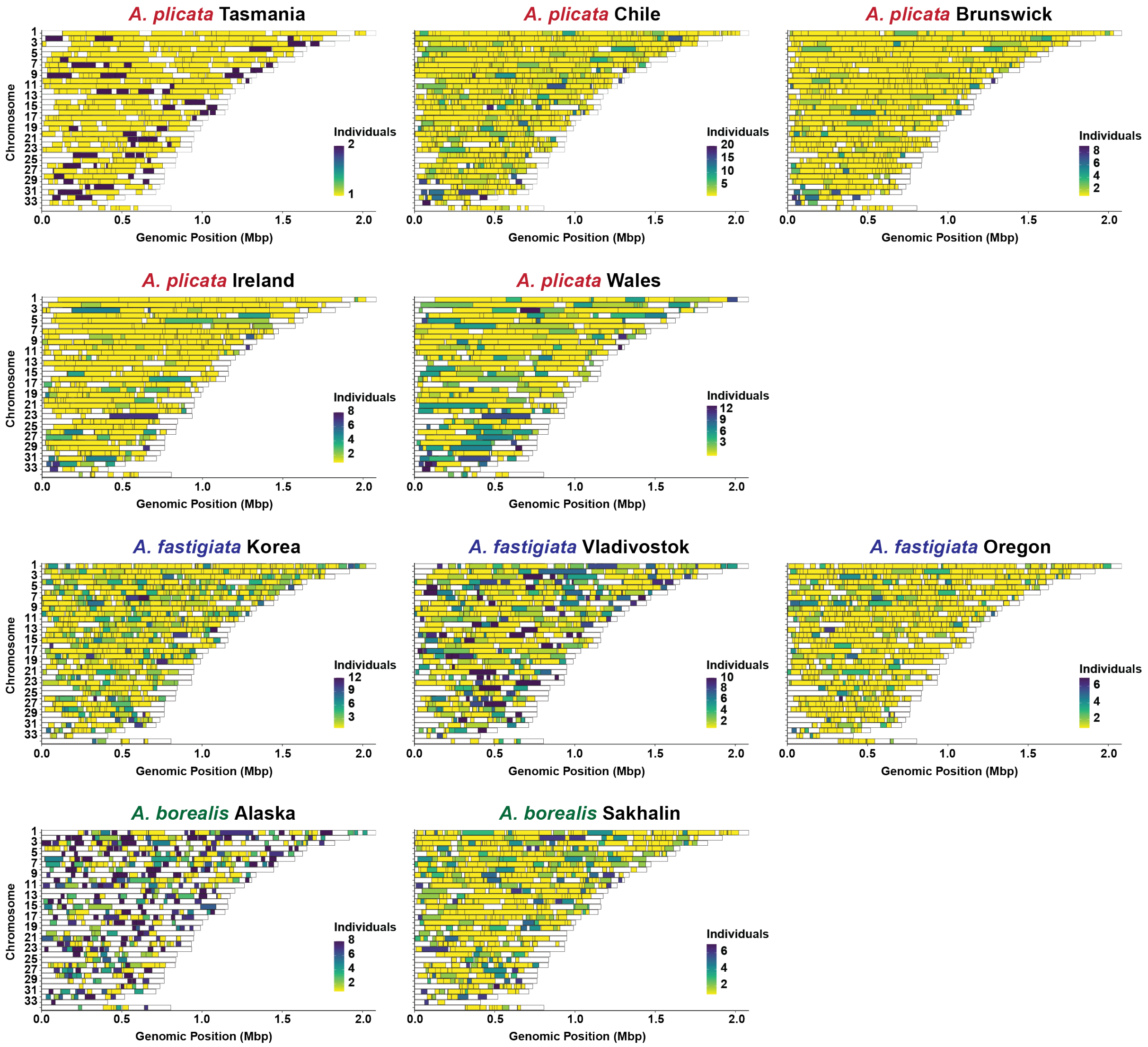

**Fig. S22. Distribution of runs of homozygosity (ROH) blocks across *Ahnfeltia* genomes, identified using PLINK v1.9 for each population.**

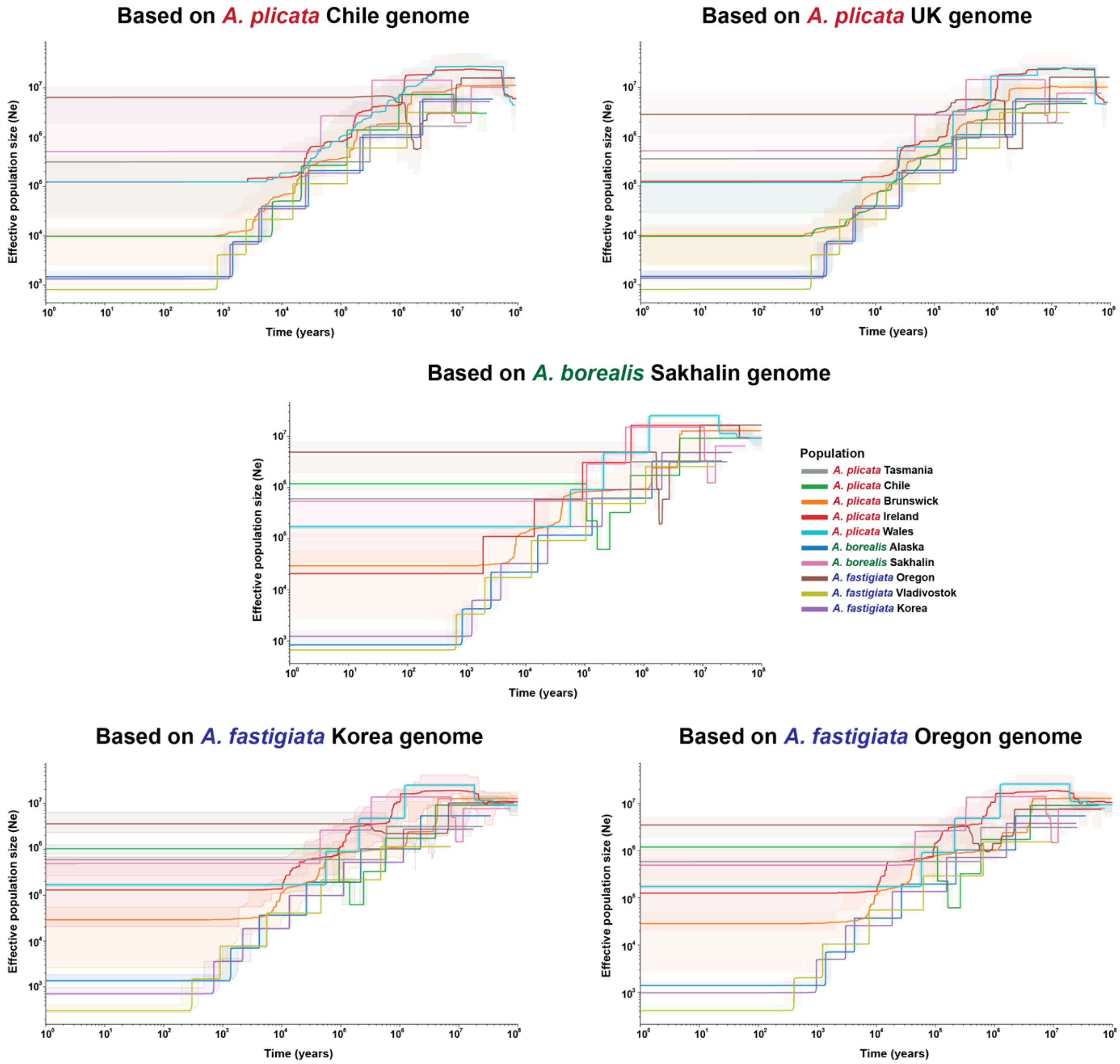

**Fig. S23. Changes in effective population size (Ne) inferred using Stairway Plot 2 based on SNP datasets mapped to five reference genomes to assess the consistency of the results.**

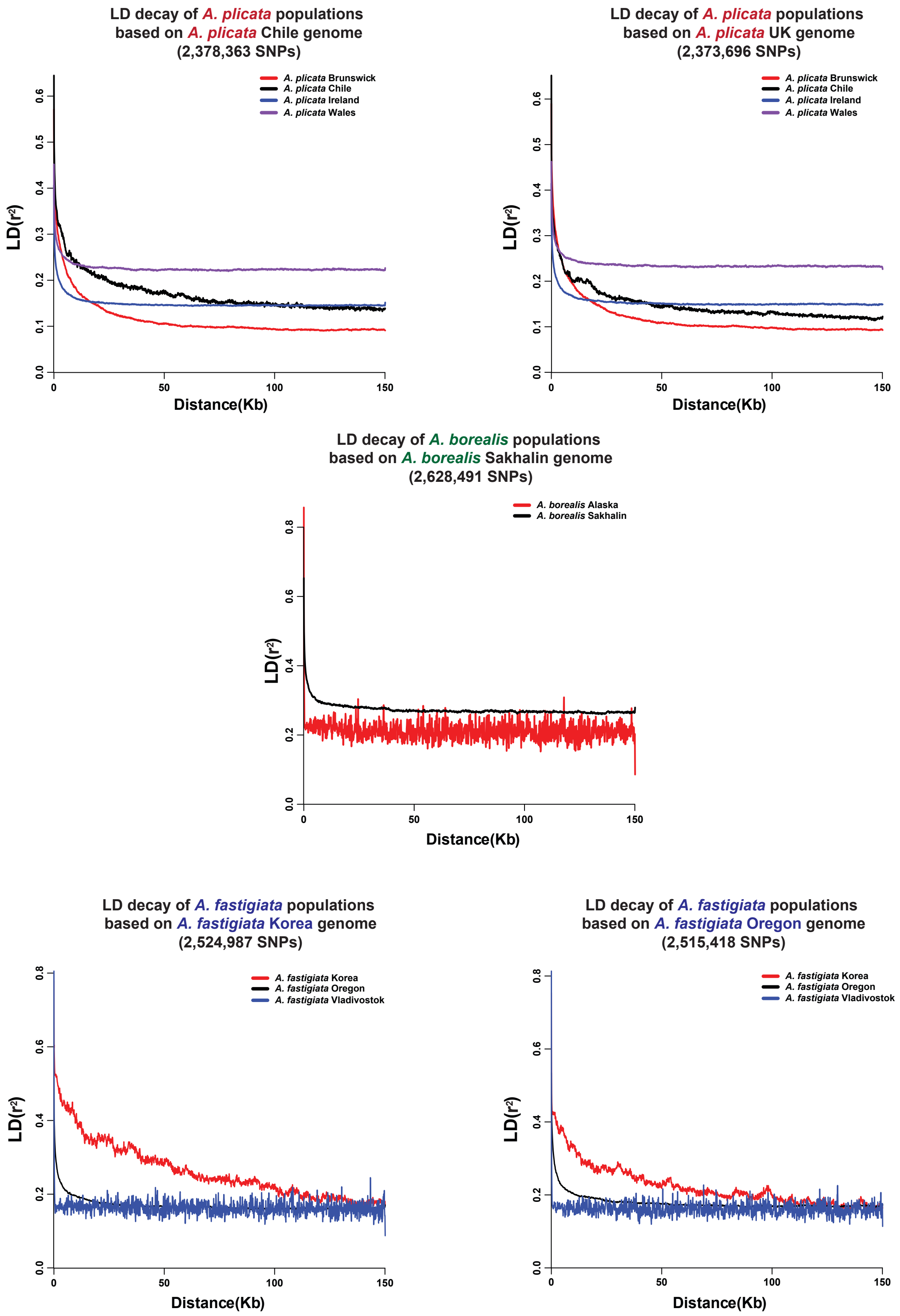

**Fig. S24. Linkage disequilibrium (LD) decay of *Ahnfeltia* populations based on each *Ahnfeltia* genome.**

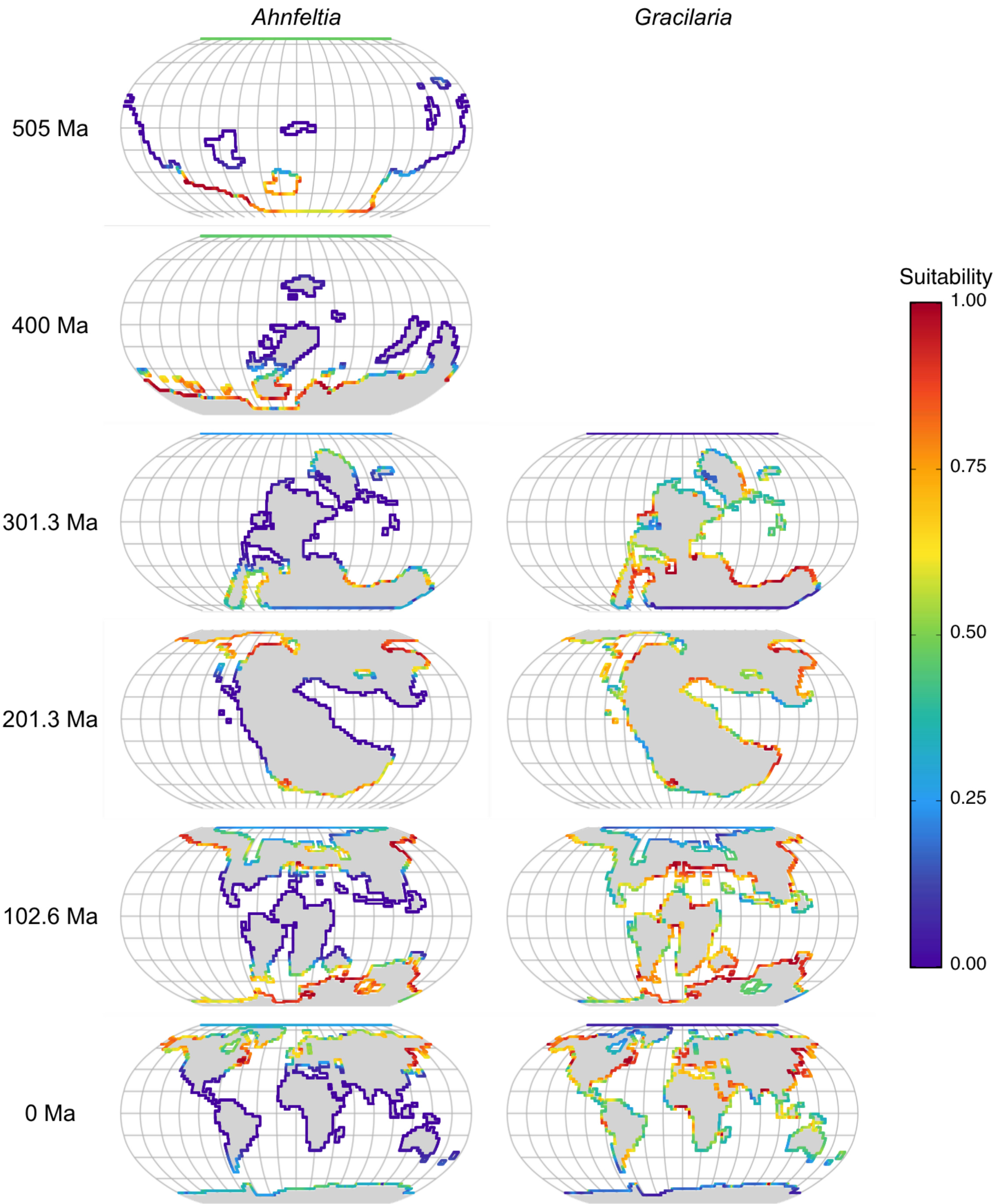

**Fig. S25. Predicted climatic suitability for *Ahnfeltia* and *Gracilaria* in coastal grid cells under the HadCM3 palaeoclimate reconstructions (using the Scotese-Foster $CO_2$ model; Valdes et al. (*85*)) at representative ~100 million-year intervals through the Phanerozoic (0, 102.6, 201.3, 301.3, 400 and 505 Ma**). Projections for *Gracilaria* are restricted to ≤300 Ma in accordance with its estimated crown lineage age. Suitability values are shown on a cloglog scale (0-1), where higher values indicate greater suitable environmental conditions for the genus. Coastal cells were enlarged for clearer visualisation but still represent suitability values from the 100-km coastal buffer (one grid cell adjacent to land).

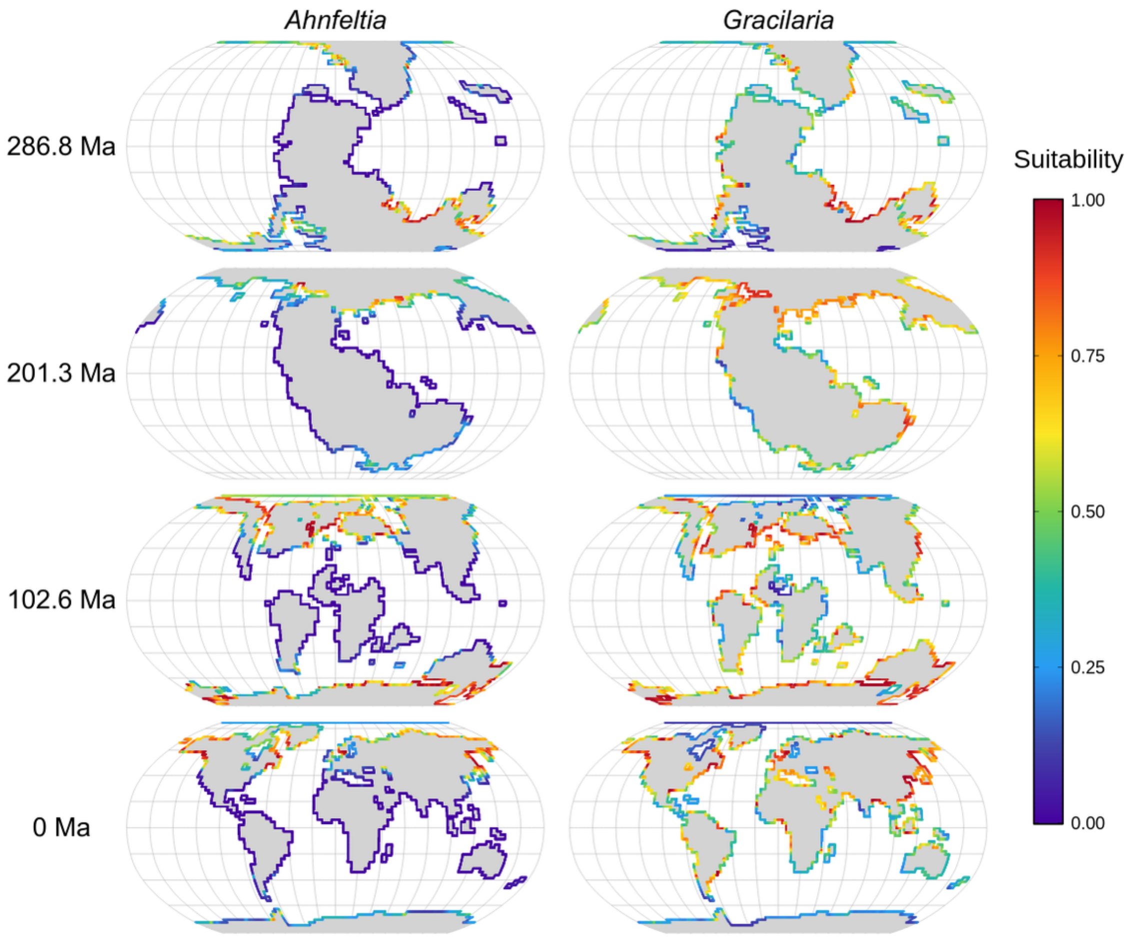

**Fig. S26. Predicted climatic suitability for *Ahnfeltia* and *Gracilaria* in coastal grid cells under the HadCM3 palaeoclimate reconstructions (using the Getech-Foster $CO_2$ model; Valdes et al. (*85*)) at representative ~100 million-year intervals (0, 102.6, 201.3 and 286.8 Ma) spanning the late Palaeozoic to the present.** Getech-Foster $CO_2$ reconstructions are available only from 286 Ma onwards and therefore omit the earliest time slices shown in other figures. Suitability values are shown on a cloglog scale (0-1), where higher values indicate greater suitable environmental conditions for the genus. Coastal cells were enlarged for clearer visualisation but still represent suitability values from the 100-km coastal buffer (one grid cell adjacent to land).

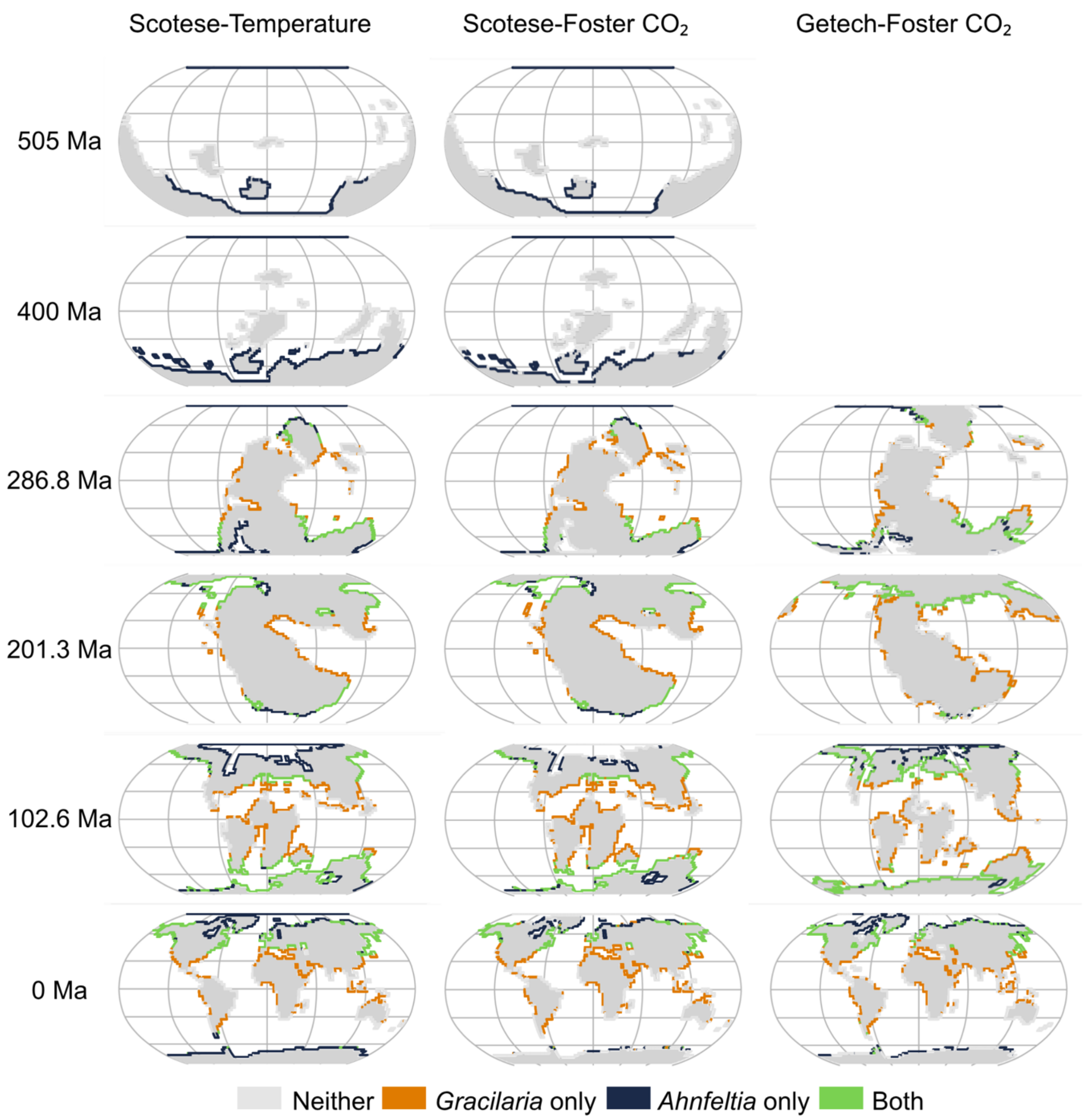

**Fig. S27. Binary climatic suitability projections for *Gracilaria* and *Ahnfeltia* under alternative palaeoclimate reconstructions across the Phanerozoic**. Projections for *Gracilaria* are restricted to ≤300 Ma in accordance with its estimated crown lineage age. Continuous suitability values were converted to presence-absence using the sensitivity-specificity threshold (see Methods). Each row represents selected projections at approximately 100 Ma intervals, and each column corresponds to one of the three palaeogeographic frameworks: Scotese-Temperature, Scotese-Foster $CO_2$, and Getech-Foster $CO_2$ (see Methods for details, Valdes et al. (*85*)). The Getech-Foster $CO_2$ reconstructions are available only for younger intervals and therefore omit the earliest time slices. Coloured cells indicate areas predicted as suitable for *Gracilaria* only (orange), *Ahnfeltia* only (navy), or both genera (green), whilst light grey cells indicate unsuitable conditions for both. Coastal cells were enlarged for clearer visualisation but still represent suitability values from the 100-km coastal buffer (one grid cell adjacent to land).

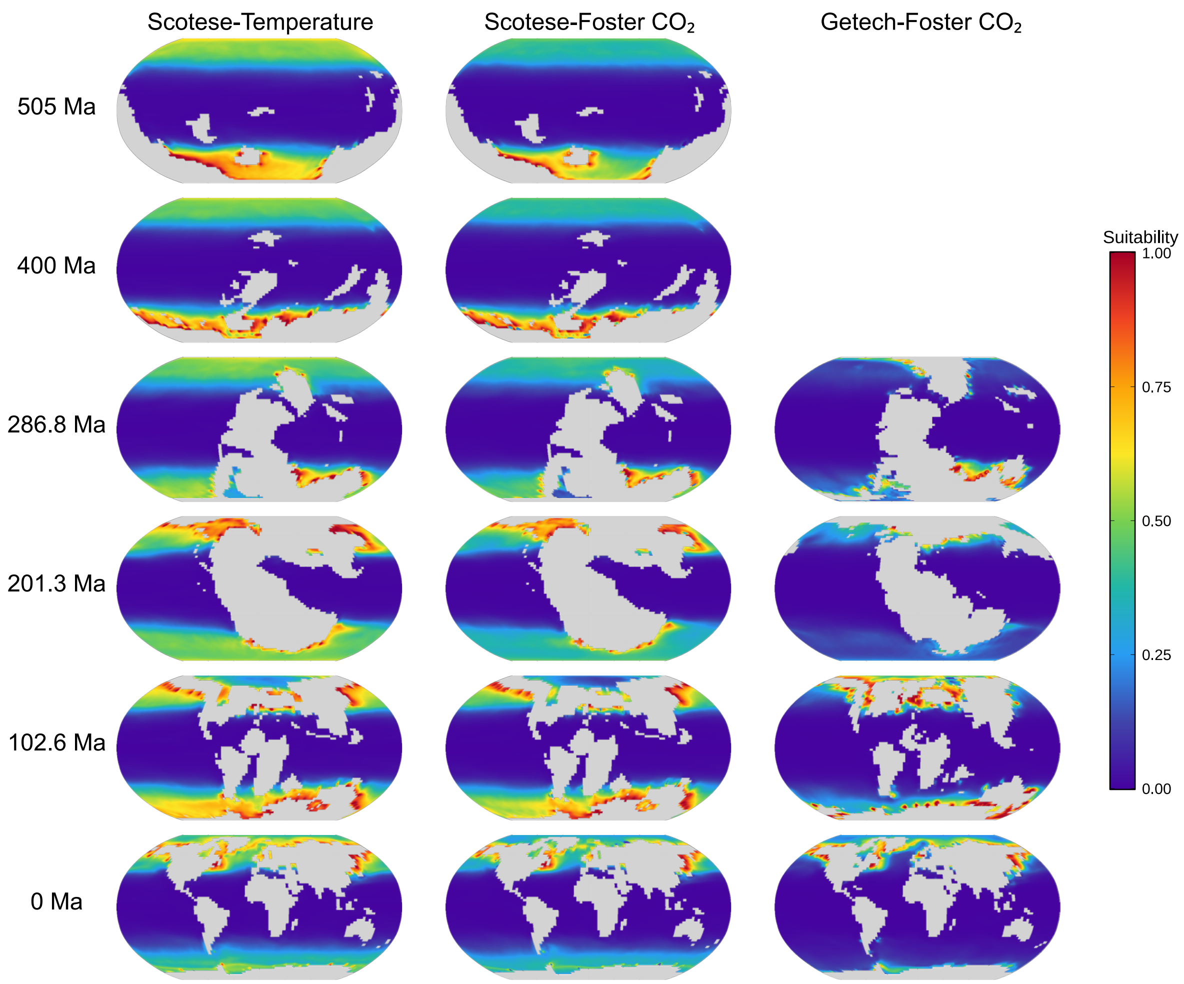

**Fig. S28. Predicted climatic suitability for *Ahnfeltia* worldwide under alternative HadCM3 palaeoclimate reconstructions (using the Scotese-Foster $CO_2$ model; Valdes et al. (*85*)) at representative ~100 million-year intervals through the Phanerozoic (0, 102.6, 201.3, 301.3, 400 and 505 Ma).** Each row represents suitability projections at approximately 100 Ma intervals, and each column corresponds to one of the three palaeogeographic frameworks: Scotese-Temperature, Scotese-Foster $CO_2$, and Getech-Foster $CO_2$ (see Methods for details, Valdes et al. (*85*)). The Getech-Foster $CO_2$ reconstructions are available only for younger intervals and therefore omit the earliest time slices. Suitability values are shown on the cloglog scale (0-1), where higher values indicate more suitable environmental conditions for the genus.

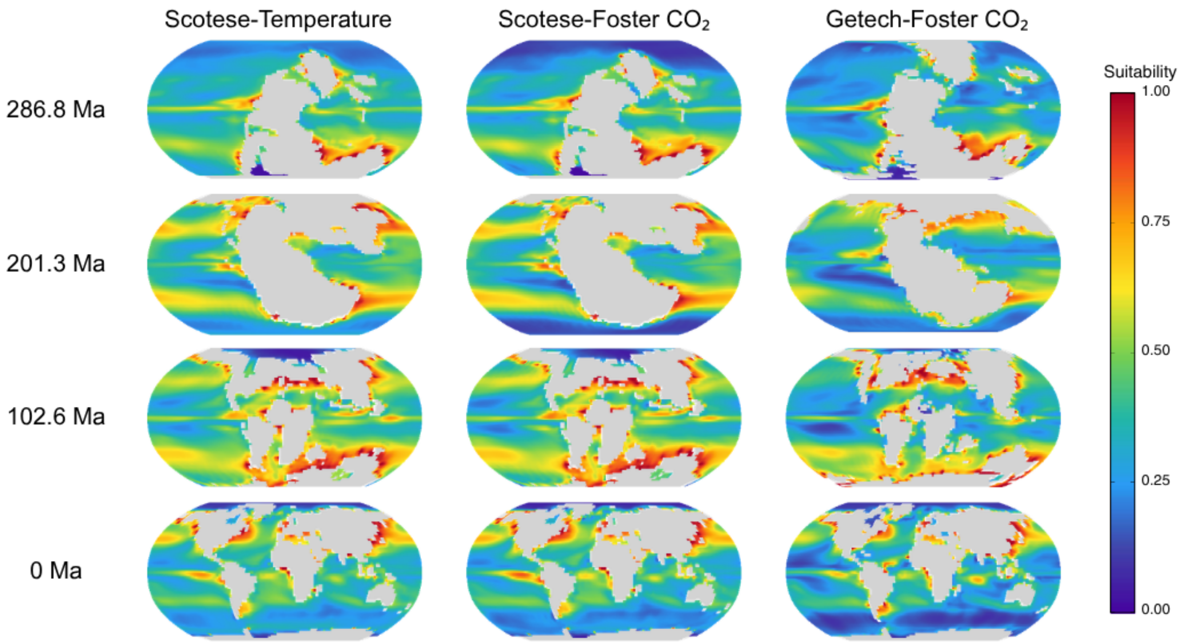

**Fig. S29. Predicted climatic suitability for *Gracilaria* worldwide under alternative HadCM3 palaeoclimate reconstructions (using the Scotese-Foster $CO_2$ model; Valdes et al. (*85*)) at representative ~100 million-year intervals through the Cenozoic to late Paleozoic (0, 102.6, 201.3, and 286.8 Ma).** Each row represents suitability projections at approximately 100 Ma intervals, and each column corresponds to one of the three palaeogeographic frameworks: Scotese-Temperature, Scotese-Foster $CO_2$, and Getech-Foster $CO_2$ (see Methods for details, Valdes et al. (*85*)). The Getech-Foster $CO_2$ reconstructions are available only for younger intervals and therefore omit the earliest time slices. Suitability values are shown on the cloglog scale (0-1), where higher values indicate more suitable environmental conditions for the genus.

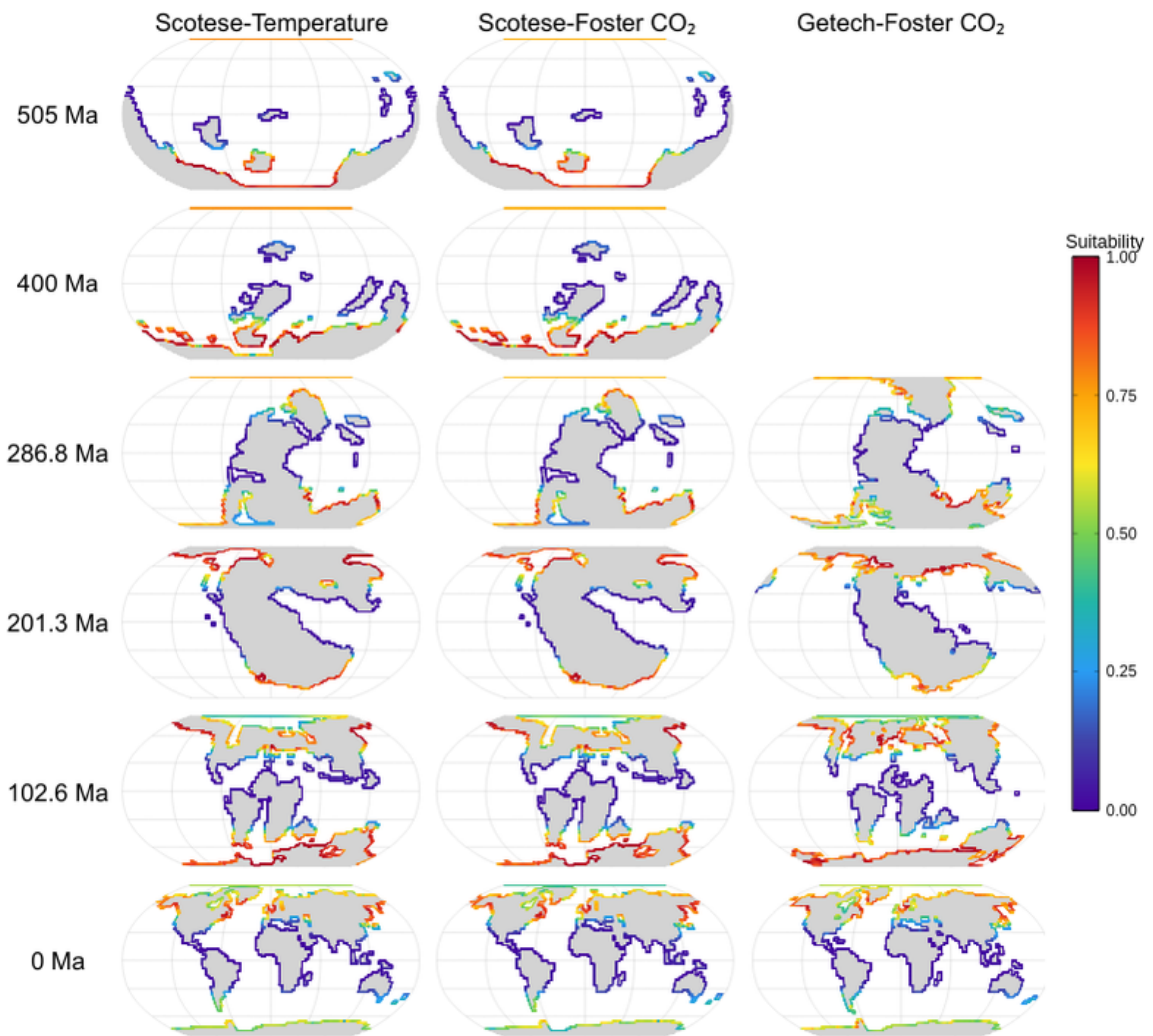

**Fig. S30. Predicted climatic suitability for *Ahnfeltia* worldwide based on the GBIF-trained model under alternative HadCM3 palaeoclimate reconstructions (using the Scotese-Foster $CO_2$ model; Valdes et al. (*85*)) at representative ~100 million-year intervals through the Phanerozoic (0, 102.6, 201.3, 301.3, 400 and 505 Ma).** The GBIF-trained model includes a larger but non-genetically verified set of occurrence records (N = 277). Each row represents suitability projections at approximately 100 Ma intervals, and each column corresponds to one of the three palaeogeographic frameworks: Scotese-Temperature, Scotese-Foster $CO_2$, and Getech-Foster $CO_2$ (see Methods for details, Valdes et al. (*85*)). The Getech-Foster $CO_2$ reconstructions are available only for younger intervals and therefore omit the earliest time slices. Suitability values are shown on the cloglog scale (0-1), where higher values indicate more suitable environmental conditions for the genus. Coastal cells were enlarged for clearer visualisation but still represent suitability values from the 100-km coastal buffer (one grid cell adjacent to land).

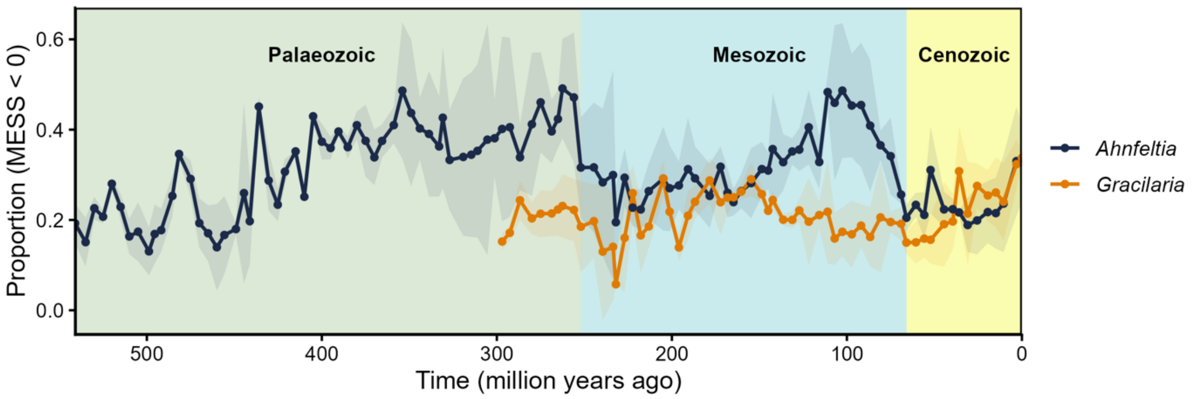

**Fig. S31. Proportion of predicted climatically suitable area exhibiting novel climatic conditions (MESS < 0) through geological time for *Ahnfeltia* and *Gracilaria*.** Lines show the mean across simulations. Shaded ribbons indicate ±1 standard deviation and background colours denote major geological eras. Higher values indicate a greater fraction of suitable habitat in extrapolated climate space and increased uncertainty in deep-time projections.

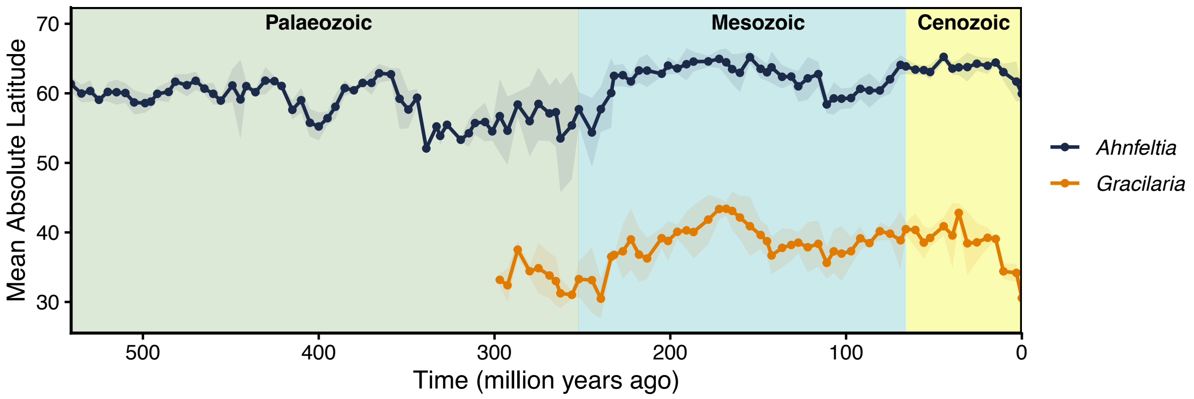

**Fig. S32. Mean absolute latitude of suitable climatic conditions through geological time for *Ahnfeltia* and *Gracilaria* after accounting for environmental extrapolation.** Lines show the mean latitude of predicted presence cells for each time slice after excluding all grid cells with negative MESS values (i.e. environments outside the range of modern training conditions), averaged across palaeoclimate simulations. Shaded ribbons indicate ±1 standard deviation and background colours denote major geological eras.

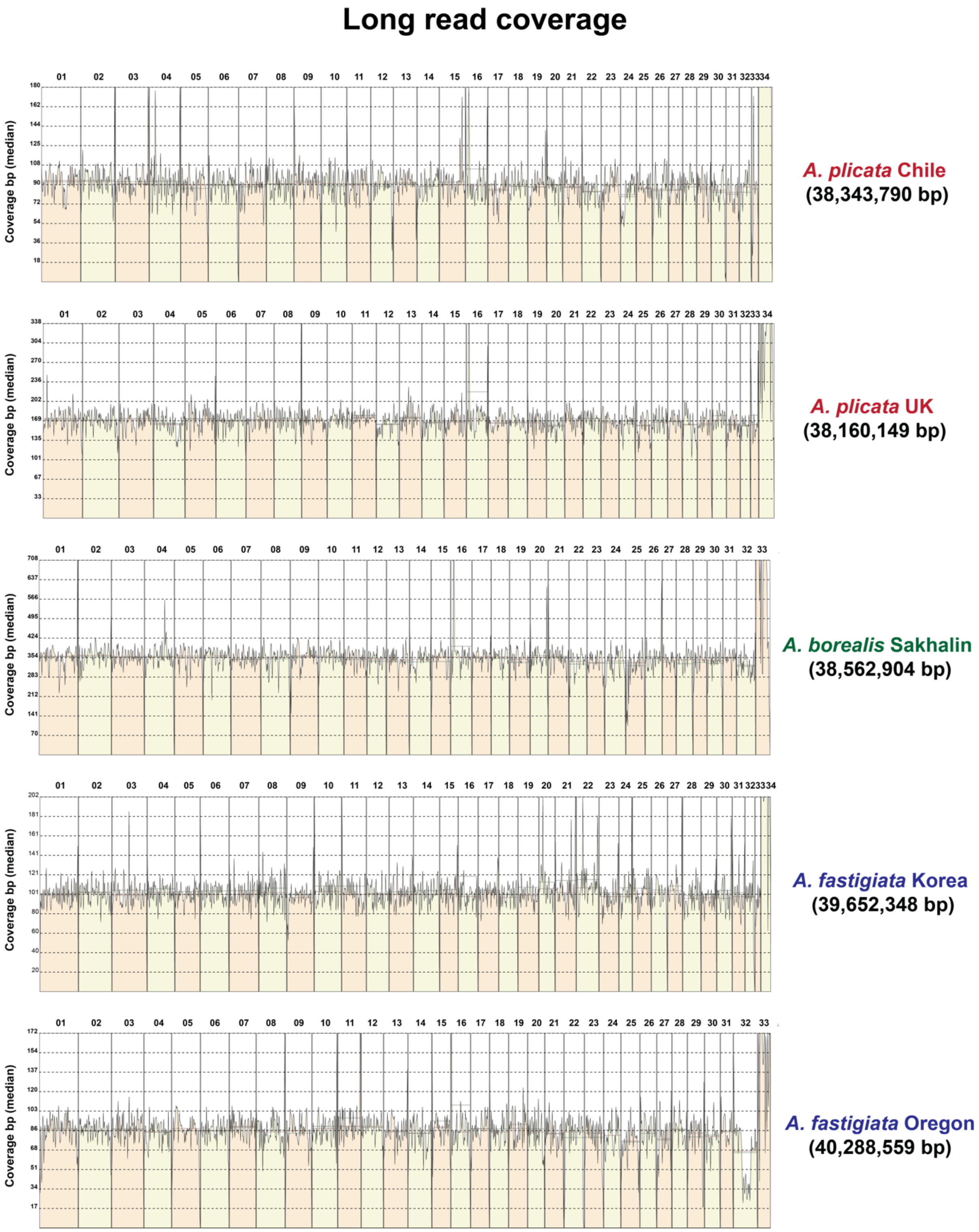

**Fig. S33. Long-read coverage of the assembled *Ahnfeltia* genomes.** This plot reflects genome completeness and assembly continuity.

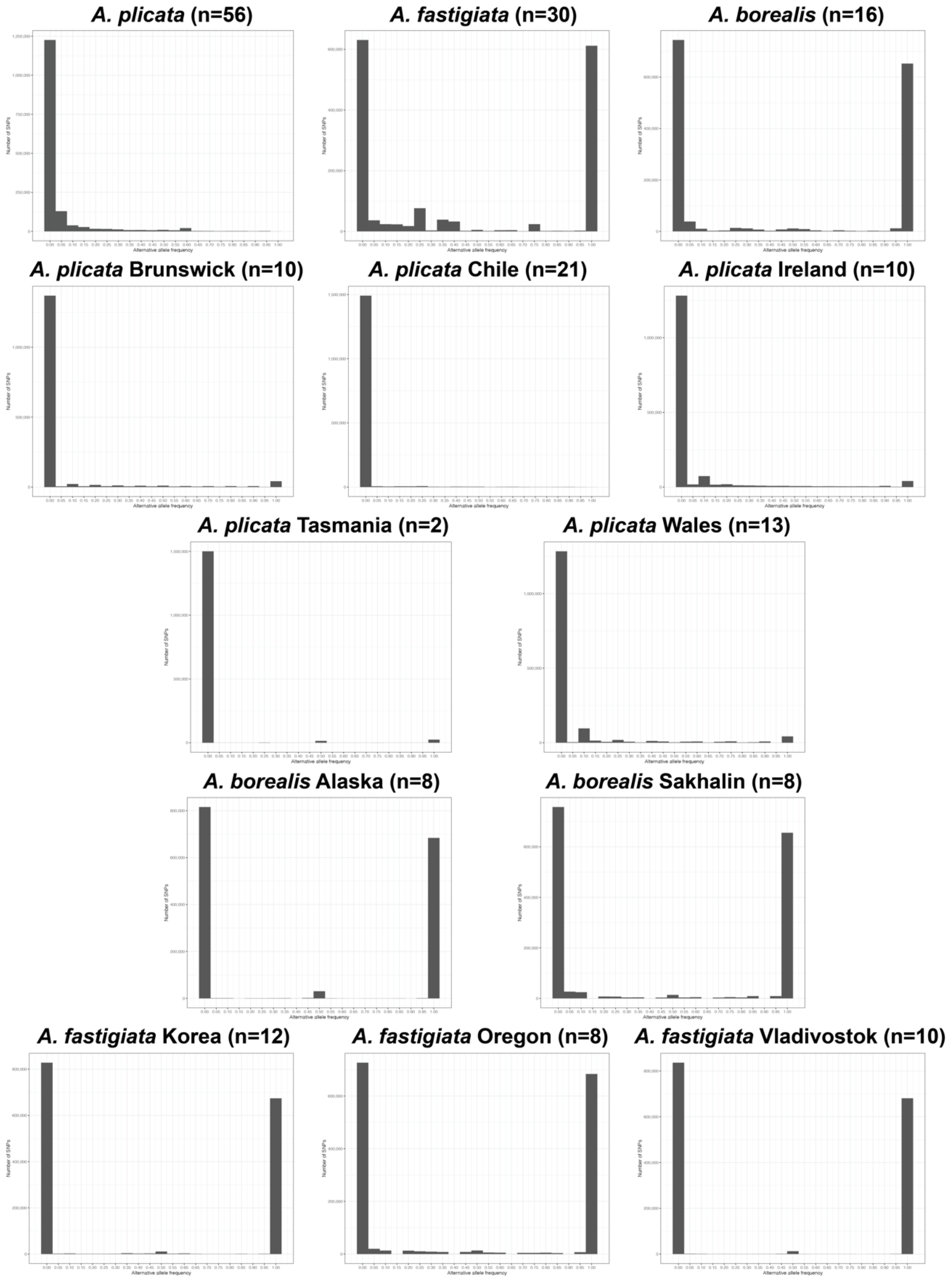

**Fig. S34. Allele frequency of *Ahnfeltia* populations based on *A. plicata* Chile genome.**

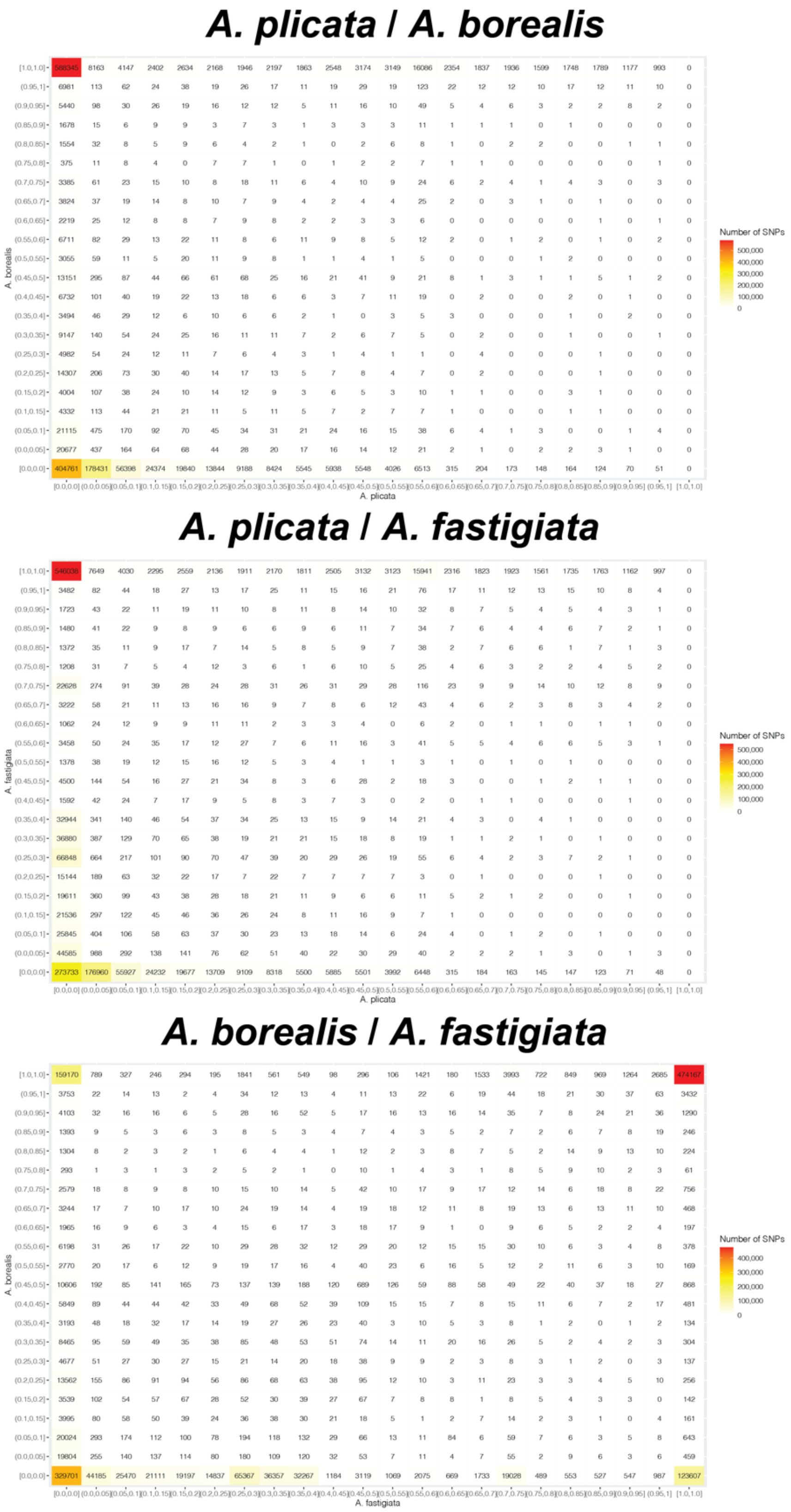

**Fig. S35. Comparison of allele frequency between *Ahnfeltia* species based on *A. plicata* Chile genome.**

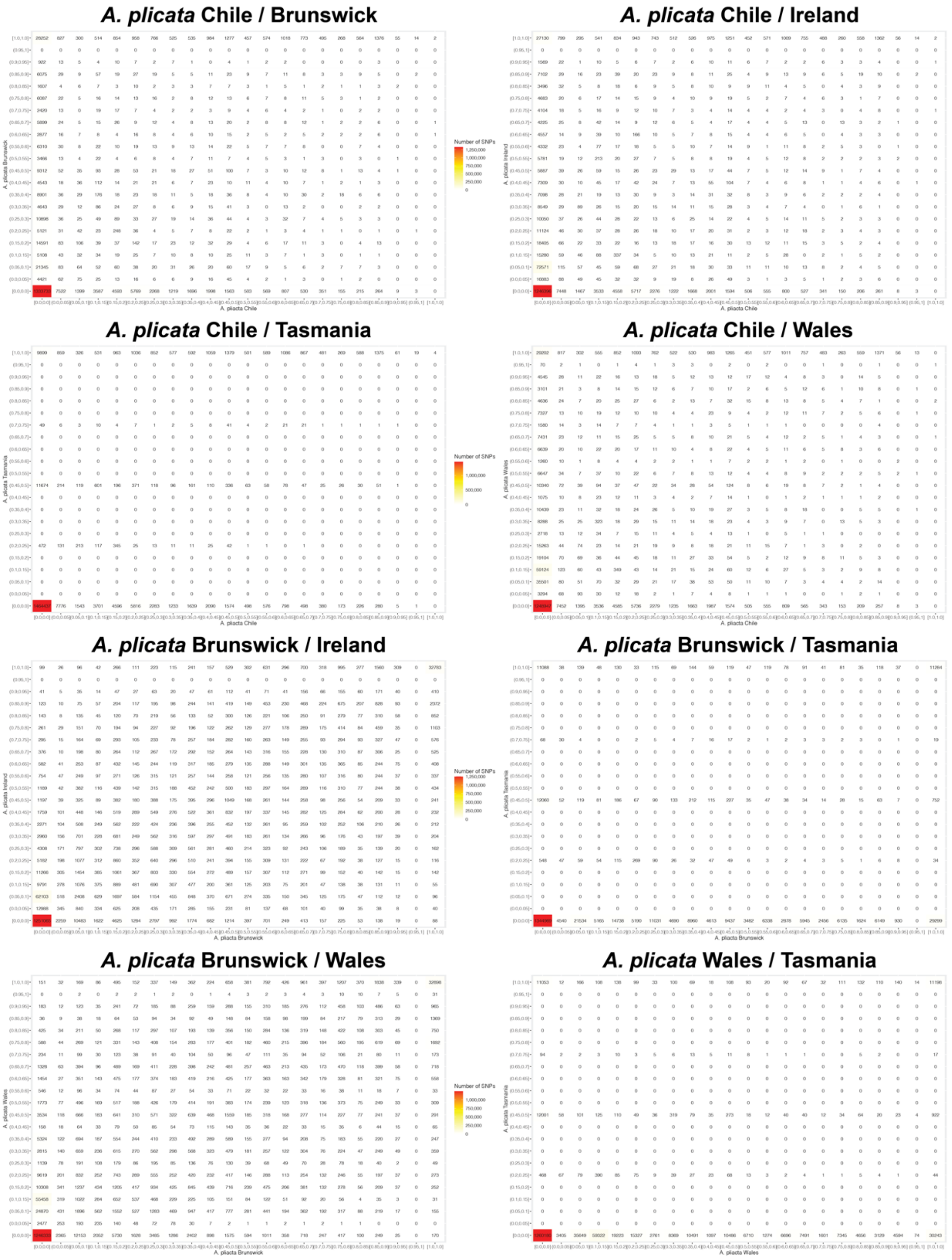

**Fig. S36. Comparison of allele frequency between *Ahnfeltia* populations based on *A. plicata* Chile genome.**

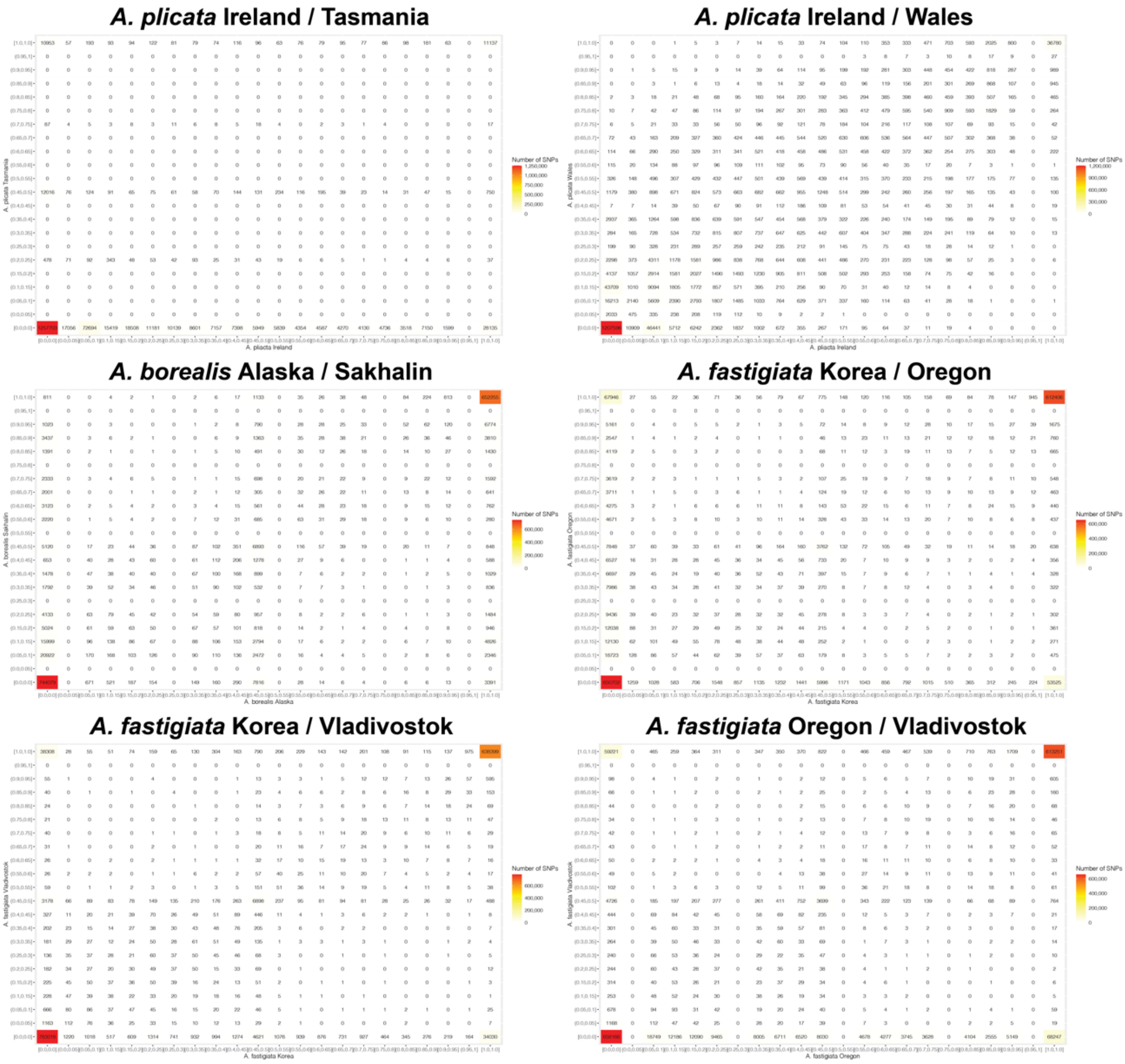

**Fig. S37. Continued.**

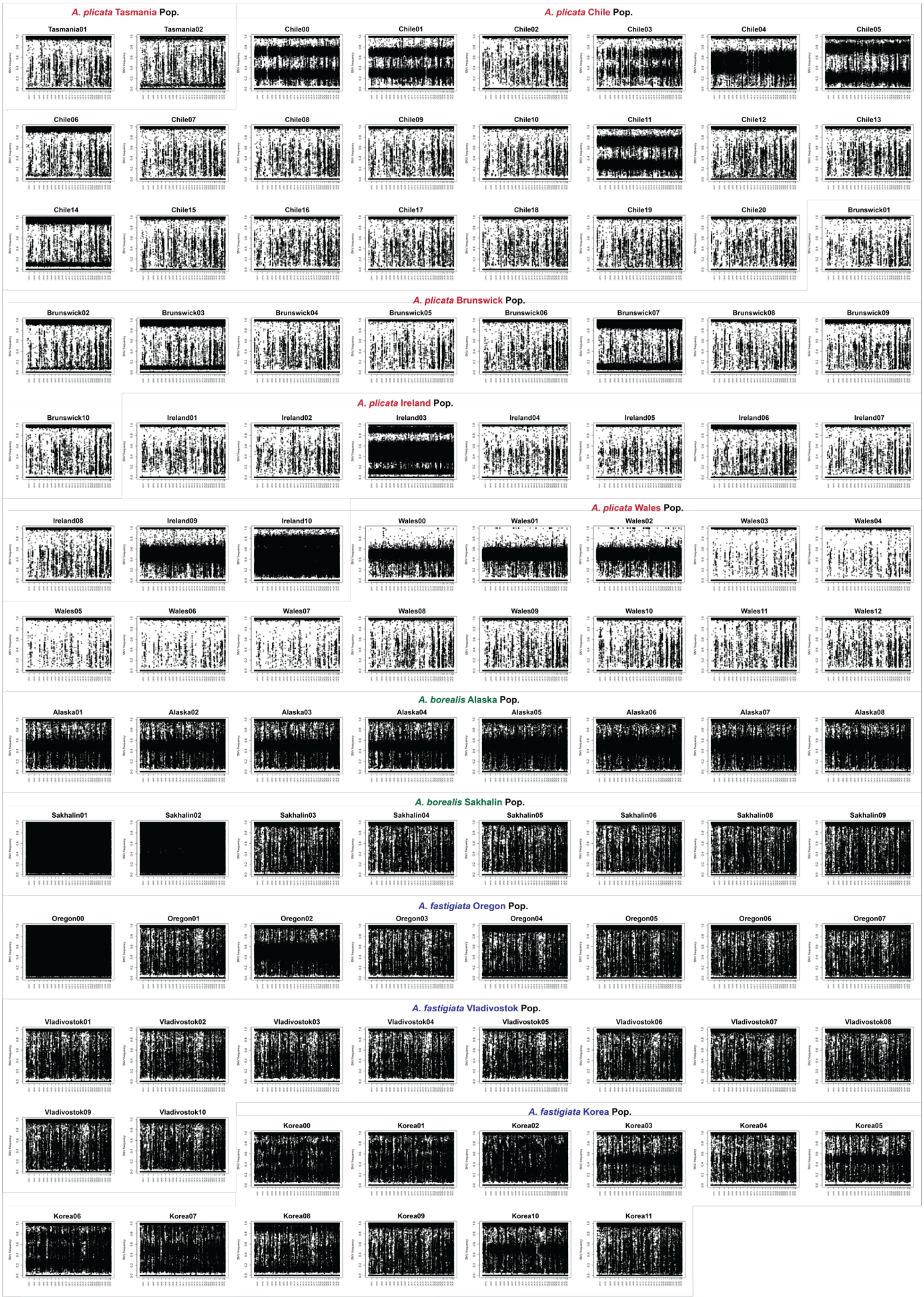

**Fig. S38. Results of the momix R package used to calculate FWS and to identify haploid and diploid signals in *Ahnfeltia* individuals.**

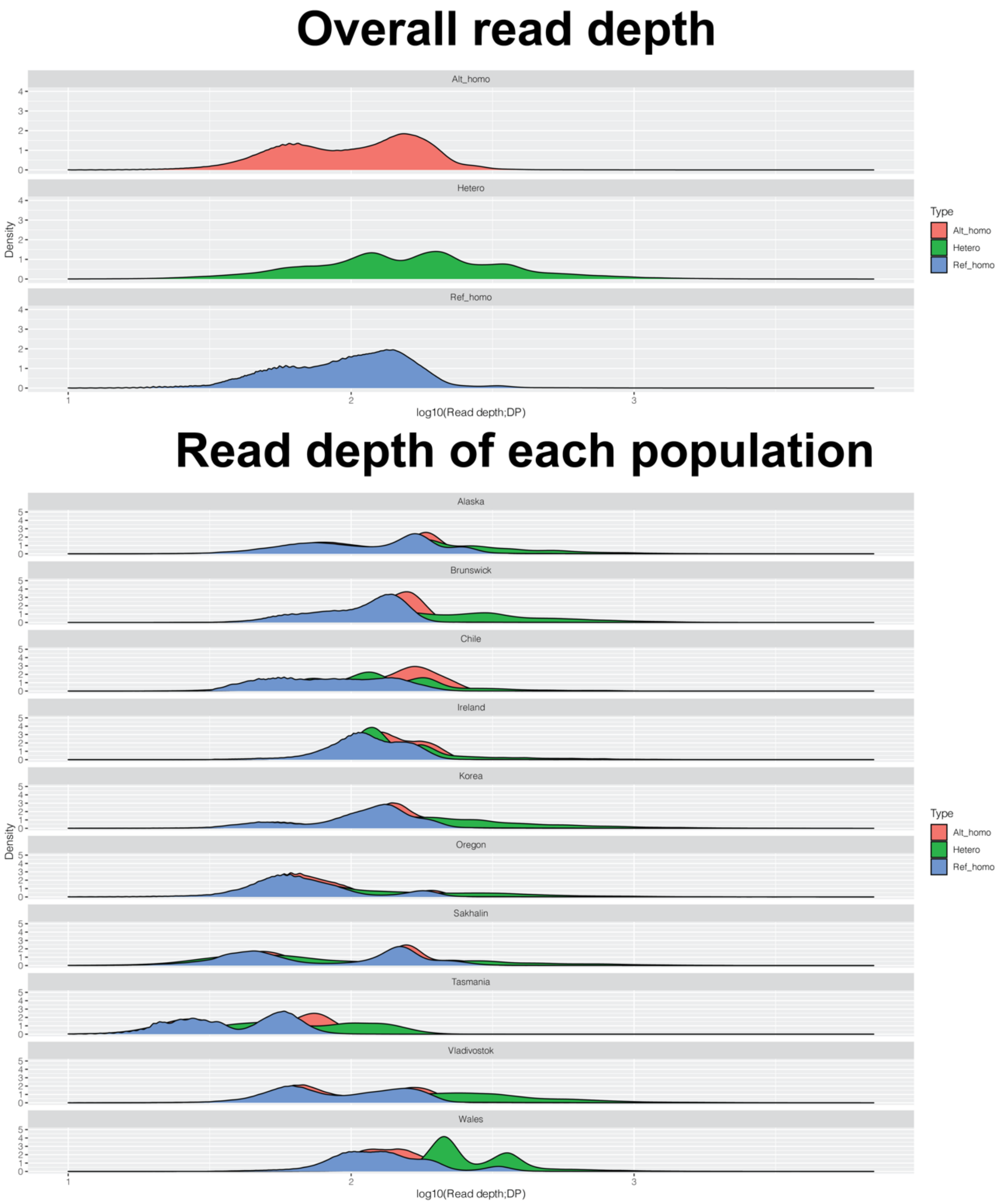

**Fig. S39. Read depth of variants depended on types of alleles and populations.**

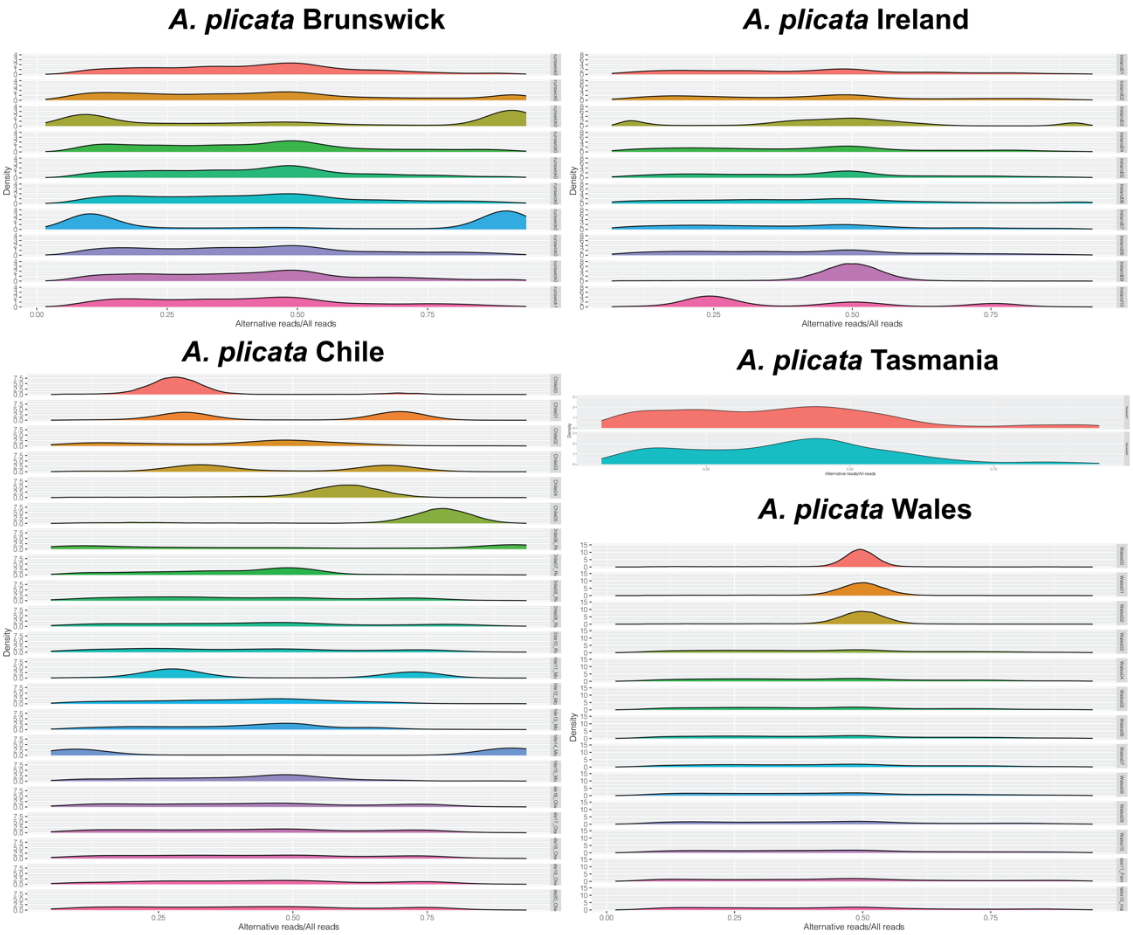

**Fig. S40. Ratio of alternative reads per all reads of *A. plicata* populations.** 0.5 value indicated heterozygous variants had exact half of alternative reads.

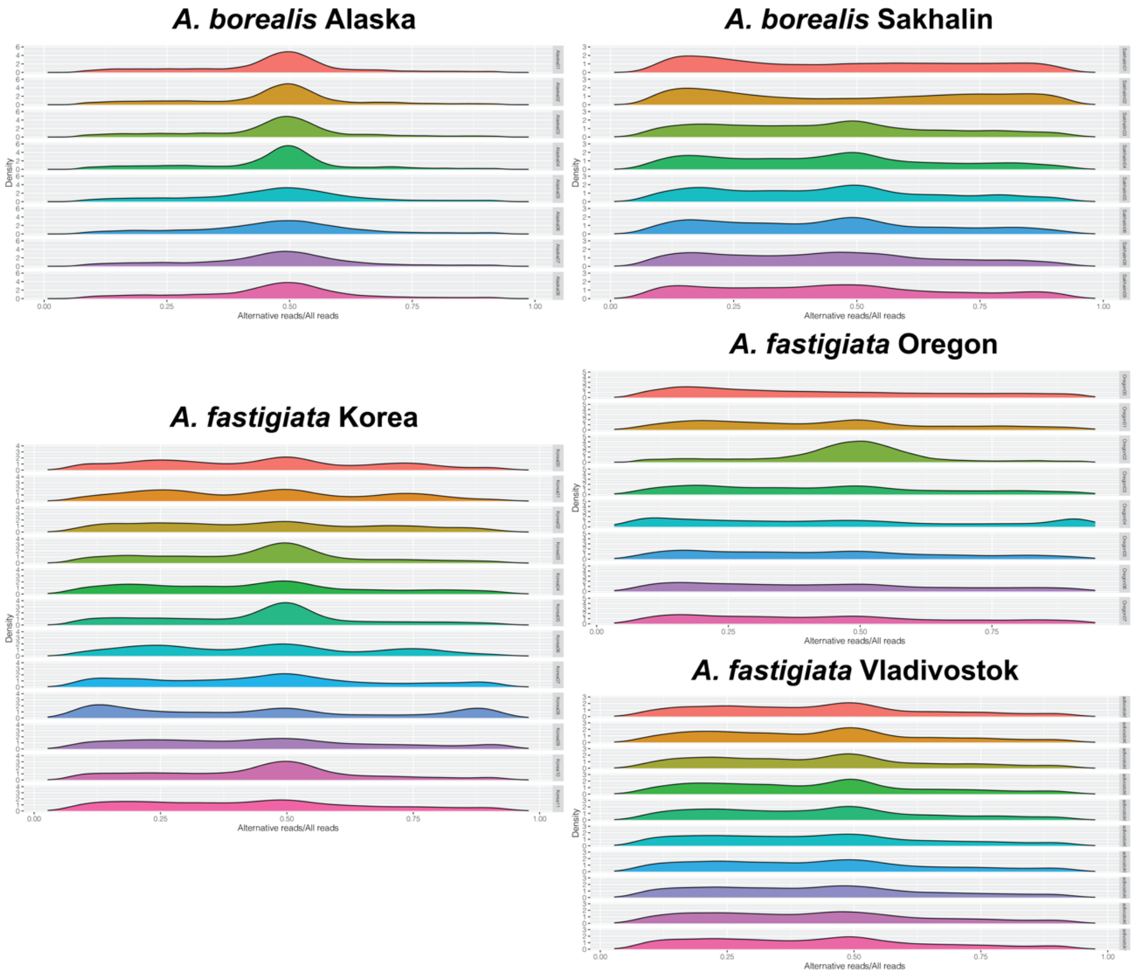

**Fig. S41. Ratio of alternative reads per all reads of *A. borealis* and *A. fastigiata* populations.**

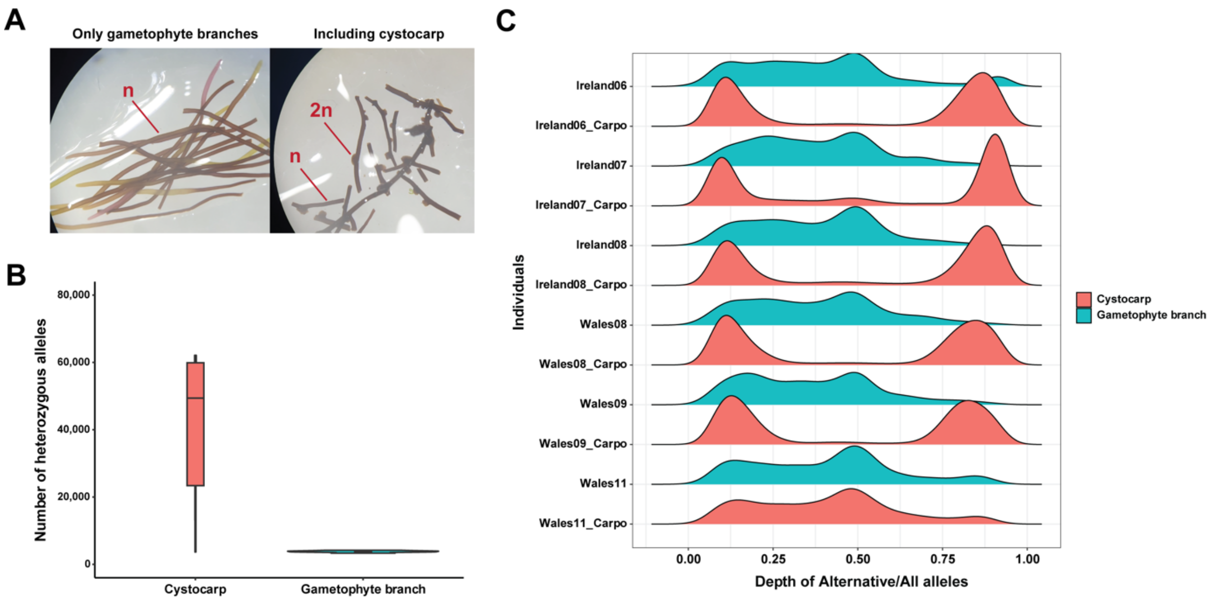

**Fig. S42. Re-examining data for clarifying types of variants from female gametophytes of *Ahnfeltia*.** (A) Photos of tested tip branches (expected as haploid) and carposporophytes (expected as diploid). (B) Number of heterozygous sites of SNPs from re-sequencing data of tip branches and carposporophytes, which indicates a higher abundance of heterozygous signals (diploid) in carposporophytes. (C) Ratio of depths of alternative/reference alleles. In purely haploid samples, allele ratios are expected to approach 0 or 1, whereas diploid samples are expected to center around 0.5. The presence of intermediate ratios indicates that diploid cells are present within gametophyte tip branches.

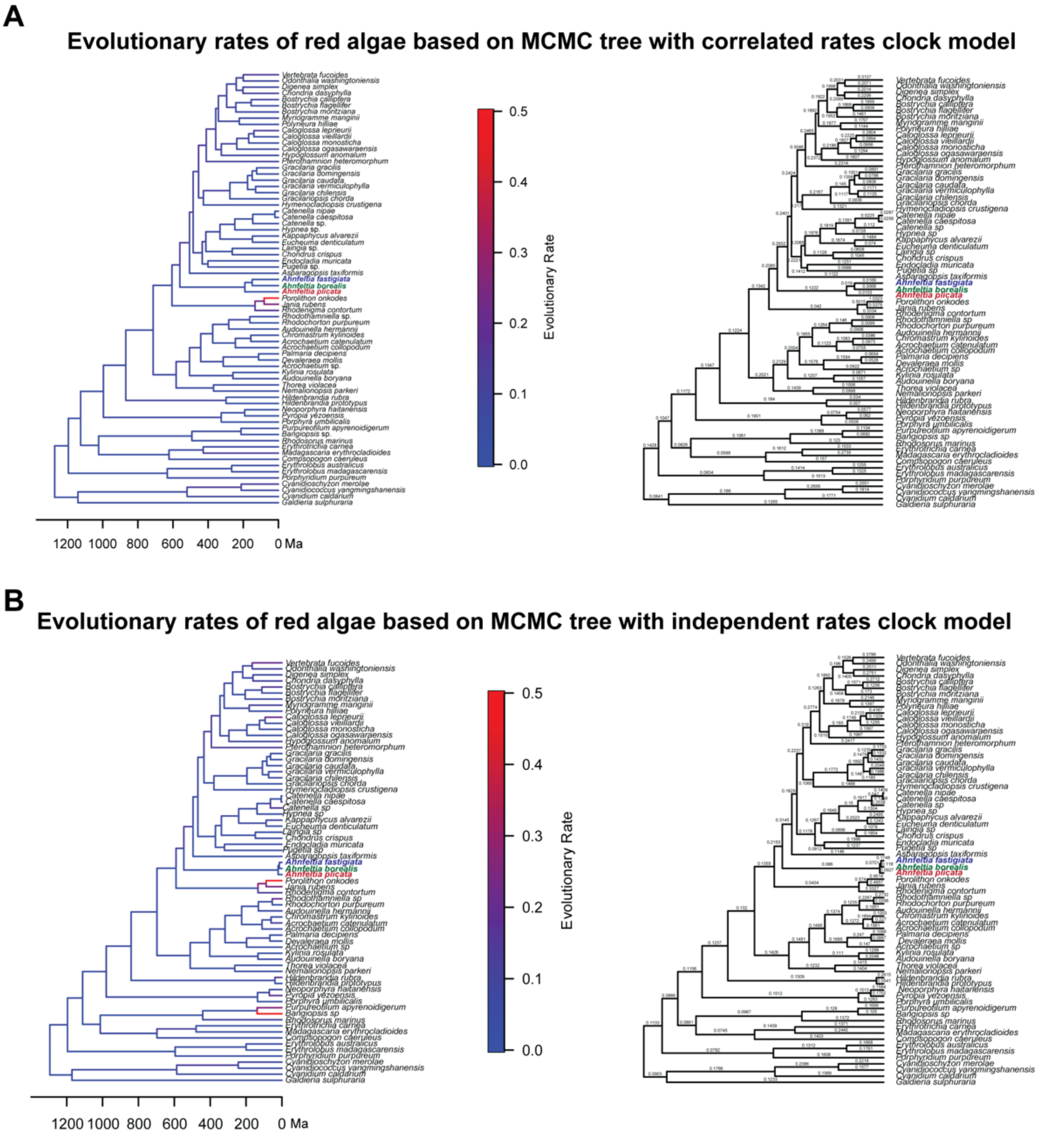

**Fig. S43. Evolutionary rates and divergence time of *Ahnfeltia* in independent and correlated models used in MCMC time estimation.** Correlated model indicates evolutionary rates of *Ahnfeltia* gradually decreased from their ancestor. However, independent model indicates evolutionary rates of *Ahnfeltia* surged within recent short period.

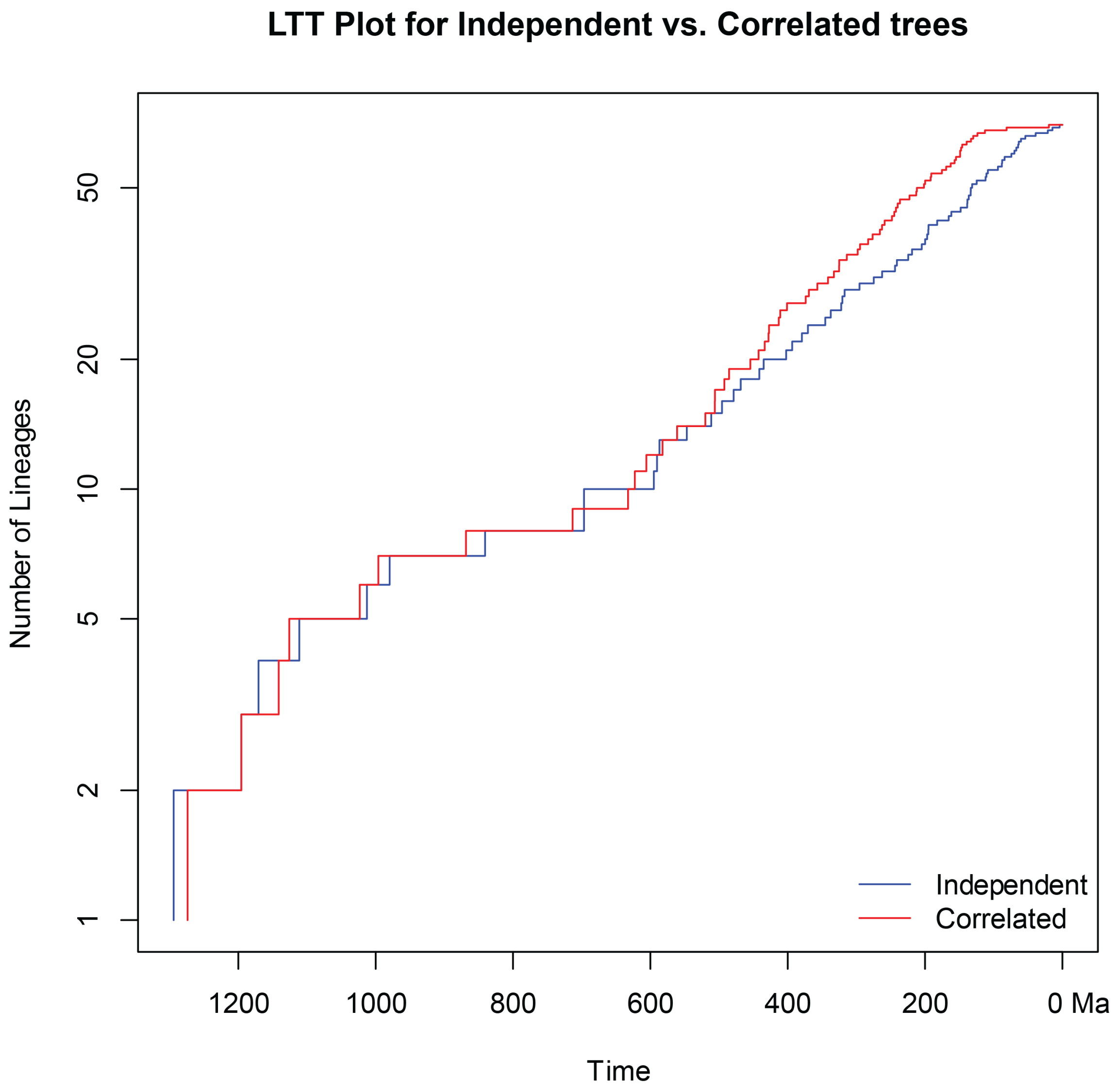

**Fig. S44. Lineage-through-time (LTT) plot to compare independent and correlated models used in MCMC time estimation**. Correlated model indicates species diversity of red algae increased in past. Independ models indicates species diversity of red algae increased recently.

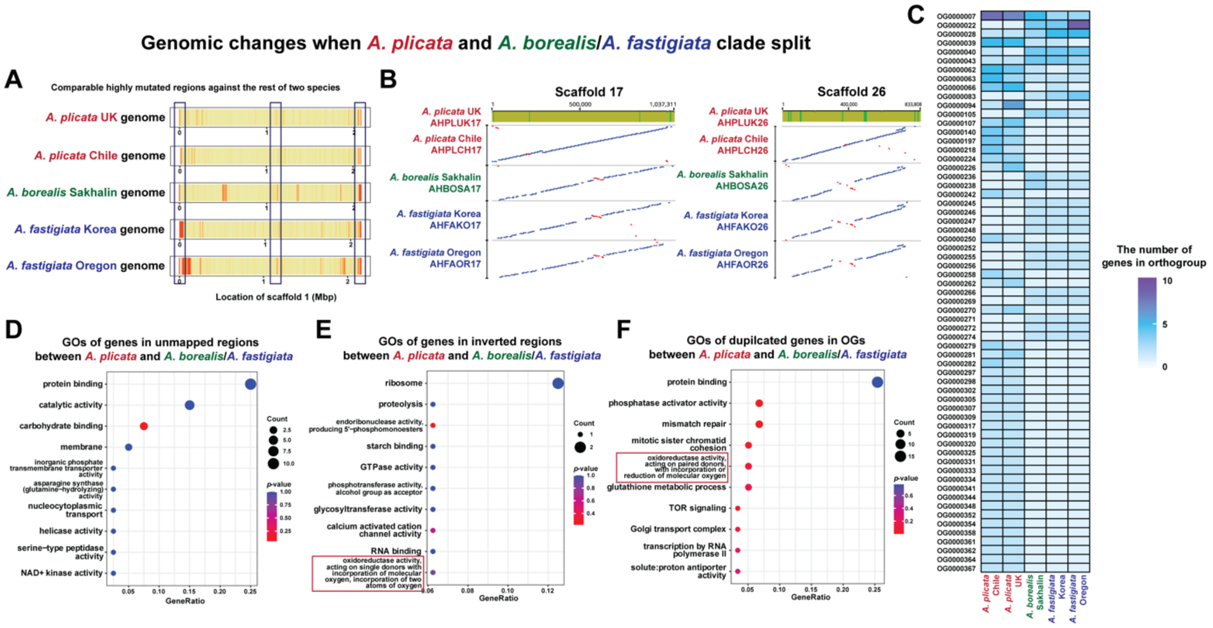

**Fig. S45. Genomic difference between *A. plicata* and *A. borealis*/*A. fastigiata* clade.** (A) Highly mutated regions between *A. plicata* and *A. borealis*/*A. fastigiata* clade verified with the unmapped regions of short read data. (B) Duplicated genes between *A. plicata* and *A. borealis*/*A. fastigiata* clade. (C) Inverted regions that occurred in *A. borealis*/*A. fastigiata* clade. (D) Gene Ontology (GO) of genes in the unmapped regions. (E) GO of duplicated genes. (F) GO of gene in inverted regions.

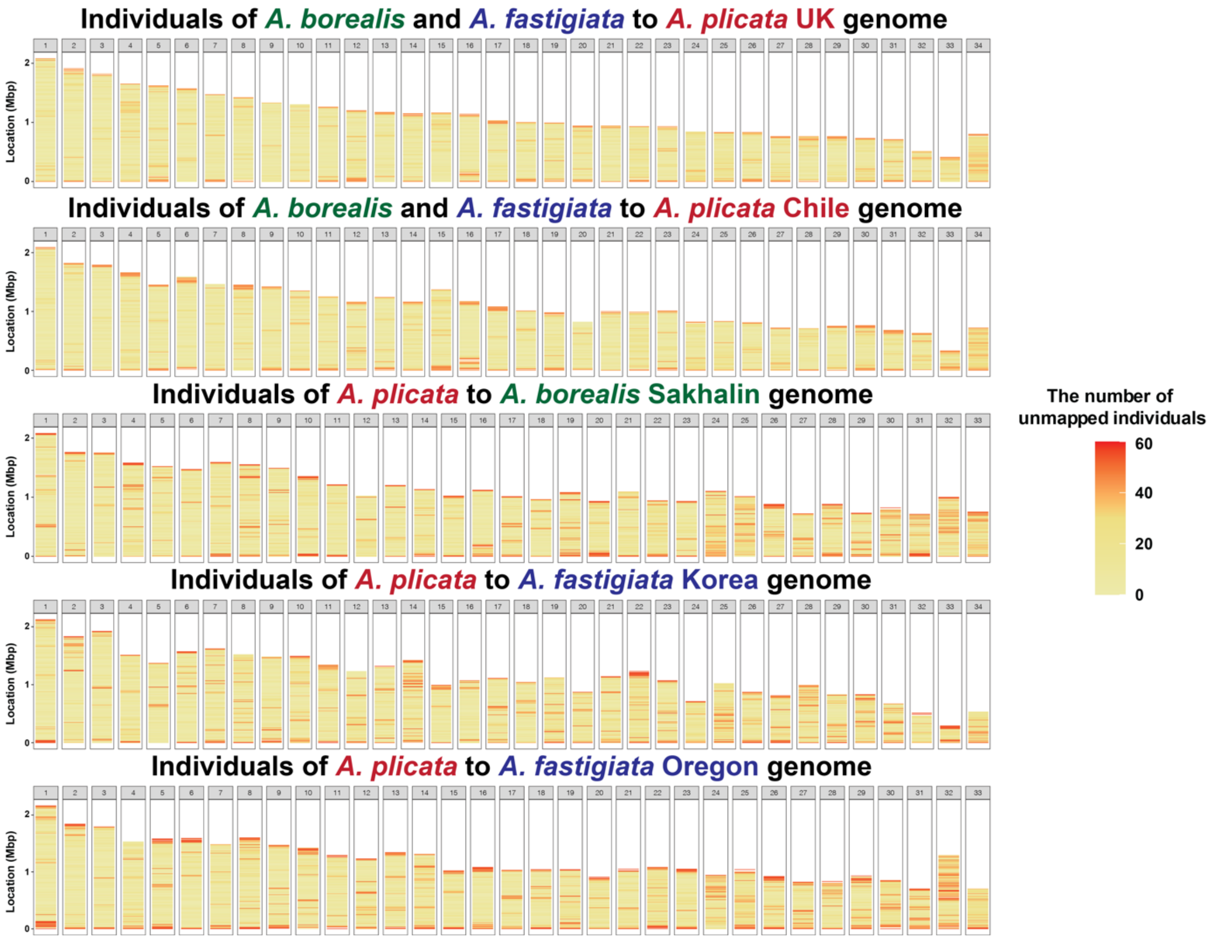

**Fig. S46. High divergence regions in chromosomes when *A. plicata* and *A. borealis*/*A. fastigiata* clades diverged.**

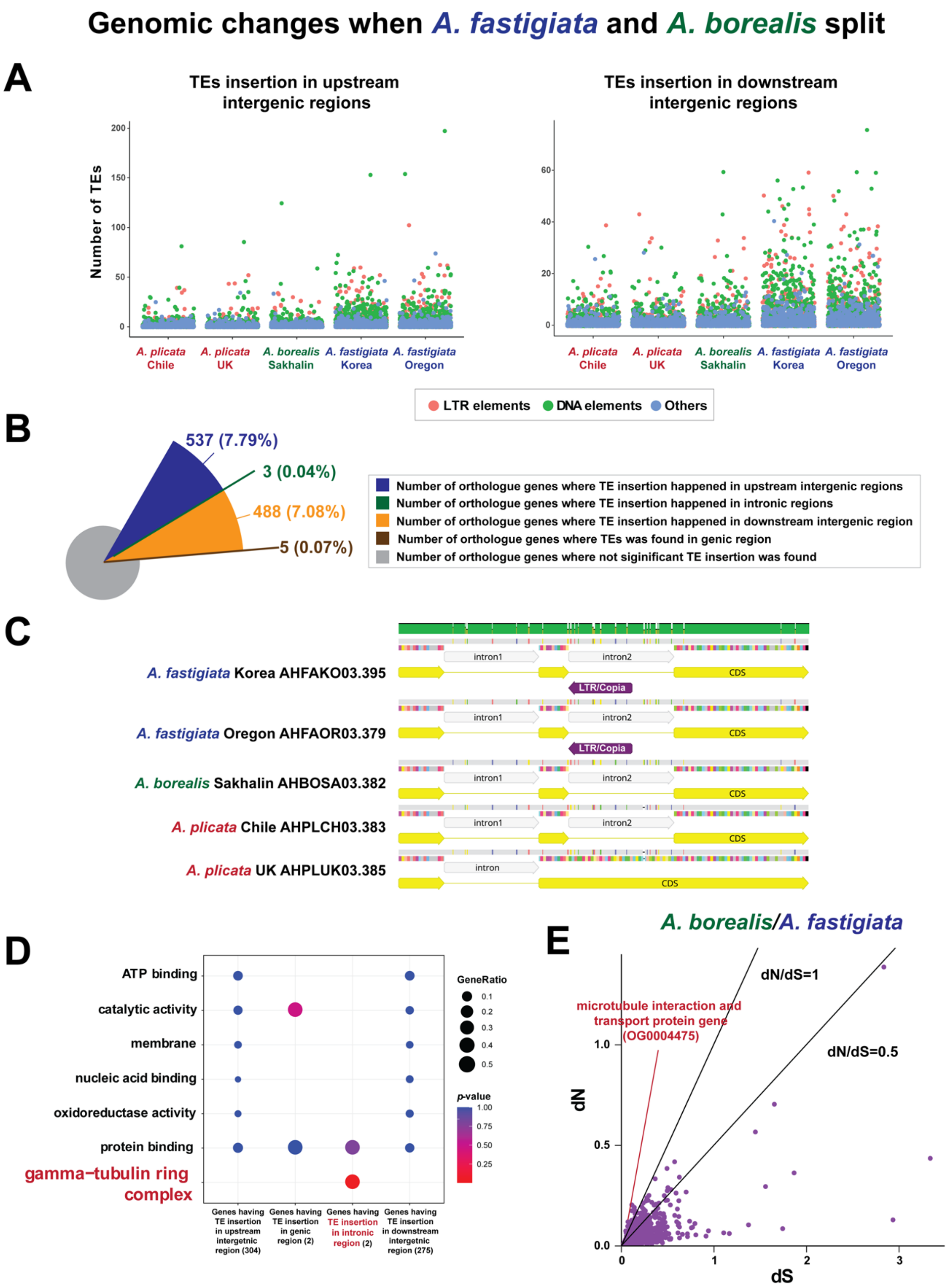

**Fig. S47. Key genomic and genetic differences associated with the divergence of *A. fastigiata*.** (A) Distribution of TE insertion sites in upstream and downstream intergenic regions of orthologue genes. (B) Number of TE insertions surrounding comparable orthologous genes across *Ahnfeltia* genomes. (C) Insertion of an LTR Copia element within a gene associated with the γ-tubulin ring complex. (D) Gene Ontology (GO) of orthologue genes where TE insertion occurred. (E) dN/dS analysis indicating positive selection in genes involved in microtubule interaction and transport.

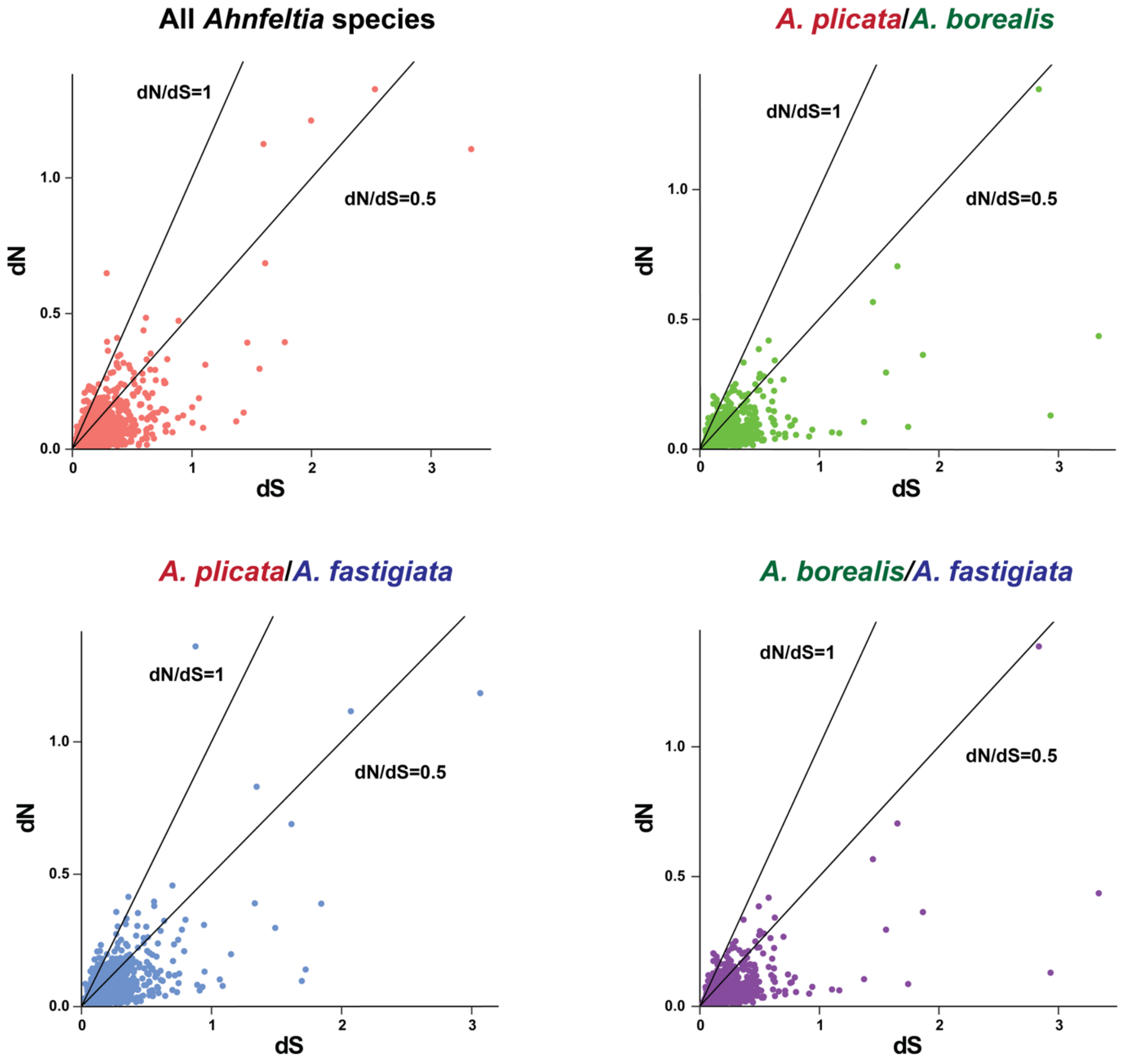

**Fig. S48. dN/dS of comparable orthologue genes (6,897) of *Ahnfeltia* species.**

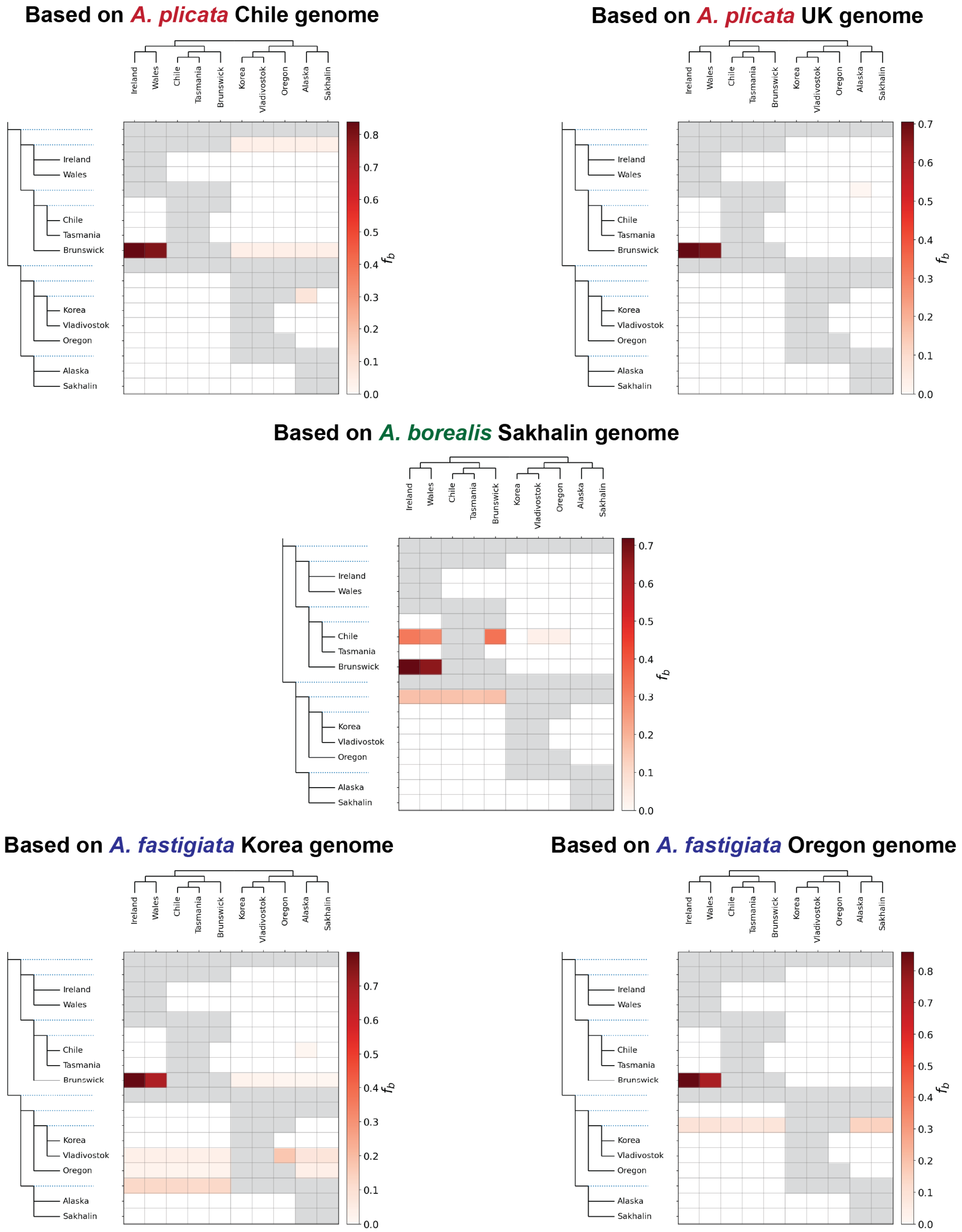

**Fig. S49. D-statistics and populational admixture test using Dsuite.** Limited gene flow was detected among populations; however, relatively strong gene flow from the Brunswick to the Ireland and Wales populations was observed, as indicated by high *f* values.

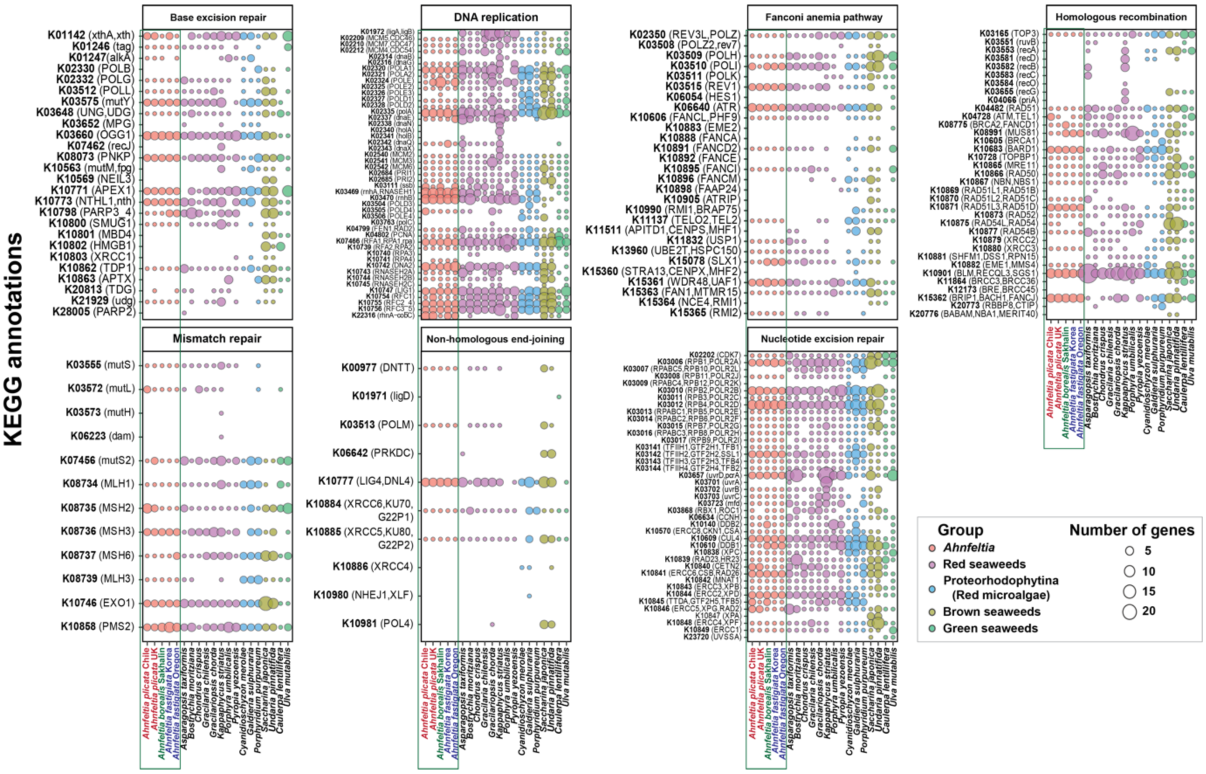

**Fig. S50. DNA repair systems of *Ahnfeltia* and other seaweeds were examined using the full set of DNA replication and repair pathways provided by KEGG (approximately 360 genes).** These genes were conserved in *Ahnfeltia* genomes, with no evidence of significant gene duplication or insertion.

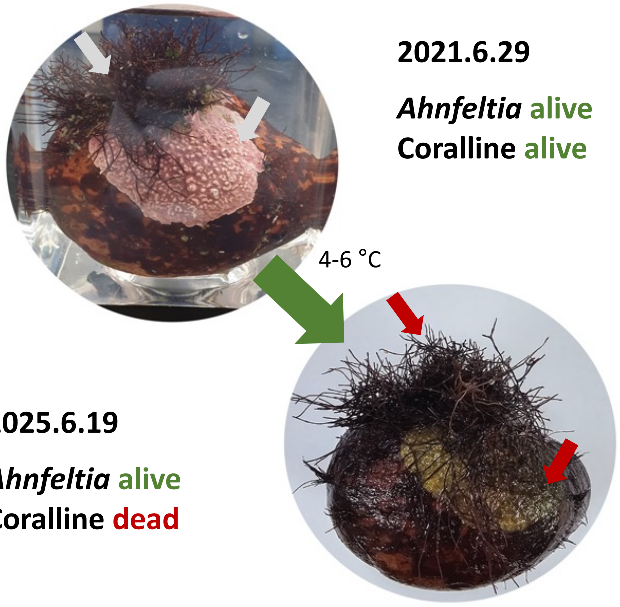

**Fig. S51. Simple experimental results demonstrating the persistence of *Ahnfeltia*.** *Ahnfeltia* and coralline algae were stored at 4 °C for four years with two seawater changes. During this period, the coralline algae died, whereas *Ahnfeltia* survived.

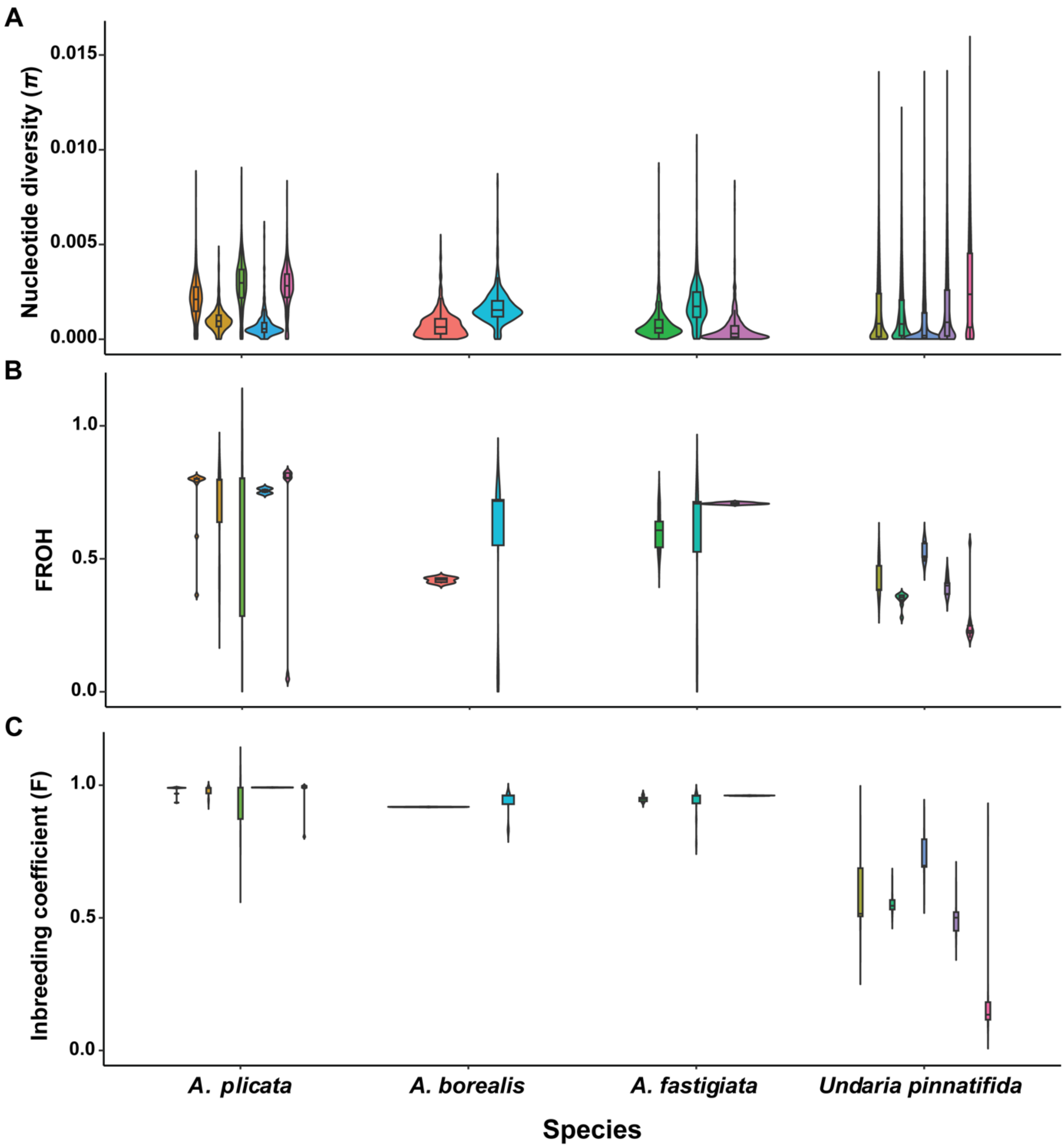

**Fig. S52. Comparison of nucleotide diversity (π) and inbreeding levels between *Ahnfeltia* species and *Undaria pinnatifida*.** (A) Differences in nucleotide diversity (π) between *Ahnfeltia* and *U. pinnatifida.* (B) Proportion of the genome in runs of homozygosity, used to estimate the inbreeding coefficient based on ROH (FROH), comparing *Ahnfeltia* and *U. pinnatifida*. (C) Inbreeding coefficient (F) of each *Ahnfeltia* population compared with that of *U. pinnatifida.*

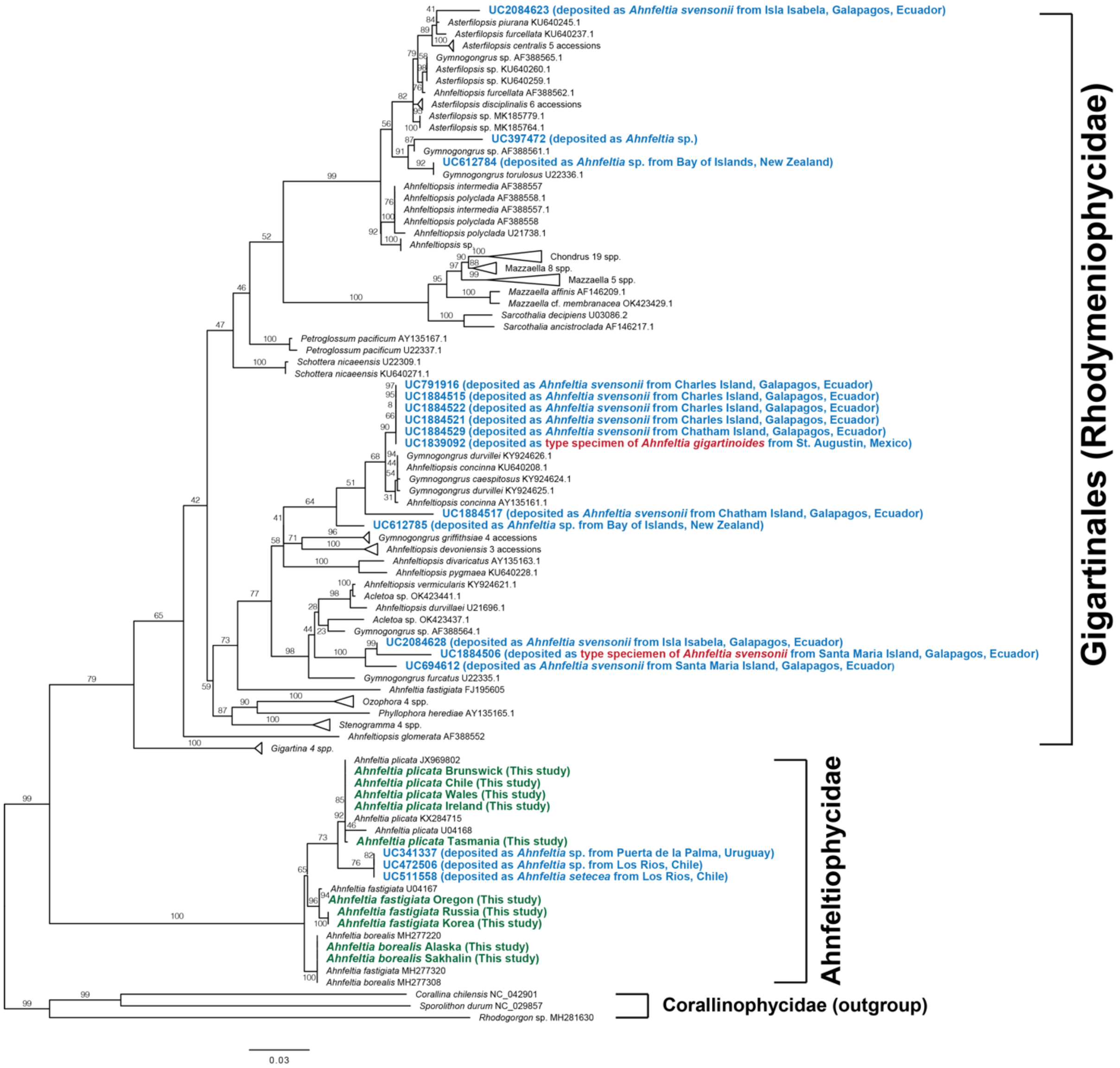

**Fig. S53. Re-examination of historical *Ahnfeltia* voucher specimens from the Berkeley Herbarium.** Most samples, including the type specimens of *A. gigartinoides* and *A. svensonii*, were identified as species belonging to the order Gigartinales. Only the samples from Uruguay and Chile were identified as *A. plicata*.

**Supplementary Table 1. Monthly palaeoenvironmental variables extracted from palaeoclimate reconstructions**. These variables were used to derive summary statistics (mean, minimum, maximum, seasonal range, and standard deviation) for niche modelling of *Ahnfeltia* and *Gracilaria*.

| Variable | Short name | Unit | Rationale | Reference |
|---|---|---|---|---|
| **Sea surface temperature** | SST | °C | Represents the thermal conditions that shape the metabolic performance, reproduction, and distributional limits of marine macroalgae. | Chefaoui et al. (*176*) |
| **Salinity** | Salinity | PSU | Reflects the osmotic environment that influences physiological tolerance and restricts red algae species to coastal areas with suitable salinity regimes. | Nejrup and Pedersen (*177*) |
| **Photosynthetically active radiation** | PAR | mol photons $m^{-2}$ $d^{-1}$ | A proxy for light availability, which is a key driver of photosynthesis, growth, and depth limits in red algae. | Raven and Hurd (*178*) |

**Supplementary Table 2. Summary of occurrence data filtering and ENM performance for *Ahnfeltia* and *Gracilaria***. The occurrence record columns show the number of records before and after quality control and spatial filtering. Values in the cross-validation (CV) columns represent the mean and standard deviation (in parentheses) of the area under the receiver operating characteristic curve (AUC) from spatial and environmental cross-validation. The final columns report AUC and Continuous Boyce Index (CBI) scores for the full models trained on all post-filtering occurrence records.

| | | **Occurrence Records** | | **CV** | | **Final Model** | |
|---|---|---|---|---|---|---|---|
| **Dataset Type** | **Genus** | **Raw** | **Processed** | **Spatial** | **Enviro.** | **AUC** | **CBI** |
| **Genetically verified** | *Ahnfeltia* | 342 | 39 | 0.87 (0.08) | 0.76 (0.14) | 0.92 (0.01) | 0.93 (0.03) |
| | *Gracilaria* | 419 | 81 | 0.68 (0.12) | 0.67 (0.24) | 0.73 (0.02) | 0.96 (0.02) |
| **GBIF** | *Ahnfeltia* | 5702 | 277 | 0.82 (0.07) | 0.72 (0.32) | 0.88 (0.01) | 0.97 (0.01) |

51. E. Derelle, C. Ferraz, S. Rombauts, P. Rouzé, A. Z. Worden, S. Robbens, F. Partensky, S. Degroeve, S. Echeynié, R. Cooke, Y. Saeys, J. Wuyts, K. Jabbari, C. Bowler, O. Panaud, B. Piégu, S. G. Ball, J.-P. Ral, F.-Y. Bouget, G. Piganeau, B. De Baets, A. Picard, M. Delseny, J. Demaille, Y. Van de Peer, H. Moreau, Genome analysis of the smallest free-living eukaryote *Ostreococcus tauri* unveils many unique

features. *Proc. Natl. Acad. Sci. U.S.A.* **103**, 11647-11652 (2006). doi:10.1073/pnas.0604795103

52. T. Sasaki, P. International Rice Genome Sequencing, The map-based sequence of the rice genome. *Nature* **436**, 793-800 (2005). doi:10.1038/nature03895

53. S. S. Merchant, S. E. Prochnik, O. Vallon, E. H. Harris, S. J. Karpowicz, G. B. Witman, A. Terry, A. Salamov, L. K. Fritz-Laylin, L. Maréchal-Drouard, W. F. Marshall, L.-H. Qu, D. R. Nelson, A. A. Sanderfoot, M. H. Spalding, V. V. Kapitonov, Q. Ren, P. Ferris, E. Lindquist, H. Shapiro, S. M. Lucas, J. Grimwood, J. Schmutz, P. Cardol, H. Cerutti, G. Chanfreau, C.-L. Chen, V. Cognat, M. T. Croft, R. Dent, S. Dutcher, E. Fernández, H. Fukuzawa, D. González-Ballester, D. González-Halphen, A. Hallmann, M. Hanikenne, M. Hippler, W. Inwood, K. Jabbari, M. Kalanon, R. Kuras, P. A. Lefebvre, S. D. Lemaire, A. V. Lobanov, M. Lohr, A. Manuell, I. Meier, L. Mets, M. Mittag, T. Mittelmeier, J. V. Moroney, J. Moseley, C. Napoli, A. M. Nedelcu, K. Niyogi, S. V. Novoselov, I. T. Paulsen, G. Pazour, S. Purton, J.-P. Ral, D. M. Riaño-Pachón, W. Riekhof, L. Rymarquis, M. Schroda, D. Stern, J. Umen, R. Willows, N. Wilson, S. L. Zimmer, J. Allmer, J. Balk, K. Bisova, C.-J. Chen, M. Elias, K. Gendler, C. Hauser, M. R. Lamb, H. Ledford, J. C. Long, J. Minagawa, M. D. Page, J. Pan, W. Pootakham, S. Roje, A. Rose, E. Stahlberg, A. M. Terauchi, P. Yang, S. Ball, C. Bowler, C. L. Dieckmann, V. N. Gladyshev, P. Green, R. Jorgensen, S. Mayfield, B. Mueller-Roeber, S. Rajamani, R. T. Sayre, P. Brokstein, I. Dubchak, D. Goodstein, L. Hornick, Y. W. Huang, J. Jhaveri, Y. Luo, D. Martínez, W. C. A. Ngau, B. Otillar, A. Poliakov, A. Porter, L. Szajkowski, G. Werner, K. Zhou, I. V. Grigoriev, D. S. Rokhsar, A. R. Grossman, The *Chlamydomonas* genome reveals the evolution of key animal and plant functions. *Science* **318**, 245-250 (2007). doi:10.1126/science.1143609

59. R. S. Harris, Improved pairwise alignment of genomic DNA. PhD thesis, The Pennsylvania State University (2007).

78. M. D. Guiry, G. M. Guiry, AlgaeBase. World-wide Electronic Publication. Galway: National University of Ireland. searched on 01 February 2026

116. K. P. Burnham, D. R. Anderson, Model selection and multimodel inference: A practical information-theoretic approach (2nd ed.). Springer-Verlag, New York. . (2002).

117. S. J. Phillips, R. P. Anderson, M. Dudík, R. E. Schapire, M. E. Blair, Opening the black box: an open-source release of Maxent. *Ecography* **40**, 887-893 (2017). doi:10.1111/ecog.03049

118. C. A. Maggs, C. M. Pueschel, Morphology and development *of Ahnfeltia plicata* (Rhodophyta): proposal of Ahnfeltiales ord. nov.[1]. *J. Phycol.* **25**, 333-351 (1989). doi:10.1111/j.1529-8817.1989.tb00131.x

119. J. Mclachlan, C. J. Bird, Geographical and experimental assessment of the distribution of *Gracilaria* species (Rhodophyta: Gigartinales) in relation to temperature. *Helgoländer Meeresuntersuchungen* **38**, 319-334 (1984). doi:10.1007/BF02027684

120. W. J. Rayment, Ahnfeltia plicata: A red seaweed. In H. Tyler-Walters & K. Hiscock (Eds.), Marine life Information network: biology and sensitivity key information reviews [online]. Plymouth: marine biological association of the United Kingdom. (2004).
121. C. Liu, P. M. Berry, T. P. Dawson, R. G. Pearson, Selecting thresholds of occurrence in the prediction of species distributions. *Ecography* **28**, 385-393 (2005). doi:10.1111/j.0906-7590.2005.03957.x
122. S. J. Phillips, R. P. Anderson, R. E. Schapire, Maximum entropy modeling of species geographic distributions. *Ecol. Model.* **190**, 231-259 (2006). doi:10.1016/j.ecolmodel.2005.03.026
123. R. J. Hijmans, terra: Spatial data analysis. R package version 1.7-71. Available at CRAN (2024). doi:10.32614/CRAN.package.terra
124. X. Feng, D. S. Park, Y. Liang, R. Pandey, M. Papeş, Collinearity in ecological niche modeling: Confusions and challenges. *Ecol. Evol.* **9**, 10365-10376 (2019). doi:10.1002/ece3.5555
125. R. J. Hijmans, S. Phillips, J. Leathwick, J. Elith, dismo: Species distribution modeling. R package version 1.1-4. https://CRAN.R-project.org/package=dismo (2017).
126. J. Elith, M. Kearney, S. Phillips, The art of modelling range-shifting species. *Methods Ecol. Evol.* **1**, 330-342 (2010). doi:10.1111/j.2041-210X.2010.00036.x
127. C. S. Lobban, P. J. Harrison, Seaweed Ecology and Physiology. Cambridge University Press, Cambridge, 384 p. (1994).
128. E. A. Hadly, P. A. Spaeth, C. Li, Niche conservatism above the species level. *Proc. Natl. Acad. Sci. U. S. A.* **106**, 19707-19714 (2009). doi:10.1073/pnas.0901648106
129. J. R. Hendricks, E. E. Saupe, C. E. Myers, E. J. Hermsen, W. D. Allmon, The generification of the fossil record. *Paleobiology* **40**, 511-528 (2014). doi:10.1666/13076
130. A. B. Smith, W. Godsoe, F. Rodríguez-Sánchez, H.-H. Wang, D. Warren, Niche Estimation Above and Below the Species Level. *Trends Ecol. Evol.* **34**, 260-273 (2019). doi:10.1016/j.tree.2018.10.012
131. D. L. Warren, R. E. Glor, M. Turelli, Environmental niche equivalency versus conservatism: quantitative approaches to niche evolution. *Evolution* **62**, 2868-2883 (2008). doi:10.1111/j.1558-5646.2008.00482.x

146. F. Denoeud, O. Godfroy, C. Cruaud, S. Heesch, Z. Nehr, N. Tadrent, A. Couloux, L. Brillet-Guéguen, L. Delage, D. McKeown, T. Motomura, D. Sussfeld, X. Fan, L. Mazéas, N. Terrapon, J. Barrera-Redondo, R. Petroll, L. Reynes, S.-W. Choi, J. Jo, K. Uthanumallian, K. Bogaert, C. Duc, P. Ratchinski, A. Lipinska, B. Noel, E. A. Murphy, M. Lohr, A. Khatei, P. Hamon-Giraud, C. Vieira, K. Avia, S. S. Akerfors, S. Akita, Y. Badis, T. Barbeyron, A. Belcour, W. Berrabah, S. Blanquart, A. Bouguerba-Collin, T. Bringloe, R. A. Cattolico, A. Cormier, H. Cruz de Carvalho, R. Dallet, O. De Clerck, A. Debit, E. Denis, C. Destombe, E. Dinatale, S. Dittami, E. Drula, S. Faugeron, J. Got, L. Graf, A. Groisillier, M.-L. Guillemin, L. Harms, W. J. Hatchett, B. Henrissat, G. Hoarau, C. Jollivet, A. Jueterbock, E. Kayal, A. H. Knoll, K. Kogame, A. Le Bars, C. Leblanc, L. Le Gall, R. Ley, X. Liu, S. T. LoDuca, P. J. Lopez, P. Lopez, E. Manirakiza, K. Massau, S. Mauger, L. Mest, G. Michel, C. Monteiro, C. Nagasato, D. Nègre, E. Pelletier, N. Phillips, P. Potin, S. A. Rensing, E. Rousselot, S. Rousvoal, D. Schroeder, D. Scornet, A. Siegel, L. Tirichine, T. Tonon, K. Valentin, H. Verbruggen, F. Weinberger, G. Wheeler, H. Kawai, A. F. Peters, H. S. Yoon, C. Hervé, N. Ye, E. Bapteste, M. Valero, G. V. Markov, E. Corre, S. M. Coelho, P. Wincker, J.-M. Aury, J. M. Cock, Evolutionary genomics of the emergence of brown algae as key components of coastal ecosystems. *Cell* **187**, 6943-6965.e6939 (2024). doi:10.1016/j.cell.2024.10.049

147. F. Leliaert, D. A. Payo, C. F. D. Gurgel, T. Schils, S. G. A. Draisma, G. W. Saunders, M. Kamiya, A. R. Sherwood, S.-M. Lin, John M. Huisman, L. Le Gall, R. J. Anderson, John J. Bolton, L. Mattio, M. Zubia, T. Spokes, C. Vieira, C. E. Payri, E. Coppejans, S. D'Hondt, H. Verbruggen, O. De Clerck, Patterns and drivers of species

diversity in the Indo-Pacific red seaweed *Portieria*. *J. Biogeogr.* **45**, 2299-2313 (2018). doi:10.1111/jbi.13410
